# RegVelo: gene-regulatory-informed dynamics of single cells

**DOI:** 10.1101/2024.12.11.627935

**Authors:** Weixu Wang, Zhiyuan Hu, Philipp Weiler, Sarah Mayes, Marius Lange, Jingye Wang, Zhengyuan Xue, Tatjana Sauka-Spengler, Fabian J. Theis

## Abstract

RNA velocity has emerged as a popular approach for modeling cellular change along the phenotypic landscape but routinely omits regulatory interactions between genes. Conversely, methods that infer gene regulatory networks (GRNs) do not consider the dynamically changing nature of biological systems. To integrate these two currently disconnected fields, we present RegVelo, an end-to-end dynamic, interpretable, and actionable deep learning model that learns a joint model of splicing kinetics and gene regulatory relationships and allows us to perform in silico perturbation predictions. When applied to datasets of the cell cycle, human hematopoiesis, and murine pancreatic endocrinogenesis, RegVelo demonstrates superior predictive power for interactions and perturbation simulations, for example, compared to methods that focus solely on dynamics or GRN inference. To leverage RegVelo’s full potential, we studied the dynamics of zebrafish neural crest development and underlying regulatory mechanisms using our Smart-seq3 dataset and shared gene expression and chromatin accessibility measurements. Using RegVelo’s in silico perturbation predictions, validated by CRISPR/Cas9-mediated knockout and single-cell Perturb-seq, we establish transcription factor *tfec* as an early driver and *elf1* as a novel regulator of pigment cell fate and propose a gene-regulatory circuit involving *tfec* and *elf1* interactions via the toggle-switch model.

## Introduction

Single-cell assays provide novel insight into the intricate processes of cell differentiation at a high resolution^1–3^. Waddington’s landscape^4^ conceptualizes the underlying phenotypic landscape with valleys and branching ridges, symbolizing potential cell states, along which differentiation trajectories unfold. However, building a concrete model of the gene regulatory network (GRN) and dynamics of a cell along this landscape is a major challenge in computational single-cell biology. Such a model would provide mechanistic insight and enable in silico predictions of dynamic effects of gene knockouts (KOs), thereby, informing wet lab experiments to uncover novel biological findings. Existing computational methods either recover cellular trajectories from single-cell measurements to align cells along their developmental paths^5,6^ while ignoring gene regulatory relationships, or they infer GRNs but omit cell dynamics.

Pseudotime^7–11^ and RNA velocity^12–14^ are the two most widely used concepts for reconstructing cell trajectories: pseudotime orders cells along the differentiation path but requires specifying the root state of the system, routinely assumes unidirectional differentiation, and lacks kinetic information to simulate cellular dynamics; RNA velocity attempts to overcome these limitations through a bottom-up, mechanistic model that describes dynamic changes in spliced mRNA. This approach provides estimates of the instantaneous cellular state change to study state transitions^15,16^ and allows to simulate cell state changes. However, despite its power in studying cellular dynamics^13,14^, RNA velocity relies on restrictive modeling assumptions such as gene independence and constant gene transcription rates^13,14^, thereby omitting, among other processes, transcriptional regulation underlying cell differentiation.

Computational advances in single-cell RNA sequencing (scRNA-seq) have introduced numerous methods for inferring gene regulatory networks (GRNs)^17–22^, and orthogonal data views such as scATAC-seq provide epigenetic priors^23–27^. Currently, analyses leverage GRNs in three main ways: (1) network metrics quantify gene importance^25,26^, (2) activities of modules help study cellular identity through more refined cell state clusters^17,23^ or changes along developmental paths^26^, and (3) in silico gene expression perturbations predict the corresponding effects on the state identity^22,23,25^. However, these analyses primarily rely on methods that do not learn or predict cellular dynamics or only link network components with cellular dynamics in a post-hoc fashion^26^. The single-cell community, thus, requires models that map gene regulation to cellular dynamics, to elucidate how GRNs orchestrate dynamic cellular processes.

To address this research and knowledge gap, we present RegVelo (Regulatory Velocity), a method to infer transcriptome-wide splicing dynamics coupled through gene regulation. RegVelo harnesses advancements in deep generative modeling to infer kinetic parameters and latent time by leveraging shared information across cells and genes^14^. Compared to our earlier work veloVI^14^ and previous other models^13,28–30^, we employ a prior GRN-informed neural network to couple gene dynamics and predict transcription rates based on regulator expression, thereby modeling transcription as a time- and regulation-dependent process. The resulting model is a nonlinear genome-wide dynamic differential equation parametrizable and learnable at scale in contrast to the single-gene closed-form dynamic systems traditionally employed. Our model provides a continuous velocity vector field, assesses the uncertainty of cellular state change along differentiation processes, and facilitates regulon or regulation-wise network perturbation simulation to associate cell fate decisions with gene regulatory mechanisms.

When applied to simulated data and cell cycle datasets, RegVelo’s velocity and latent time inference outperforms competing methods that employ less faithful models of transcription dynamics^28–31^; on datasets of human hematopoiesis and mouse pancreatic endocrinogenesis, RegVelo’s velocity inference entailed more consistent fate mapping and identification of putative driver genes compared to scVelo^13^ and veloVI^14^. By combining our inference paradigm with CellRank 2’s model-agnostic framework^16^, we correctly predicted the effects of network perturbations on cell fate decisions and identified gene regulatory circuitry underlying lineage-specific events. Finally, by applying RegVelo to zebrafish neural crest formation, we fully recapitulated and faithfully predicted cell fate priming and key lineage drivers. Our model identified *tfec* as an early key transcriptional regulator controlling pigment cell formation and discovered a previously unknown pigment lineage driver, *elf1*, via in silico perturbation. We validated these predictions using in vivo perturbation experiments, demonstrating excellent accordance and significant improvements over existing perturbation prediction models.

## Results

### Learning gene regulation informed cellular dynamics

RegVelo is a Bayesian deep generative model to describe cellular dynamics that takes unspliced and spliced abundances as input and outputs a cell and gene-specific latent time, the posterior distribution of velocities, and a gene regulatory network (GRN) representing transcriptional regulation. RegVelo finds these estimates by modeling unspliced and spliced RNA readouts for each gene in each cell using kinetic and neural network parameters describing dynamic transcription, splicing, degradation, latent time, and a GRN. Specifically, our model takes into account that upstream regulators control the transcription of target genes^32^, an important aspect of biological processes ignored by existing models. GRNs traditionally describe these regulatory mechanisms, allowing us to model non-constant, GRN-informed transcription rates. Combined with the remaining rates and GRN, we thereby model the developmental landscape and align cells along it.

RegVelo represents the GRN as a weight matrix, parametrized by a shallow neural network to estimate transcription rates for each gene in each cell. Each entry of the GRN matrix describes the effect of a regulator on the transcription of its targets: positive values describe activation, negative values repression, and zero entries indicate the absence of a regulatory relationship. During model training, we update the GRN parameters to infer the transcriptional dynamics of each gene in a data-driven fashion. At the same time, we constrain the GRN weights by prior knowledge of gene regulation using a penalty function; RegVelo allows prior information on gene regulation curated from single-cell multi-omics experiments^33^ or public gene regulation databases^34^. In general, GRN inference often faces two limitations: erroneous a priori estimated GRNs and the need to account for sparsity. Initial GRN estimates are imperfect due to the inference from noisy data or incomplete understanding of the regulatory mechanisms of biological systems, and sparsity is a common assumption in GRN inference^25,35^ based on studies of real biological systems^36^. To address these challenges, RegVelo improves initial GRN estimates by learning new potential regulatory interactions and incorporates priors in the parameter estimation that impose sparsity constraints either through a sparse prior network or L1-regularizing the Jacobian matrix of the dynamical system to impose smoothness of the dynamics (Methods).

RegVelo incorporates cellular dynamic estimates by first encoding unspliced (μ) and spliced RNA (*s*) readouts into posterior parameters of a low dimensional latent variable - the *cell representation* with a neural network. An additional neural network takes samples of this *cell representation* as input to parameterize gene-wise latent time as in our previous model veloVI. We then model splicing dynamics with ordinary differential equations (ODEs) specified by a base transcription and GRN weight matrix *W*, describing transcription and inferred by a shallow neural network, constant splicing and degradation rate parameters *β* and *γ*, respectively, and estimated cell and gene-specific latent times (Figure 1a; Methods). Importantly, existing methods for inferring RNA velocity consider a set of decoupled one-dimensional ODEs for which analytic solutions exist, but RegVelo relies on the single, high-dimensional ODE

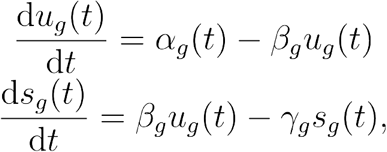

that is now coupled through gene regulation-informed transcription

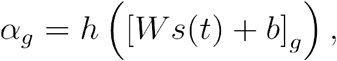

where *g* indicates the gene and *h* is a non-linear activation function (Methods). We predict the gene and cell-specific spliced and unspliced abundances using a parallelizable ODE solver^37^, as this new system does not pose an analytic solution anymore; compared to previous approaches, we solve all gene dynamics at once instead of sequentially for each gene independently of all others. The forward simulation of the ODE solver allows for computing the likelihood function encompassing all neural network and kinetic parameters (Figure 1a). We assume that the predicted spliced and unspliced abundances are the expected value of the Gaussian likelihood of the observed dataset and use gradient-based optimization to update all parameters (Methods). After optimization, we define cell-gene-specific velocities as splicing velocities based on the estimated splicing and degradation rates and predicted spliced and unspliced abundance. Overall, RegVelo allows sampling predicted readouts and velocities from the learned posterior distribution.

**Figure 1.**
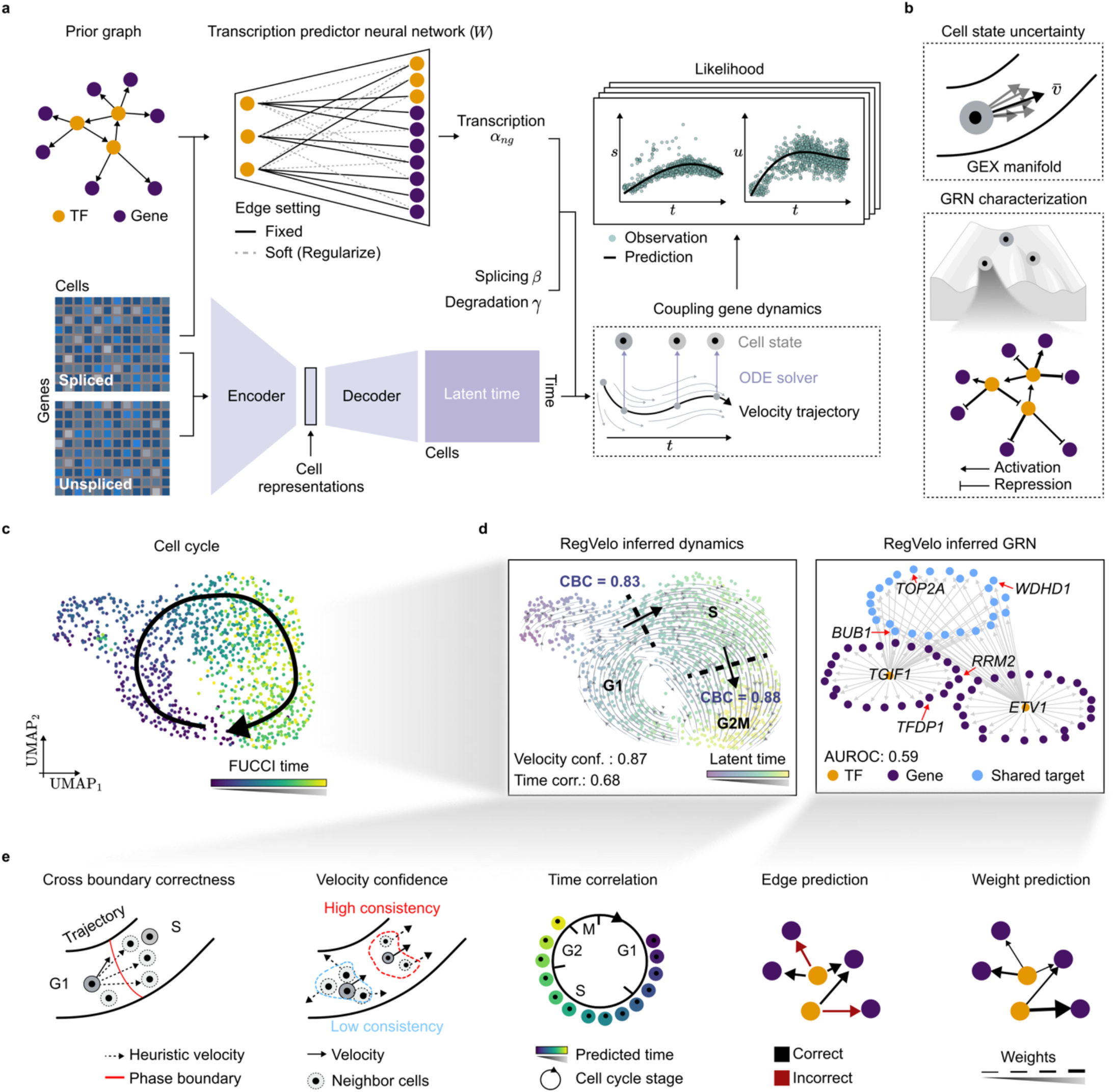
RegVelo models dynamics of cells and their underlying gene regulatory networks (GRNs) jointly. **a**. RegVelo encodes the unspliced and spliced abundance of scRNA-seq data into the cell representation through a neural network and feeds the cell representation into a decoder neural network, which outputs a cell-gene-specific latent time. Simultaneously, a prior gene regulatory graph curated from orthogonal data sources like ATAC-seq, multiome measurements, or databases guides a transcription predictor neural network. During training, this network refines and learns gene regulatory relationships, encoding each cell’s (*n*) and gene’s (*g*) transcription rate (α_*ng*_). Combined with the additionally estimated splicing and degradation rates *β* and *γ*, respectively, the model defines coupled gene dynamics and employs a parallel, high-dimensional ODE solver^37^ to estimate and predict dynamic gene expression. The forward simulation allows for defining a likelihood function encompassing all neural network and kinetic parameters, optimized in an end-to-end fashion using stochastic variational inference^38^. **b**. As a Bayesian generative model, the variational inference formulation quantifies the uncertainty of velocity estimation^14^ (top), and coupling gene dynamics with gene regulation allows us to characterize the underlying GRN of the phenotypic space (bottom). **c**. UMAP representation of a cell cycle dataset of 1146 U2OS-FUCCI cells^39^, colored by the FUCCI-derived cell-cycle score; the black arrow indicates the ground-truth transition direction. **d**. The UMAP representation colored by the RegVelo-inferred latent time and overlaid with the corresponding velocity stream and inference metrics (left; CBC: cross-boundary correctness; conf.: confidence; corr: correlation). We indicate the top two regulators (orange dots) and cell cycle-related genes (red arrows) in the fitted GRN (right). **e**. Conceptualization of five metrics to evaluate RegVelo’s inferred velocities, latent time, and GRN: the CBC quantifies how accurately inferred velocities recapitulate known cell state transitions, velocity consistency evaluates the coherence of velocities within neighboring cells, the latent time analysis assesses how well inferred latent times align with the ground truth, and comparing predicted gene regulatory edges to know relationships evaluates the GRN inference (TF: transcriptional factors; GEX: gene expression; Methods).

RegVelo outputs parameter estimates including the inferred GRN, latent time, and the posterior velocity distribution, allowing us to quantify intrinsic and extrinsic cell uncertainty (Figure 1b): intrinsic uncertainty captures uncertainty of the cell state change along the phenotypic manifold approximated by the first-order direction, while extrinsic uncertainty reflects uncertainty in future cell states^14^. RegVelo generalizes our previous model veloVI by explicitly incorporating a GRN, thereby offering a more sophisticated and realistic model of Waddington’s epigenetic landscape (Figure 1b). As such, RegVelo models the dynamics of a system of gene regulatory interactions that impose constraints and drive cell development, rather than merely modeling gene splicing^13^ or mapping cells across different time points via expression similarity^40^. Moreover, using different gene regulatory systems, RegVelo can generate velocity samples from the respective posterior distributions. By comparing velocity vector fields of perturbed regulatory systems, RegVelo also directly links gene regulation to cellular dynamics.

### Adding regulatory constraints enhances inference of cell cycle dynamics

Inference of transcriptional dynamics is a challenging task and difficult to evaluate as ground truth kinetic rates and state changes usually do not exist. To assess RegVelo’s ability to correctly infer cellular dynamics and gene regulation, nonetheless, we benchmarked it across a range of simulated and real-world datasets. First, we benchmarked our model on simulated data for which ground truth rates and latent time exist and found that RegVelo consistently outperformed competing methods (Supplementary Notes 1 and 2).

Next, we turned to a cell cycle dataset as a real-world setting where we could robustly approximate the correct differentiation history and direction from fluorescence markers.

Fluorescent ubiquitination-based cell-cycle indicator (FUCCI) U2OS cells undergo clear unidirectional transitions during the cell cycle, moving from the G1 to S to G2M phase, and yield a protein-derived cell-cycle score as a proximal measure of ground truth time (Figure 1c)^39^. After confirming that RegVelo recovered the underlying dynamics faithfully, we investigated the inferred GRN next (Figure 1d; Methods). Importantly, we did not provide a prior-informed GRN as an initial estimate to RegVelo but used a fully connected network where each gene interacts with every other gene, instead - a choice that allowed us to quantify RegVelo’s ability to infer gene regulation in an unbiased manner. To preprocess nascent and mature mRNA, we followed our previous approaches for RNA velocity inference and provided our generative model with gene expression smoothed by first order over cellular neighborhoods (Methods)

We verified that RegVelo accurately recovered the overall direction of the cell cycle by comparing the extrapolated future states of cells with prior knowledge on state transitions. Additionally, we confirmed that the inferred velocity field was consistent, which is desirable given the unidirectional nature of the underlying process. Specifically, we relied on the cross-boundary correctness (CBC) score^16,28^ that quantifies how faithfully a vector field aligns with known state transitions, *i*.*e*., G1 to S, and S to G2M phase in our scenario (Figure 1d,e and Supplementary Fig. 1a). Indeed, RegVelo achieved a comparatively high score (CBC=0.864 out of 1), high velocity consistency (0.873 out of 1), and its estimated latent time correlated highly with the FUCCI score used as ground truth (Spearman correlation = 0.683; possible values between −1 and 1). Turning to the RegVelo-inferred GRN, we focused on the most central part of the estimated network, identifying the main regulators TGIF1 and ETV1 among the most connected factors, and their top targets included cell cycle progression-related genes like *BUB1*^*41*^, *TFDP1*^*42*^, and *TOP2A*^*43*^ (Figure 1d; Methods). To consistently verify RegVelo’s ability to identify biologically correct regulons from the transcriptome data, we first curated a list of putative targets of cell cycle-related TFs^44^ from the ChIP-Atlas database^45^ as ground truth; the ChIP-Atlas integrates ChIP-seq, ATAC-seq and Bisulfite-seq experiments at large scale to offer epigenetic insight. Our method yielded highly concordant results, quantified by edge prediction performance and the correlation between the predicted regulation coefficient and the ChIP-Atlas binding score (Figure 1d, Supplementary Fig. 1b; Methods).

Existing methods for inferring splicing dynamics do not consider regulatory information, and approaches that estimate GRNs ignore cellular dynamics. To assess how combining the two aspects with RegVelo compares to methods focussing on a single facet, we applied our previous inference schemes veloVI and scVelo’s^13^ *EM* model, and the GRN inference models CellOracle^25^, GRNBoost^46^, and Pearson correlation. Whereas veloVI and scVelo assume constant rates, recent extensions like UniTVelo^28^, VeloVAE^29^, TFvelo^31^, and cell2fate^30^ try to model transcription as a time-dependent process. We, thus, also benchmarked against these velocity methods. Overall, RegVelo outperformed the other velocity methods in terms of CBC and latent time inference; among the methods that inferred the correct direction, RegVelo also had the highest velocity consistency score, showing that its velocity vector field aligned best with the ground-truth direction of the cell cycle (Supplementary Fig. 1a). To further assess whether proximal cells on the phenotypic manifold have similar latent times, we calculated the local consistency of inferred latent time for RegVelo compared to veloVI. RegVelo showed significantly better local latent time consistency than veloVI (two-sided Welch’s *t*-test, *P* < 1 × 10^−10^)), indicating that RegVelo gives smoother latent time estimates (Supplementary Fig. 1c). In terms of GRN inference, RegVelo outperformed competing approaches both on edge and regulation coefficient prediction (Supplementary Fig. 1b).

To confirm that the observed improvement of RegVelo over other methods is indeed related to incorporating gene regulation, we tested to what extent alterations to the estimated network influenced the inferred dynamics. To assess the influence of the GRN in general, we predicted splicing dynamics without any regulatory relationships present, and to mimic spurious gene-gene interactions, we randomly reconnected genes in RegVelo’s network (Methods). As expected, both scenarios led to a lower CBC score and latent time correlation (Supplementary Fig. 1d), highlighting that incorporating gene regulation improves cellular dynamic inference.

These results on both simulation and real data demonstrate RegVelo’s ability to correctly and consistently infer cellular dynamics and gene regulation more faithfully than competing methods.

### Regulatory-constrained dynamic inference accurately recovers lineage formation in pancreatic endocrinogenesis

RegVelo is a generative model that couples cellular dynamics with regulatory networks. We can, thus, perform in silico counterfactual inference to test the cellular response upon unseen perturbations of a TF in the regulatory network: for a trained RegVelo model, we ignore regulatory effects of the TF by removing all its downstream targets from the GRN, *i*.*e*., depleting the regulon, and generate the perturbed velocity vector field (Figure 2a; Methods). The dissimilarity between the original and perturbed cell velocities - the perturbation effect score - reflects the local changes on each cell induced by perturbations; we quantify this score with cosine dissimilarity (Methods).

**Figure 2.**
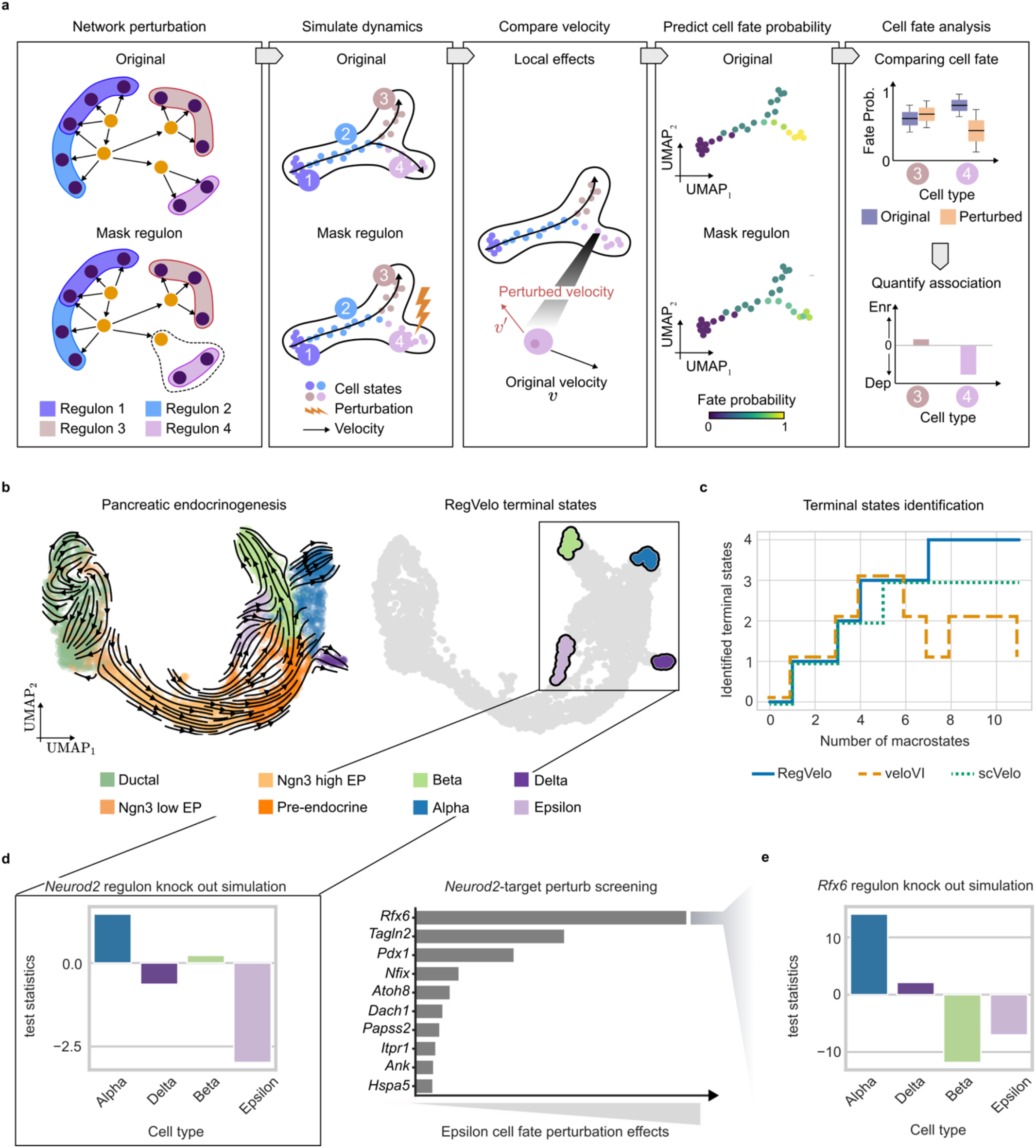
Perturbation simulation quantifies genetic regulation effects on cell fate decisions in pancreatic endocrinogenesis. **a**. As an inference model that incorporates gene regulatory mechanisms, RegVelo can simulate the dynamics resulting from in silico perturbations by masking regulons to mute the functions of the corresponding TFs. Comparing velocity differences for each cell quantifies local perturbation effects, and passing the original and perturbed velocity estimations into the typical CellRank workflow^15,16^ quantifies cellular fates, where changes in cell fates identify perturbation effects (Methods). Box plots are schematics indicating the median (center line), interquartile range (hinges), and whiskers at 1.5x interquartile range. **b**. UMAP embedding of 3696 pancreatic endocrinogenesis cells, colored by cell type according to the original study and the RegVelo-inferred velocity stream (left), and the terminal states predicted by CellRank using RegVelo’s velocity field (right). **c**. The number of correctly identified terminal states compared to the number of specified macrostates indicates how faithfully a method recovers known biology with CellRank; different colors and line styles indicate different methods. **d**. Cell fate perturbation effects estimated from *Neurod2* regulon perturbation simulation (left), and the corresponding ten *Neurod2*-targets affected most in the Epsilon lineage. **e**. Cell fate perturbation effects estimated from *Rfx6* regulon perturbation simulation.

RNA velocity describes a high dimensional vector field representing cellular change along the phenotypic manifold but lacks interpretability and quantifiable measures of the long-term cell behavior. We recently proposed CellRank^15,16^ to bridge this gap by leveraging gene expression and an estimated vector field to model cell state transitions through Markov chains and infer terminal cell states. For each terminal state identified, CellRank calculates the probability of a cell transitioning to this state - the fate probability - that allows us to predict the cell’s future state. By combining RegVelo’s generative model with CellRank, we connect gene regulation with both local cell dynamics and long-term cell fate decisions, and how they change upon in silico perturbations. In the context of our perturbation analyses, we compare CellRank’s prediction of cell fate probabilities for the original and perturbed vector fields, to find enrichment (increased cell fate probability) or depletion (decreased cell fate probability) effects towards terminal states (Figure 2a).

To highlight how our model elucidates cellular dynamics by incorporating gene regulation, we applied RegVelo to a pancreatic endocrine development scRNA-seq dataset^47^. To guide RegVelo’s inference of the underlying cellular dynamics, we estimated a prior GRN by scanning for motifs within candidate cis-regulatory elements (CREs) and integrating gene expression and peak accessibility information with Pando^24^ to define the prior GRN estimate (Methods). For this GRN construction, we relied on a multiome dataset of paired ATAC and gene expression of the same system at embryonic days 14.5 and 15.5^48^.

Passing RegVelo’s inferred velocity field to CellRank, we delineated the cycling population of ductal cells and predicted all four terminal states, namely Alpha, Beta, Delta, and Epsilon cells^47^ (Figure 2b,c; Methods). We specifically explored how these lineages form, predicting that some Epsilon cells become progenitors of Alpha cells, which is consistent with current reports^47–49^ (Supplementary Fig. 2a). We identified two states based on fate probabilities that divide the Epsilon cell population: state A expressed Alpha cell-specific genes highly (*Pou6f2*^*50*^, *Irx1*^*51*^, and *Smarca1*^*52*^), and state B *Neurog3*, consistent with Neurogenin3 activity-dependent Epsilon cell genesis^53^ (Supplementary Fig. 2b; Methods).^53^ Similar to our analyses of the cell cycle data, we deleted or shuffled the learned GRN to generate new vector fields to verify the utility of combining transcriptional dynamics with gene regulation; both alterations led to the loss of identified terminal states (Supplementary Fig. 2c). An additional indication for the GRN’s importance is that our previous methods veloVI and scVelo achieved a lower terminal state identification (TSI) score^16^ that quantifies how accurately a method identifies terminal states (Supplementary Fig. 2d; Methods).

### RegVelo disentangles ductal and epsilon cell dynamics in pancreatic endocrinogenesis through in silico perturbations

Following our analysis of cell maturation, we investigated how to interpret the role of gene regulation in the cycling population of ductal cells with RegVelo. By correlating gene expression with a cell cycling phase score inferred from prior knowledge on cycling markers, we identified *E2f1* as the highest-ranked gene (Supplementary Fig. 3a,b; Methods). To predict the cell set related to E2f1 functions, we removed the E2f1 regulon from the GRN and used cosine dissimilarity to compare the vector fields before and after perturbation (Supplementary Fig. 3c). We thereby identified a ductal cell subpopulation exhibiting large change that overlaps mostly with the S-phase and, thus, the stage influenced most by *E2f1*^*54*^ (Fisher exact test, *P* < 1 × 10^−10^ Supplementary Fig. 3d). Importantly, this perturbation effect score does not simply reflect *E2f1* expression, as Ngn3 high EP cells express *E2f1* at a higher level than ductal cells (Supplementary Fig. 3d). Moreover, gene ontology (GO) enrichment analysis suggests that the predicted E2f1 downstream targets mainly enrich the cell cycling phase transition-related pathway (Supplementary Fig. 3e), aligning with previous findings that E2f1 is a cell cycling regulator^55^. These findings highlight that the RegVelo-predicted perturbation effects score can help elucidate and identify the role of different cell states and pathways.

We explored the *E2f1* regulon network further to investigate whether RegVelo could recover known E2f1 downstream targets that contribute to cell cycle regulation. As a first step, we validated that the inferred gene regulatory network is identifiable and robust under different runs of RegVelo (Pearson correlation > 0.95; Supplementary Fig. 3f). We then used the prior GRN as a baseline to compare the RegVelo-inferred ductal cell GRN performance when predicting E2f1 targets (Methods). Using E2f1 ChIP-seq results from the ChIP-Atlas database as ground truth, RegVelo’s GRN showed significant improvement over the prior GRN in predicting E2f1 targets (two-sided Welch’s *t*-test, *P* < 1 × 10^−10^; Supplementary Fig. 3g). Focusing on the top 20 predicted targets, we recovered eight cycling-related genes, including *Cdt1*^*56*^, *Uhrf1*^*57*^, *Stmn1*^*58*^, *Id2*^*59*^, *Mcm6*, and *Mcm7*^*60*^. The Pando-based prior GRN, however, did not include key cell cycling modulators such as *Hells* and *Cenpk* as E2f1 targets, whereas the RegVelo-inferred GRN correctly predicted them among the top downstream effectors (Supplementary Fig. 3h). Recent experimental results validate these connections^61,62^. In summary, RegVelo accurately infers cellular dynamics and gene regulation, enabling the prediction of perturbation effects.

Next, we used RegVelo to understand the perturbation effects of TFs on cell fate decisions. This analysis required us to ensure that RegVelo gives stable cell fate probability prediction to avoid stochastic effects caused by random parameter initialization, for example. To verify robustness, we measured the correlation of velocities or cell fate probabilities between pairs of estimates from different runs of the model (Methods). RegVelo scored a high mean correlation and low variance, implying that RegVelo has high identifiability on cell fate prediction (Supplementary Fig. 4a). Next, we quantified the perturbation effect of each TF regulon and validated its stability over multiple model fits for each lineage (Supplementary Fig. 4b; Methods). Having verified RegVelo’s robust inference, we reasoned that the TFs corresponding to the biggest depletion effects after in silico perturbation pose putative drivers of the specific lineage. Using this perturbation simulation-based ranking of putative lineage drivers, RegVelo accurately predicted known drivers in the four lineages, including Smarca1^52^, Arx^63^ in the Alpha, Pdx1^64^, Mnx1^65^ in the Beta, Hhex^66^ in the Delta lineage (Supplementary Fig. 4c) and achieved an overall high driver ranking performance, evaluated via the AUROC (AUROC=0.95; Methods).

We further focused on the poorly understood process of Epsilon cell differentiation and observed that our in silico screening predicted Neurod2 as one of the putative drivers of Epsilon cell differentiation (Supplementary Fig. 4c). Previous studies have found that pancreatic cells transiently expressed *Neurod2* during pancreatic endocrinogenesis^67^ and suggested the role of *Neurod2* in Epsilon cell conformation^48^. To uncover the potential mechanisms of *Neurod2* in orchestrating Epsilon cell differentiation, we employed RegVelo’s generic circuit screening feature to perturb each downstream target of Neurod2 and predict the outcomes of these perturbations on Epsilon differentiation. According to these results, perturbation of the Neurod2-*Rfx6* regulation contributes the most to the maturation of Epsilon cells (Figure 2d), an observation consistent with recent studies highlighting the importance of this regulation^48^. Perturbing the Rfx6 regulon suggests its involvement in Beta and Epsilon cell differentiation, an observation supported by previous experimental studies^68^ (Figure 2e).

### RegVelo recovers regulatory motifs governing cell fate decisions in hematopoietic differentiation

While previous RNA velocity models successfully recovered the general mechanisms of pancreatic endocrine development, the same methods did not provide insight into gene regulation, could not predict perturbation effects, and failed altogether when applied to hematopoietic datasets. The erroneous inference of cell dynamics primarily arises from the assumption of constant transcription rates, which does not hold true in the dynamic regulation processes during hematopoiesis^16,69,70^. As RegVelo does not model transcription as constant, we hypothesized that our proposed model would not face the same limitation and, additionally, inform on gene regulatory processes during hematopoiesis. Further, hematopoiesis is a well-studied system, offering an opportunity to further validate our TF perturbation predictions based on the literature.

To apply RegVelo to a hematopoiesis dataset including five distinct lineages^22^ (Figure 3a), we first curated a prior GRN, relying on two orthogonal resources to mitigate the effect of incompleteness: as a first estimate, we used a publicly available GRN inferred from single-nucleus ATAC and gene expression measurements of bone marrow mononuclear cells with *Dictys*^*26*^. We then extended this representation with an additional GRN curated from human hematopoiesis ChIP-seq datasets from the UniBind database^71^ to obtain our final prior GRN. After feeding this GRN into our standard RegVelo workflow and passing the estimated vector field to CellRank, the resulting fate probabilities correctly described the known lineage commitment and hierarchy of hematopoiesis and allowed us to recover all five terminal states and recapitulate known state transitions faithfully (Figure 3b, Supplementary Fig. 5a; Methods). We also observed a change in RegVelo’s intrinsic estimation uncertainty from high in progenitor cells (MEP and GMP) to low in terminal states (Figure 3c), consistent with the fact that mature cells do not change their identity. Having validated RegVelo’s ability to correctly model cellular change along the phenotypic manifold, we analyzed in silico perturbation predictions and the recovered GRN, next.

**Figure 3.**
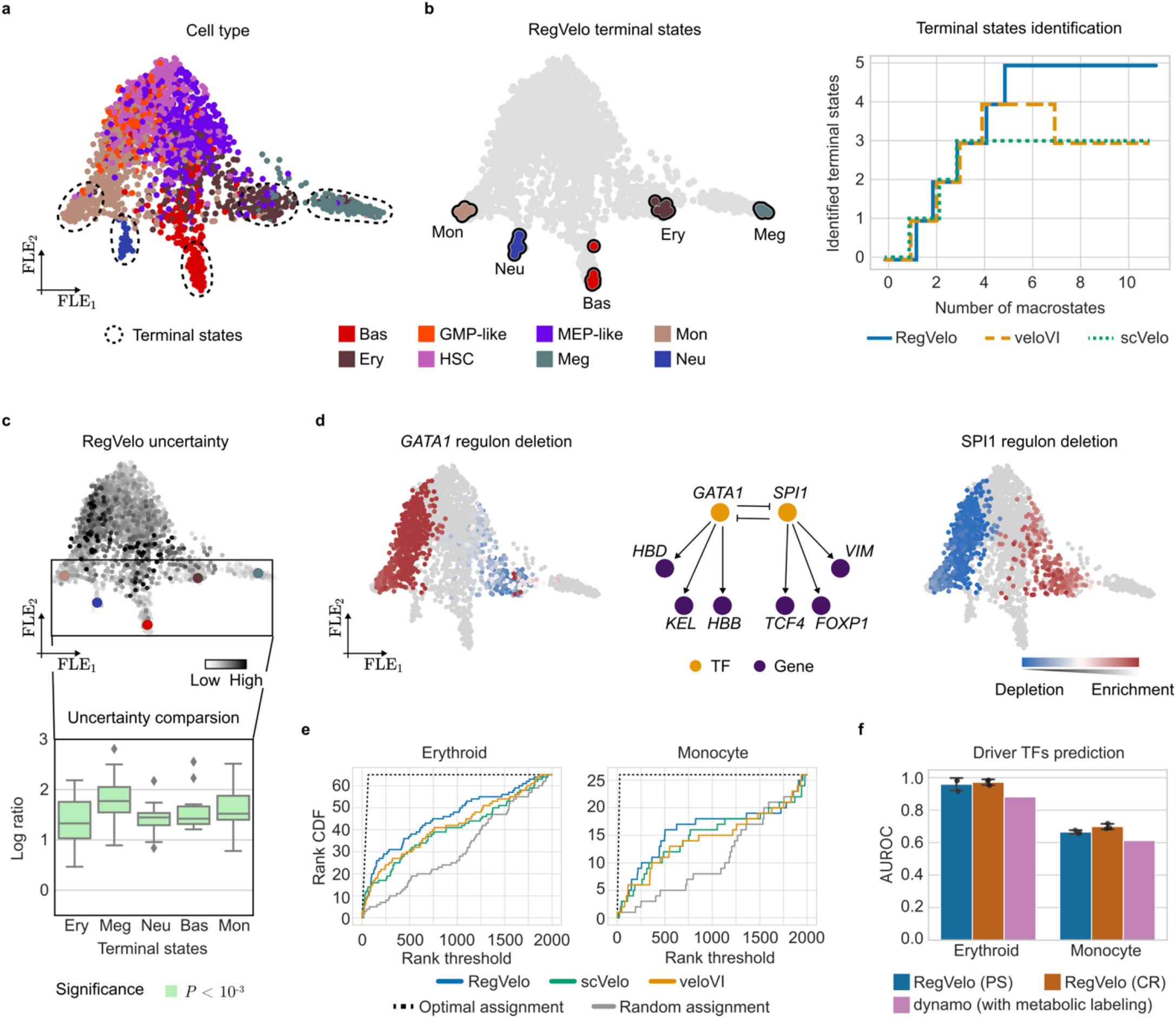
Inferring regulatory dynamics during human hematopoiesis differentiation. **a**. Force-directed layout embedding (FLE) of 1947 human hematopoietic stem and progenitor cells (HSPCs), colored according to original cell type annotation (HSC: hematopoietic stem cell; Meg: megakaryocyte; Ery: erythrocyte; Bas: basophil; Neu: neutrophil; Mon: monocyte; MEP-like: megakaryocyte and erythrocyte progenitor-like; GMP-like: granulocyte and monocyte progenitor-like). **b**. Combining the RegVelo state change estimates with CellRank to infer terminal states recovers all five terminal states (left). The number of correctly identified terminal states compared to the number of specified macrostates highlights that RegVelo performs optimally and outperforms competing methods (right; Methods) **c**. FLE of HSPCs colored by RegVelo-inferred intrinsic uncertainty and terminal states (top; Methods), and its uncertainty compared to veloVI in each terminal state based on a log-transformed ratio (bottom; Methods). Box plots indicate the median (center line), interquartile range (hinges), and 1.5x interquartile range (whiskers) (*N* = 30 cells). **d**: FLE embedding colored by RegVelo’s cell-specific perturbation effect in the erythroid and monocyte lineage after deleting the GATA1 (left) and SPI1 (right) regulon, and the core regulatory subnetwork surrounding the predicted toggle-switch between GATA1 and SPI1 (middle). **e**. Ranking of known lineage-associated genes for the erythroid and monocyte lineages using different methods with the CellRank workflow (Methods). **f**. AUROC of predicting lineage drivers for erythroid and monocyte cells (PS: perturbation simulation; CR: CellRank; *N* = 3 trained models each).

Deletion of the GATA1 regulon, a well-known master regulator of the megakaryocyte-erythroid progenitor (MEP) lineage^72^, results in RegVelo-predicted depletion effects in erythrocytes and enrichment effects in monocytes. Similarly, deleting the master regulator of the granulocyte-monocyte progenitor (GMP) lineage^73^, the SPI1 regulon, led to enrichment effects in the erythrocyte branch. RegVelo’s GRN also included the toggle switch motif of SPI1 and GATA1, consistent with previous experimental studies^74,75^. At the same time, RegVelo accurately predicted GATA1 downstream effectors directly influencing the hemoglobin gene expression, aligning with previous experimental findings on the *GATA1* downstream regulatory network^76^ (Figure 3d).

To show that RegVelo provides insight beyond the capability of competing methods, we benchmarked the inferred cellular dynamics compared to veloVI and scVelo (Methods). Following CellRank 2’s workflow^16^, we used the log2 ratio of the CBC scores as a quantitative metric for evaluating velocity inference and observed that RegVelo outperformed scVelo in 5 out of 6 cell state transitions. Although veloVI shows an overall comparable CBC score of forward cell state transition to RegVelo, it results in lower CBC scores of backward cell state transition, indicating an incorrect backflow in the veloVI vector field (Supplementary Fig. 5b,c; Methods). To assess the effect of model regularization, we compared the results of unregularized and J-L1 regularized RegVelo estimates. The regularization led to improved cell state transition predictions, indicating that increased model sparsity improves dynamic inference, consistent with previous studies on NeuralODE models^77^.

Using cell-cell transition probabilities computed with CellRank to infer terminal states, RegVelo was the only method to recover all five terminal states (TSI = 0.95), and it significantly outperformed the other methods (one-sided Welch’s *t*-test: RegVelo vs. veloVI, *P* = 1.59 × 10^−5^; RegVelo vs. scVelo: *P* = 2.23 × 10^−5^; Figure 3a,b, Supplementary Fig. 6a; Methods). As an extension of veloVI, we specifically compared RegVelo inference results to veloVI’s predictions: first, we examined how robust each method’s velocity or latent time estimations were under different runs. Overall, RegVelo’s estimates were more stable and associated with a higher consistency compared to veloVI (Supplementary Fig. 6b). Next, compared to RegVelo’s dynamically decreasing intrinsic estimation uncertainty, veloVI’s inference resulted in high intrinsic uncertainty in every cell type (Supplementary Fig. 6c). Finally, the predicted cell fate probabilities of veloVI and scVelo violated known transitions (Supplementary Fig. 6d).

Estimating fate probabilities with CellRank based on RegVelo’s vector field allows us to identify putative drivers by correlating them with gene expression^16^. We employed this strategy to investigate whether incorporating gene regulation in the splicing model leads to better identification of known genes associated with lineage priming compared to classical models that omit gene regulation (Methods). To do so, we first curated a list of known markers of the lineages to compare it to the predictions of each method. Next, we ranked genes using different velocity methods and assessed how well the ranked genes matched the sets of known markers. Here, we focused on the monocyte and erythroid lineages, as they represent two distinct branches, characterized by different markers. Compared to competing methods, using RegVelo’s velocity estimates as input for CellRank achieved the best ranking for both the monocyte and erythroid lineage (Figure 3e). Notably, either by ranking according to perturbation simulation or by using CellRank with RegVelo estimated velocities, RegVelo outperformed dynamo with respect to driver TF prediction on both lineages (Methods), even though the method has access to metabolic labeling information (Figure 3f).

The analysis of the hematopoietic system with RegVelo highlighted our model’s capability to accurately recover lineage hierarchy and regulatory connections, outperforming competing approaches including those with access to more rich data such as metabolic labels.

### RegVelo screens TF knockout effects for cell fate decisions in zebrafish

Neural crest (NC) cells contribute a broad range of derivative cell types to functionally distinct tissues and organs, including bone, cartilage and connective components of the craniofacial skeleton, the majority of the body’s pigment cells, and elements of the peripheral nervous system. Deciphering dynamic programs that control neural crest formation holds promise of understanding vertebrate evolution, developmental disorders, and neural-crest-derived cancers^78–80^. Based on our promising in silico validation experiments on the cell cycle, pancreatic endocrinogenesis, and hematopoiesis, we hypothesized that RegVelo could be more generally used to combine high-throughput single-cell transcriptomic readouts with GRN priors into an interpretable, dynamic, and actionable model of neural crest formation. To decode the complex regulatory dynamics underlying neural crest development, we employed Smart-seq3^81^ to profile 1,180 neural crest cells and their derivatives at seven time points, achieving a median coverage of 7,791 genes per cell (Figure 4a,b and Supplementary Fig. 7a,b). We then applied RegVelo with a prior GRN inferred with SCENIC+^23^ based on a matched single-cell multiome dataset^82^ (Figure 4a; Methods).

**Figure 4.**
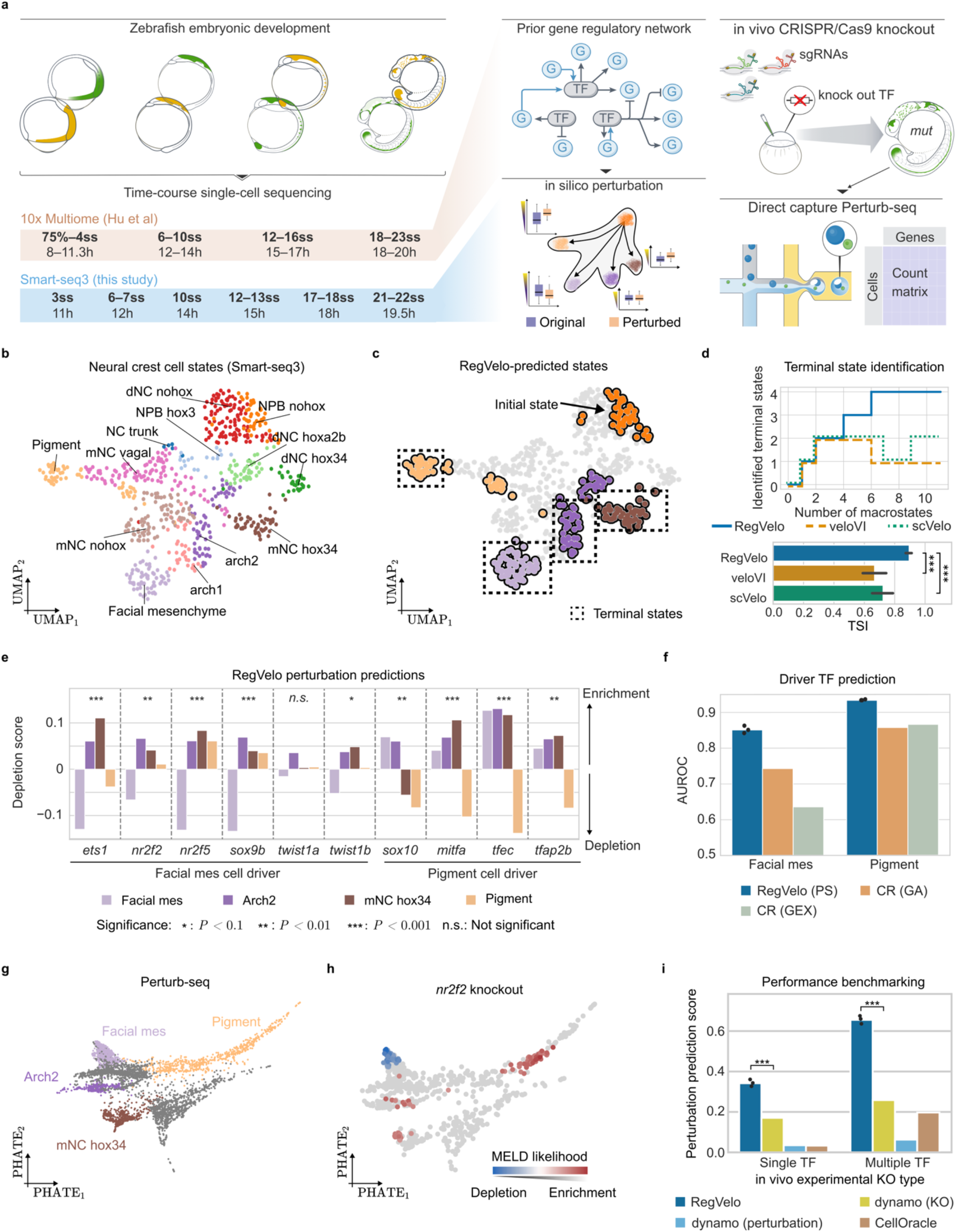
RegVelo uncovers key regulators of cell fate decisions during zebrafish neural crest development. **a**. Schematic of the experimental design for time-resolved single-cell gene expression and multimodal assays and in vivo Perturb-seq to capture neural crest cell development (ss: somite stage; h: hour post fertilization; mut: mutant). Box plots indicate the median (center line), interquartile range (hinges), and 1.5x interquartile range (whiskers); the shown box plots are schematics. **b**,**c**. UMAP embedding of 697 neural crest cells from Smart-seq3 dataset colored by cell states (b) and known initial and terminal states (c) (NPB, neural plate border; dNC, delaminating neural crest; mNC, migratory neural crest; mes, mesenchyme; arch2, second pharyngeal arch). **d**. We plot the number of identified terminal states against the number of macrostates specified (top) and the terminal state identification (TSI) score using RegVelo (blue), veloVI (orange), and scVelo (green) (*N* = 20; two-sided Welch’s *t*-test, *P* < 0.001). Error bars correspond to 95% confidence intervals. Line styles and colors indicate different methods. **e**. RegVelo-predicted cell fate perturbation effects of known lineage drivers. We used a student’s t-test to assess the significance of the perturbation effects on mNC head mesenchymal or pigment lineages. **f**. AUROC of lineage driver prediction for facial mesenchymal and pigment lineage specification based on RegVelo’s perturbation simulation (PS), or CellRank using gene expression (GEX) or gene activity (GA) of TFs. **g**. PHATE embedding of 39,453 neural crest development cells from the joint Perturb-seq datasets with terminal states colored. **h**. PHATE embedding colored by in vivo *nr2f2* perturbation score quantified by MELD, Only cells with MELD likelihood scores in the top 20% (>0.8 quantile) or bottom 20% (<0.2 quantile) are colored; others are shown in grey. (*N* = 834 cells including both control and *nr2f2* knockout) **i**. Benchmark of in silico perturbation effect prediction using RegVelo, dynamo, and CellOracle (*N* = 3 trained RegVelo models; one-sided Welch’s *t*-test, *P* = 0.0004 for single knock out and *P* = 0.00015 for multiple knockout).

As the general state transitions and differentiation order of neural crest development are known, we confirmed that RegVelo adheres to the established differentiation hierarchy before relying on RegVelo’s in silico perturbation scheme to identify lineage regulators. Indeed, RegVelo demonstrated highly accurate dynamic inference as it allowed us to correctly identify all terminal states, namely post-otic migratory neural crest (mNC hox34), non-hox facial mesenchymal, second pharyngeal arch (arch2), and pigment cells (Figure 4c,d). Additionally, the RegVelo-inferred latent time showed a positive correlation with the actual developmental stages (Supplementary Fig. 7c,d), and accurately placed the non-hox neural plate border (NPB nohox) cell state at the start of the developmental process^83^ (Figure 4c and Supplementary Fig. 7c). Taken together, these findings verify the high fidelity of RegVelo’s velocity inference compared to other methods (Figure 4d).

To systematically identify putative lineage drivers based on in silico regulon perturbation, we followed our previous analyses to quantify perturbation effects of cell differentiation towards the four terminal states (Methods). RegVelo accurately predicted the perturbation effects of known drivers of neural crest derivatives, including key regulons of the pigment (mitfa^84^, sox10^85^) and mesenchymal (sox9b^86^, nr2f2^87^, twist1b^88^) lineages (Figure 4e).

As RegVelo allows us to simulate perturbed dynamics *de novo*, we were able to identify lineage drivers expressed at the beginning of a trajectory but not in the terminal states, while correlation-based methods tended to overlook the same genes (Supplementary Fig. 7e; Methods). For instance, neural plate border and delaminating neural crest cells (early states during NC formation) highly expressed the cranial mesenchymal lineage drivers, *ets1*^*89*^, *nr2f5*^*87*^, *sox9b*^*90*^, and *twist1b*^*91*^ that were significantly downregulated in cells at the terminal states (log_2_, FC < 0, *P* < 0.001; Supplementary Fig. 7e-g). While RegVelo correctly predicted these genes as drivers of the mesenchymal lineage, a correlation-based analysis with the classical CellRank workflow and in silico perturbation approach by dynamo^22^ erroneously identified them as putative drivers of post-otic migratory neural crest development (mNC hox34) (Supplementary Fig. 8a-c). Finally, our RegVelo-based workflow ranked these genes consistently higher than the correlation-based approach (Figure 4f).

Advances in single-cell Perturb-seq have enabled precise testing of transcription factor function through targeted knockouts in living organisms^92^. Using this approach, we validated RegVelo’s predictions of TF knockout with our improved CRISPR/Cas9 and direct capture Perturb-seq workflow^80^ (Figure 4a,g): we designed two to four chemically modified single guide RNAs^93^ to target each TF (*tfec, mitfa, bhlhe40, tfeb, elf1, nr2f2*, and *nr2f5*) to maximize the knockout efficiency and computed the in vivo perturbation effects, defined as the MELD likelihood scores of lineage changes between wild type and mutant embryos, where a score lower than 0.5 indicates depletion (Methods). Regarding the known factors, we found the experimental results consistent with the computational predictions. For instance, the in vivo knockout of the top pigment driver, *mitfa*^*84*^, resulted in pigment lineage depletion (lineage-specific median MELD likelihood = 0.3354), while knocking out the cranial ectomesenchymal driver *nr2f2*^*94*^ depleted mesenchymal cells as predicted by RegVelo (lineage-specific median MELD likelihood = 0.47, Figure 4h). We next systematically examined the in silico knockout performance using the in vivo Perturb-seq data.

To evaluate the overall performance of RegVelo’s perturbation simulation, we extended our seven-TF Perturb-seq data with an additional complementary Perturb-seq dataset^80^ to define a joint panel of 14 single-TF and 8 multiple-TF knockouts and its downstream cell type depletion effects as a notion of ground truth. Next, we evaluated model performance by simulating single or multiple TF knockouts to compute the Spearman correlation between the predicted depletion score and the experimental derived depletion likelihood and compared it to dynamo^22^ and CellOracle^25^ that employ either only dynamic modeling or GRN inference (Methods). For the single-TF knockout scenario, RegVelo predictions resulted in a nearly 2-fold improvement (mean Spearman correlation of 0.35) over the other two methods (Spearman correlation < 0.17; Figure 4i), and we observed a similar trend for the multiple-TF knockout experiment (average Spearman correlation of 0.60 vs. 0.22; Figure 4i).

Overall, having established that RegVelo correctly and robustly infers the dynamic cellular change of known zebrafish neural crest development, we sought to employ our framework to identify novel regulatory relationships driving this embryonic developmental process.

### tfec regulates pigment cell development as an early driver

Previous studies have established the differentiation programs of pigment progenitor cells into iridophores, melanophores, and xanthophores at later developmental stages^95,96^. However, the initial specification of the pigment lineage within the migratory neural crest remains unresolved. We, thus, turned to the trained neural crest RegVelo model to identify putative drivers and found *bhlhe40, tfec*, and *mitfa*^*97*^ ranked as the top three genes (Figure 5a), and the known pigment lineage^98^ markers *cdkn1ca* and *atp6ap2* among the top positively regulated targets of tfec (Figure 5b,c). Additionally, other target genes of the tfec regulon, such as *lrmda* and *psen2*, were enriched in pathways of melanocyte differentiation (adjusted *P* = 0.004; Figure 5c). We, thus, hypothesized that the tfec transcription factor contributes to early pigment cell fate priming and turned to orthogonal computational and experimental approaches for validation.

**Figure 5.**
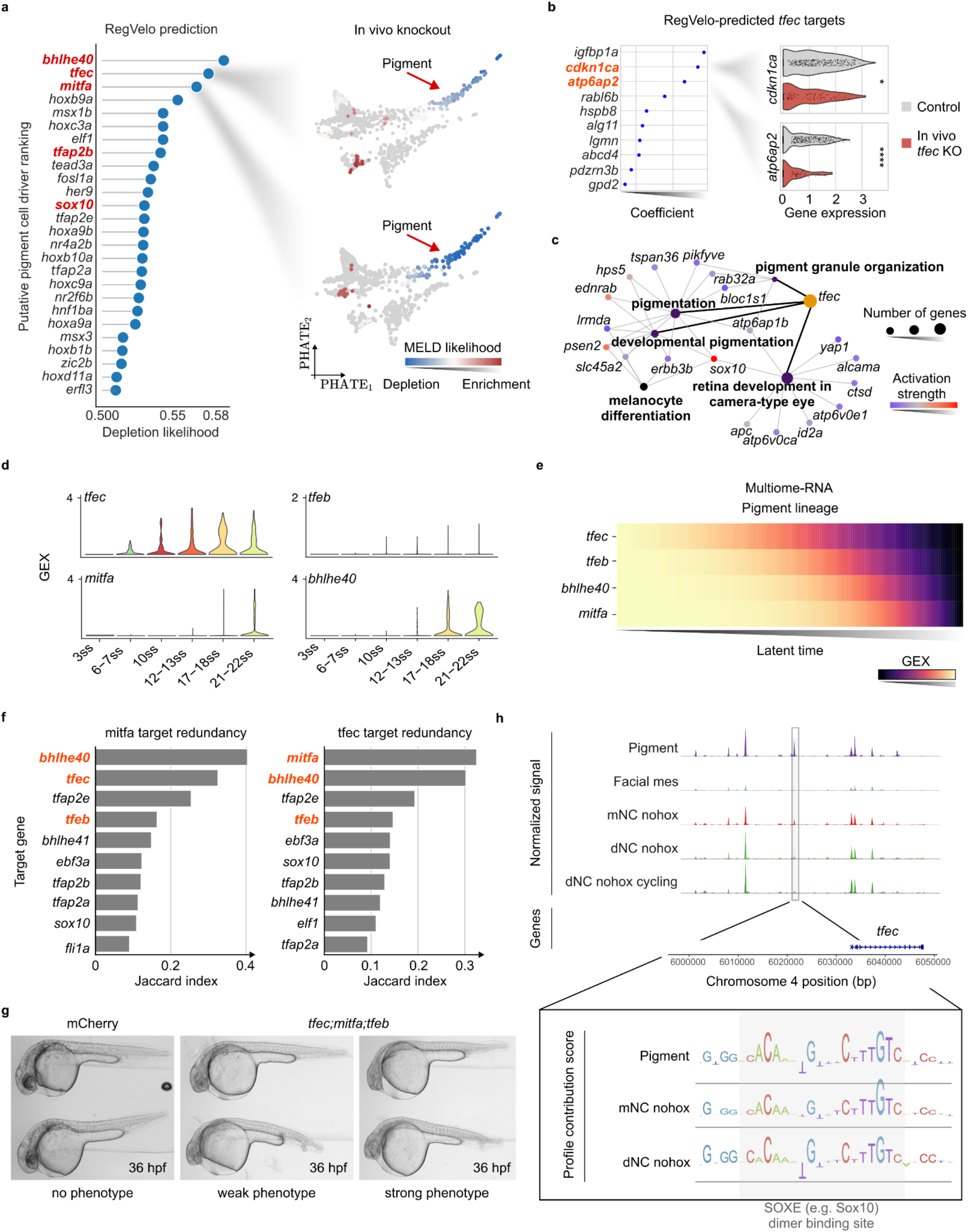
tfec regulates pigment cell development as an early driver. Identification of top putative driver TFs of the pigment lineage through RegVelo’s in silico simulation, highlighting known pigment drivers in red (left). The PHATE embeddings visualize the MELD likelihood, indicating the pigment lineage depletion in *tfec* and *mitfa* knockouts (right). The top 10 positive regulated tfect targets predicted by RegVelo and ranked by inferred weights (left). Violin plot of top target gene expression in in vivo Perturb-seq dataset (right; *N* = 779 cells; one-sided Welch’s *t*-test, *P* = 0.02 for *cdkn1ca, P* = 3.44 × 10^−8^ for *atp6ap2*). **c**.Gene ontology network analysis of RegVelo-predicted targets. **d**. Expression patterns of representative bHLH factors in the pigment lineage within our Smart-seq3 data along developmental stages. **e**. Gene expression patterns from multiome data of representative bHLH factors in the pigment lineage over latent time as computed by original study^82^. **f**. Bar plots present the regulons with the highest Jaccard index in RegVelo inferred GRN, indicating shared target genes with the mitfa(+) and tfec(+) enhancer-driven regulons and implying potential redundancy among tfec, mitfa, bhlhe40, and tfeb factors highlighted in red. **g**. Contrast images illustrate the albino phenotype in F_0_ crispants after 36 hours past fertilization (hpf), following the triple knockout of *tfec;mitfa;tfeb*, without, with weak, and with strong phenotype (Chi-square test of independence, *P* < 1 × 10^−10^ (experiment 1) and *P* < 1 × 10^−10^ (experiment 2)). **h**. The enhancer candidate upstream of *tfec* contains TF binding motifs for Sox10, indicated by the ChromBPNet-predicted contribution score. This enhancer candidate is mildly accessible in migratory neural crest cells and more accessible in the pigment lineage, indicated by the ATAC peak signals.

Single-cell transcriptomic analysis of both our Smart-seq3 dataset and a published multiome dataset^82^ identified *tfec* as the earliest expressed basic helix-loop-helix (bHLH) TF, preceding the precursor driver *mitfa* and others such as *tfeb* and *bhlhe40* (Figure 5d,e). Targets of tfec in RegVelo’s GRN and targets of the known pigment lineage drivers mitfa, tfeb, and bhlhe40 largely overlapped, implying redundant functions of drivers in pigment lineage specification (Figure 5f, Supplementary Fig. 9a). Additionally, our Perturb-seq experiment showed profound depletion effects for *tfec* knockouts compared to *mitfa* in both early and late pigment lineages (two-sided paired Wilcoxon test, *P* < 1 × 10^−12^; Supplementary Fig. 9b). In fact, only the simultaneous knockout of *mitfa, tfec*, and *tfeb* produced a significant phenotype of pigment depletion in 2-days-post-fertilization embryos compared to the single-gene knockouts^99^ (Figure 5g). We performed two experiments with no phenotype (pigmented), weak phenotype (some pigmentation), and strong phenotype (no pigmentation) embryos. Experiment 1 (experiment 2) included 35 (44) control and 0 (0) knockout embryos without a phenotype, 1 (4) control and 4 (17) knockout embryos with weak phenotype, and 0 (0) control and 43 (28) knockout embryos with a strong phenotype (Chi-square test of independence, *P* < 1 × 10^−10^ (experiment 1) and *P* < 1 × 10^−10^ (experiment 2)).

To understand the regulatory relationships of our newly identified pigment lineage driver, tfec, we investigated its upstream regulators by leveraging the multiome-ATAC data^82^. We used the base-resolution contribution scores derived from the convolutional neural network models^82^ trained and interpreted by ChromBPNet^100^ which indicated the putative TF binding instances (Methods). The master regulator of NC development, Sox10, is essential for pigment lineage specification^85^ - a fact that we accurately rediscovered through our RegVelo predictions (Figure 5a). Our ChromBPNet-based analysis identified Sox10 binding sites in the upstream *cis-*regulatory element of *tfec* and, this, further supported this fact (Figure 5h).

Overall, our experimental validations and orthogonal computational approaches support RegVelo’s prediction that *tfec* acts downstream of sox10 and implies that tfec specifies the fate of pigment cell lineages ahead of other pigment specification TFs from the bHLH family, such as mitfa, tfeb, and bhlhe40.

### RegVelo identifies an early pigment lineage driver, elf1, together with its regulatory context

In addition to traditionally recognized bHLH factors, RegVelo ranked elf1, a TF from the erythroblast transformation specific (ETS) family, among the top ten putative regulators of pigment fate decisions (Figure 5a and Figure 6a,b). We again relied on our in vivo Perturb-seq data as well as whole-mount mRNA in situ hybridization by hybrid chain reactions (HCR)^101^ to verify this predicted association. Indeed, the Perturb-seq analysis revealed a significant depletion effect in the pigment lineage upon *elf1* knockout (Figure 6b), and HCR showed decreased pigment cells in *elf1-*knockout embryos in both cranial and post-otic regions, verifying *elf1* functioning in pigment lineage specification at all axial levels (Figure 6c). Moreover, *elf1* knockouts were enriched for mesenchymal fates (facial mesenchyme, the first and second pharyngeal arches), accompanied by downregulation of *mitfa* and *sox10*, suggesting that elf1 represses the migratory states and mesenchymal fates (Supplementary Fig. 9c).

**Figure 6.**
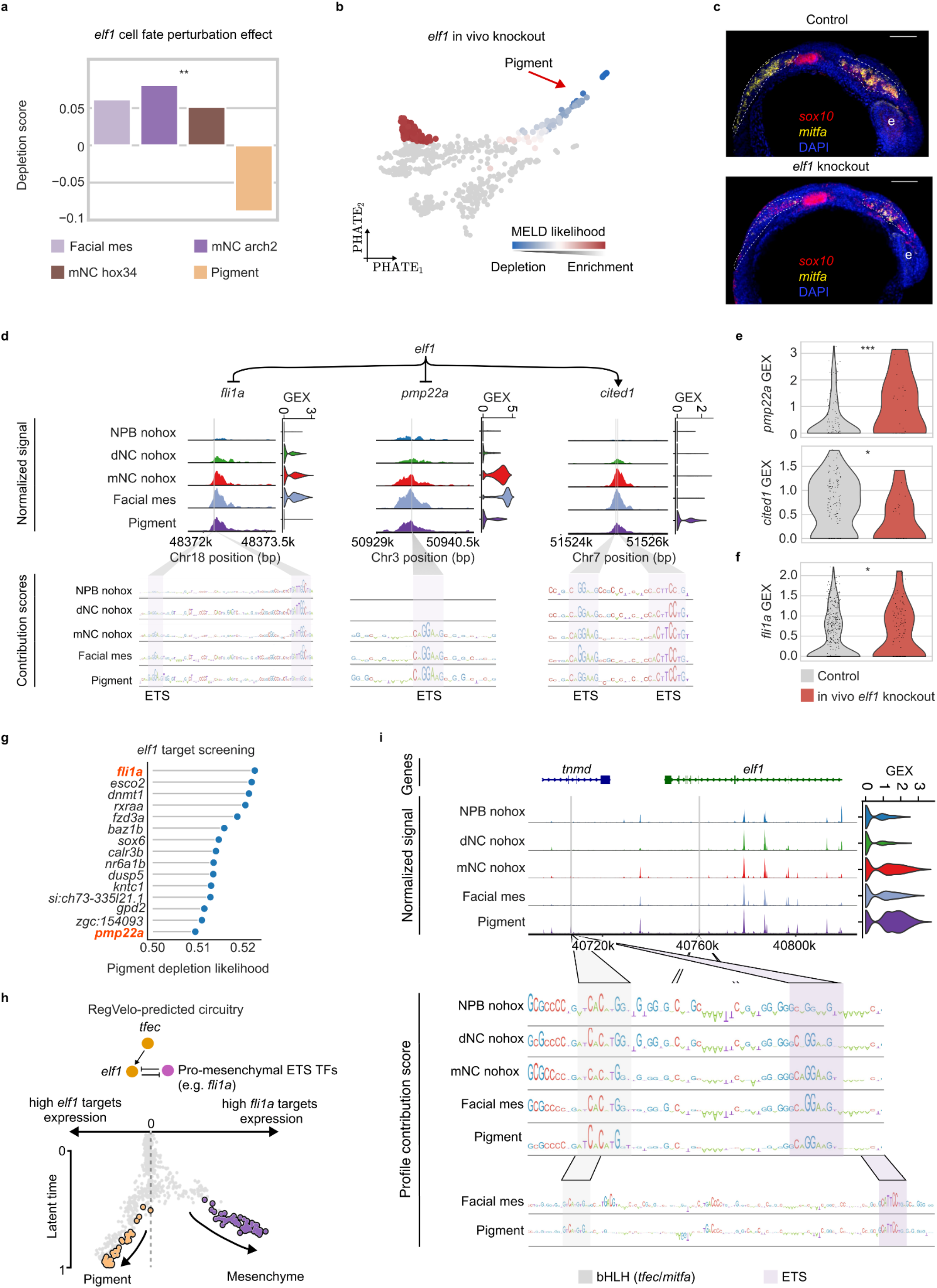
RegVelo identifies an early pigment lineage driver, *elf1*, together with its regulatory context. **a**. Cell fate perturbation effects estimated from elf1 regulon knockout simulation. **b**. PHATE embedding colored by in-vivo *elf1* perturbation scores quantified by MELD. **c**. HCR demonstrates a reduction in *mitfa*-expressing pigment lineage in *elf1* crispants compared to the wild-type embryo at 21 ss. **d**. Putative target genes *fli1a, pmp22a*, and *cited1* downstream of elf1, based on chromatin accessibility of *cis*-regulatory element candidates (top left track plots), the cluster-specific gene expression (top right violin plots), and the highlighted ETS TF binding instances (bottom). **e**. Violin plots show the expression of top targets genes between control and *elf1* knockout for the pigment cell lineage in the in vivo Perturb-seq dataset (*N* = 157 cells) (adjusted *P* = 2.4 × 10^−10^ for *pmp22a, P* = 0.02 for *cited1*, by edgeR). **f**. Violin plot of *fli1a* gene expression in Perturb-seq dataset (*N* = 779 cells) of all neural crest cells in two conditions (adjusted *P* = 0.004, by limma DE test). **g**. The top ten RegVelo-predicted targets of *elf1* affected most in the pigment lineage. **h**. RegVelo’s predictions suggest that a toggle-switch circuitry governs the cell fate decision between the pigment and mesenchymal lineages (top). We visualize the branching point and highlight the balance between these gene regulatory programs via the log_2_-fold change of the average expression levels of all fli1a targets versus all elf1 targets in each cell (x-axis) along the cell differentiation latent time inferred by RegVelo (y-axis). **i**. Two candidate *cis-*regualtory elements of *elf1* contain the TF binding instances of bHLH family TFs and ETS family TFs.

To decipher the regulatory mechanism of elf1, we investigated its downstream targets, next. By integrating chromatin accessibility profiles and ChromBPNet’s contribution scores, we identified the ETS TF binding sites upstream of *fli1a*^*82*^, *pmp22a*^*102*^, *arl6ip1*^*103*^, *hmgn2*^*104*^, and the novel pigment lineage marker *cited1* (Figure 6d,e and Supplementary Fig. 9d,e). Based on the Perturb-seq data, we confirmed through differential expression analysis that the target genes *pmp22a* and *cited1* were significantly differentially expressed in the *elf1*-knockout pigment lineage, while the *fli1a* expression was upregulated in *elf1-*knockout neural crest and derivative cells, suggesting a compensatory mechanism (adjusted *P* < 0.02; Figure 6e,f). This indicated these genes as direct candidate targets downstream of elf1. We further screened generic circuits with RegVelo using in silico knockouts to assess the effect of perturbing downstream regulatory connections of elf1 on pigment differentiation. This screening identified *fli1a* and *pmp22a* downstream of *elf1* as top perturbed factors, revealing potential regulatory mechanisms linking *elf1* to downstream pathways of neural crest migration (Figure 6g).

Finally, we sought to identify the core subnetwork of pigment fate priming (Figure 6h and Supplementary Fig. 9a): comparing the expression of targets ordered by RegVelo’s latent time revealed a clear bifurcation between pigment and mesenchymal fate with elf1 targets upregulated in the pigment lineage, and pro-mesenchymal ETS TFs upregulated correctly in the mesenchymal lineage^82^ (Figure 6h), again implying elf1’s active role in fate decision towards pigment cells. To further establish regulatory relationships of *elf1*, we identified putative TF binding sites of mitfa and tfec present in *elf1*’s enhancer regions (Figure 6i) and found that *elf1* expression was downregulated in tfec knockouts (Supplementary Fig. 9f). Upstream *cis*-regulatory elements of both *elf1* and *fli1a* included ETS factor binding sites, and the *fli1a* knockouts resulted in the upregulation of *elf1* (Figure 6d,i and Supplementary Fig. 9g).

Based on these findings, we have identified a core module within RegVelo’s complex GRN, represented as the following subnetwork: the early pigment driver tfec activates the pro-pigment TF *elf1*, and elf1 and pro-mesenchymal ETS TFs suppress each other to drive the neural crest cell fate bifurcation into pigment and mesenchymal fates, following a toggle switch model^105^.

## Discussion

We have presented RegVelo, a novel variational inference framework for learning RNA velocity as well as the underlying regulatory network jointly from single-cell gene expression data. The step from decoupled single gene models to a fully integrated dynamic model by explicitly incorporating gene regulation makes our method compare favorably to previous methods on dynamic inferences; we found significant improvements on simulated and real datasets in terms of velocity estimation, latent time compared to velocity models, and GRN inference compared to static network inference approaches. Notably, the forward simulation based on our learned gene regulatory dynamics provides a simple virtual cell^106^*-*style downstream analysis to decipher regulatory mechanisms in cell fate decision behavior.

In particular, we recovered murine differentiation trajectories and regulatory mechanisms during pancreatic endocrinogenesis: for the ductal cell cycling process, we accurately predicted drivers and their regulatory effects, and faithfully described lineage conformations by in silico simulation. We also showcased dynamic inference on a human hematopoiesis dataset where previous RNA velocity methods failed or performed worse even with access to more rich information like metabolic labels, recovered the toggle-switch network motif between GATA1 and SPI1 (PU.1), and illustrated, using RegVelo’s perturbation scheme, how these genes relate to the erythroid and monocyte-fate decisions.

While most ETS factors have pro-mesenchymal bias in zebrafish cranial neural crest^82^, RegVelo led us to report elf1 as an exception that terminates migration and promotes pigment fates. Our finding is consistent with the DNA-binding anti-cooperativity of ELF1 caused by an electrostatic repulsion mechanism, in contrast to FLI1 and ETS1^107^. Moreover, our finding also aligns with the classic theory of cell fate decisions that two transcriptional programs compete to drive bifurcation events^108^. In the zebrafish system, we speculate that the ETS family TF Elf1 directly and competitively binds the same TF binding site of other ETS factors and, thus, can take a contradictory role against these other factors, supporting the bifurcation mechanism at the regulatory level. Our newly found driver for the pigment lineage, elf1, has also been reported as the intermediate-state marker in melanoma^109^, where the tumor cell states intermediate or neural crest-like, mesenchyme-like, and melanoblast-like were reported to mirror normal cell states in the neural crest cellular trajectories^110^. Future studies can investigate whether elf1 is involved in a shared regulatory circuitry between neural crest fate decisions and melanoma. Overall, our systematic in silico and in vivo TF knockout datasets serve as valuable resources for studying cranial neural crest development and relevant disease, as well as for benchmarking the accuracy of knockout simulation algorithms.

As an end-to-end dynamic, interpretable, and actionable model, RegVelo distinguishes itself from other studies that focus on interpreting cell dynamics through gene regulation and often require nonlinear models like support vector machines or variational autoencoders to predict RNA velocity from the transcriptome^22,111,112^. RegVelo offers significant advantages over these methods in three aspects: (1) These approaches depend on predefined cellular velocity estimates^22,111,112^ from models ignoring regulatory events^13,14^. In contrast, RegVelo simultaneously learns gene regulation and cellular velocity. (2) RegVelo integrates prior knowledge about GRN structure learned from various modalities, such as chromatin accessibility, by incorporating gene regulation into velocity estimation. (3) Previous models for estimating splicing dynamics predict perturbation effects by manually activating or suppressing gene expression^22,111^, where the output represents the immediate perturbed direction of each cell. Importantly, classical RNA velocity approaches study completely decoupled systems, thereby ignoring gene-gene dependencies, and, thus, they cannot predict perturbation effects. RegVelo, however, encodes perturbation by directly changing the underlying regulatory circuits and simulating gene expression dynamics under the perturbed regulatory system. This forward simulation allows for the prediction of nonlinear perturbation effects over latent time rather than focusing only on the immediate first-order direction, which misses long-term consequences. We demonstrated RegVelo’s ability to predict such long-term effects of regulators by successfully recovering early-expressed lineage drivers in zebrafish neural crest cell development. Notably, we used in vivo perturbation experiments to validate the TF perturbation effect prediction across four lineages. Our results demonstrated a significant improvement over existing dynamics-based perturbation prediction methods and our GRN-infused hybrid model holds promise to be more robust on out-of-distribution data than purely data-based generative models^113^.

RegVelo stands as the first model to extend the currently employed RNA velocity model from a pure de novo discovery tool for trajectory inference into a simulator for gene regulatory dynamics but leaves room for future extensions. The deep integration between RegVelo and CellRank offers a causal mechanism linking gene regulation and changes to cell fate decisions, as opposed to a merely correlation-based approach. Meanwhile, as an extension of veloVI, introducing prior information on gene regulation and model regularization, RegVelo leads to better identifiability on several downstream statistics compared to veloVI. However, in a more general modeling scenario, in particular in complex or understudied systems, proper accurate prior information on gene regulation is unfeasible as a rich literature background is lacking or epigenetic information for scATAC-seq, for example, is sparse or does not exist. As shown, RegVelo can mitigate the effect of a poor prior, but we anticipate that incorporating additional types of prior information, such as pseudotime, real time, or the number of lineages, regularizes the model and thereby improves the inference of gene regulatory dynamics further. While this incorporation may contravene its inherent characteristics as an unsupervised trajectory inference tool, RegVelo also serves as a simulator of transcriptional regulation, providing insights into gene regulation and cell fate priming that are far more valuable than merely recapitulating the overall trajectory.

Recent experimental advances have enabled the measurement of various modalities. Built upon veloVI, RegVelo is a flexible model that can easily generalize to additional modalities to overcome current limitations in model formulation and assumptions. For example, incorporating metabolic labeling data or in vivo time-resolved scRNA-seq data like Zman-seq^114^ enables the development of new differential equations to describe transcriptional regulation and infer cell-specific kinetic rates, such as degradation^22^. Similarly, cross-linking immunoprecipitation (CLIP) and Deamination Adjacent to RNA modification Target (DART) sequencing may allow for estimating cell-specifc splicing rates. The two protocols have provided insights into RNA binding protein (RBP)-RNA interactions at the single-molecule level^115,116^ and combining such post-translational gene-gene interactions can improve the regulatory modeling among different molecules. Moreover, while RegVelo models transcription in a context-specific manner, we and other existing frameworks typically assume that gene expression depends on the abundance of upstream RNA regulators rather than protein levels - integrating ribosome profiling data enables modeling the translation process and quantifying the protein activity of regulators which enables a more nuanced estimation of transcription rates for target genes^117^. Finally, extending the biophysical model of splicing kinetics beyond mRNA toward chromatin and protein dynamics will lead to a more complete and accurate description of the underlying process. How to correctly incorporate these alternative views is unclear or not yet possible, though, as the data types differ from transcriptomic count data, and shared measurements in the same cell do not yet exist.

Overall, RegVelo innovates transcriptome-wide dynamic modeling from pure trajectory inference to uncovering causal mechanisms involved during cell fate decisions. With this flexible model as a prototype, we expect future extensions by incorporating additional regulatory layers at different molecular levels, ultimately moving towards a more comprehensive and interpretable model capable of simulating and predicting cell behavior and responses even more accurately.

## People

We thank all members of the Theis and Sauka-Spengler laboratories for helpful discussions. W.W. thanks Dr. Xiaojie Qiu for his feedback on RegVelo concerning implementation and benchmarking aspects, A. Palma and M. Lienen for helpful discussions about parallel ODE solvers and neural ODE models, D. Klein for insights into pancreatic endocrine development and L. Kummerle for discussing RegVelo perturbation benchmark results. We thank the MRC WIMM Flow Cytometry Facility, MRC WIMM Advanced Single Cell OMICS Facility, MRC WIMM Centre for Computational Biology, MRC WIMM and Stowers Institute Sequencing Facilities, and Wolfson Imaging Centre for their help and technical support in the zebrafish study part. Z.H. thanks the HPC core facility of the Medical Research Institute at Wuhan University for technical support. Scientific illustrations shown in Figure 4a were created by Uta Mackensen.

## Author’s contributions

W.W., Z.H. and P.W. contributed equally. W.W., P.W., M.L., F.J.T. conceptualized the study, and Z.H., T.S.-S. conceptualized the zebrafish study design. W.W. designed and implemented RegVelo with contributions from P.W. and M.L. Z.H. and S.M. performed wet-lab experiments involving zebrafish. W.W., Z.H., P.W. designed and implemented analysis methods with contributions from J.W. and Z.X. T.S.-S. and F.J.T. supervised the work. P.W., W.W., Z.H., T.S.- S. and F.J.T. wrote the manuscript. All authors read and approved the final paper.

## Data availability

The cell cycle, human hematopoiesis, and pancreatic endocrinologist data presented and used in this study are publicly available via the original publications; we provide additional access to each of these datasets and the zebrafish data via a figshare collection at https://figshare.com/account/home#/projects/226860. The zebrafish data is available at https://research.stowers.org/compbio/ucsc_cellbrowser/.

## Code availability

RegVelo is released under the BSD-3-Clause license, with code available at https://github.com/theislab/regvelo. Code to reproduce the results in the paper can be found at https://github.com/theislab/regvelo_reproducibility.

## Supplementary Materials

Supplementary Figures: Supplementary Figures 1-9

Supplementary Notes: Supplementary Notes 1, 2

Supplementary Tables: Supplementary Tables 1, 2

## Supplementary Figures

**Suppl. Fig 1.**
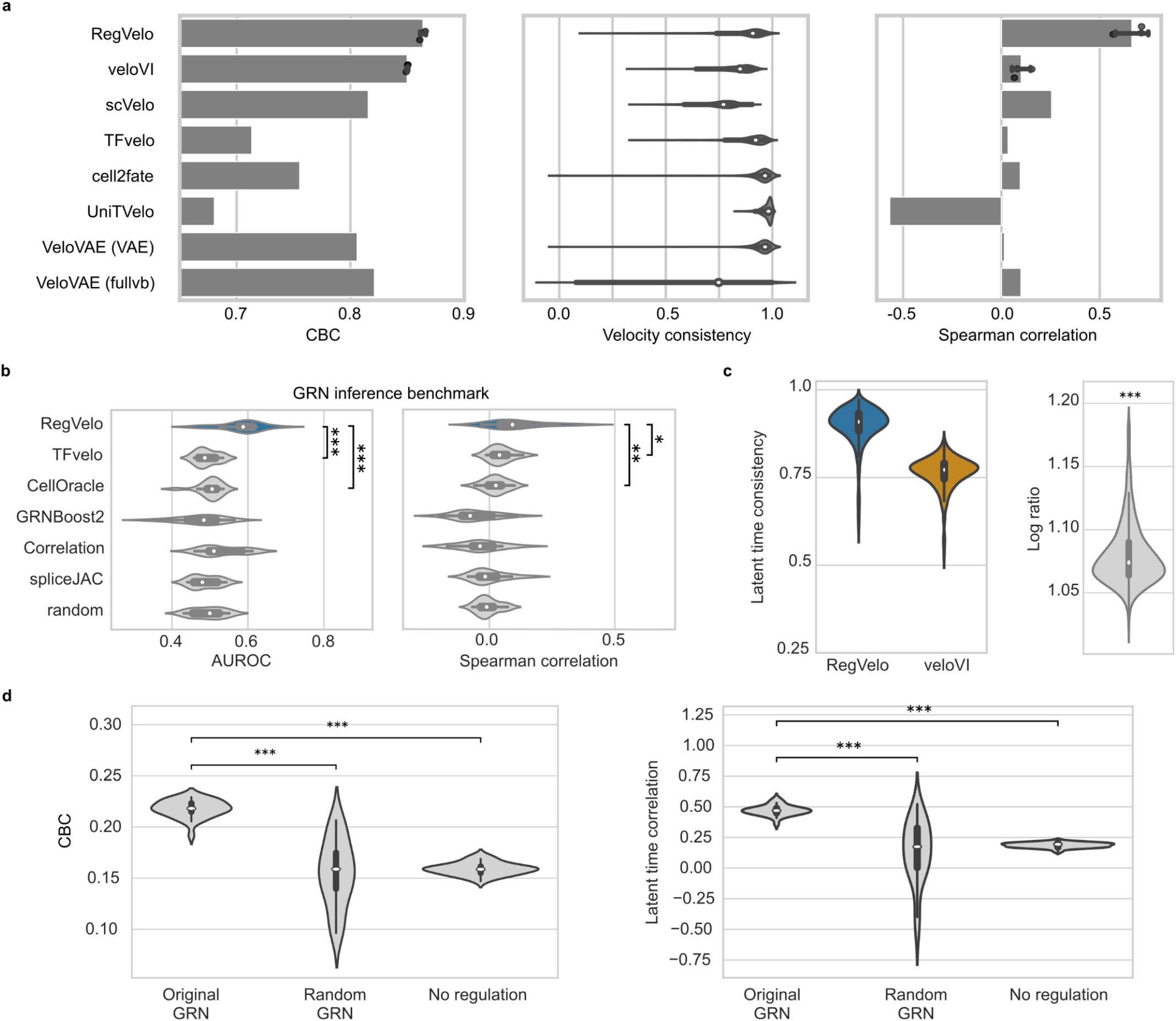
Benchmarking RegVelo against competing approaches on a cell cycling dataset. **a**.Cross-boundary correctness (CBC) score of RegVelo and veloVI (*N* = 3 trained models), and six other velocity inference methods (left), velocity consistency (middle) and correlation between ground truth and inferred latent times (*N* = 3 trained models for RegVelo and veloVI). Box plots indicate the median (center line), interquartile range (hinges), and 1.5x interquartile range (whiskers) (*N* = 1146 cells). **b**. Latent time consistency (left) and log_2_ ratio (right) of RegVelo’s and veloVI’s consistency score. Box plots indicate the median (center line), interquartile range (hinges), and 1.5x interquartile range (whiskers) (*N* = 1146 cells (left) and ratios (right) each). We used a one-sided Welch’s t-test to assess significance (*P* < 1 × 10^−10^). **c**. Comparison of CBC score (left) and Spearman correlation between ground truth and estimated latent time (right), using original GRN (original), randomly shuffled GRN (random) or no GRN (No regulation) to run RegVelo (*N* = 30 repeats). We used a one-sided Welch’s t-test to assess significance (*P* < 1 × 10^−10^); box plots indicate the median (center line), interquartile range (hinges), and 1.5x interquartile range (whiskers) (*N* = 30 runs). **d**. Comparison of GRNs inferred by GRN inference methods and a random baseline to a ChIP-Atlas-derived proximal ground truth GRN. We assessed performance based on an AUROC (left) and Spearman correlation between ChIP-Atlas-derived TF-target binding score and weight estimates (right). We used a one-sided Welch’s t-test to assess significance (AUROC: RegVelo vs. TFvelo = *P* = 8.7 × 10^−6^, RegVelo vs. CellOracle *P* = 1.19 × 10^−6^; Spearman correlation: RegVelo vs. TFvelo *P* = 0.019, RegVelo vs. CellOracle *P* = 0.0016). Box plots indicate the median (center line), interquartile range (hinges), and 1.5x interquartile range (whiskers) (*N* = 15 TFs).

**Suppl. Fig 2.**
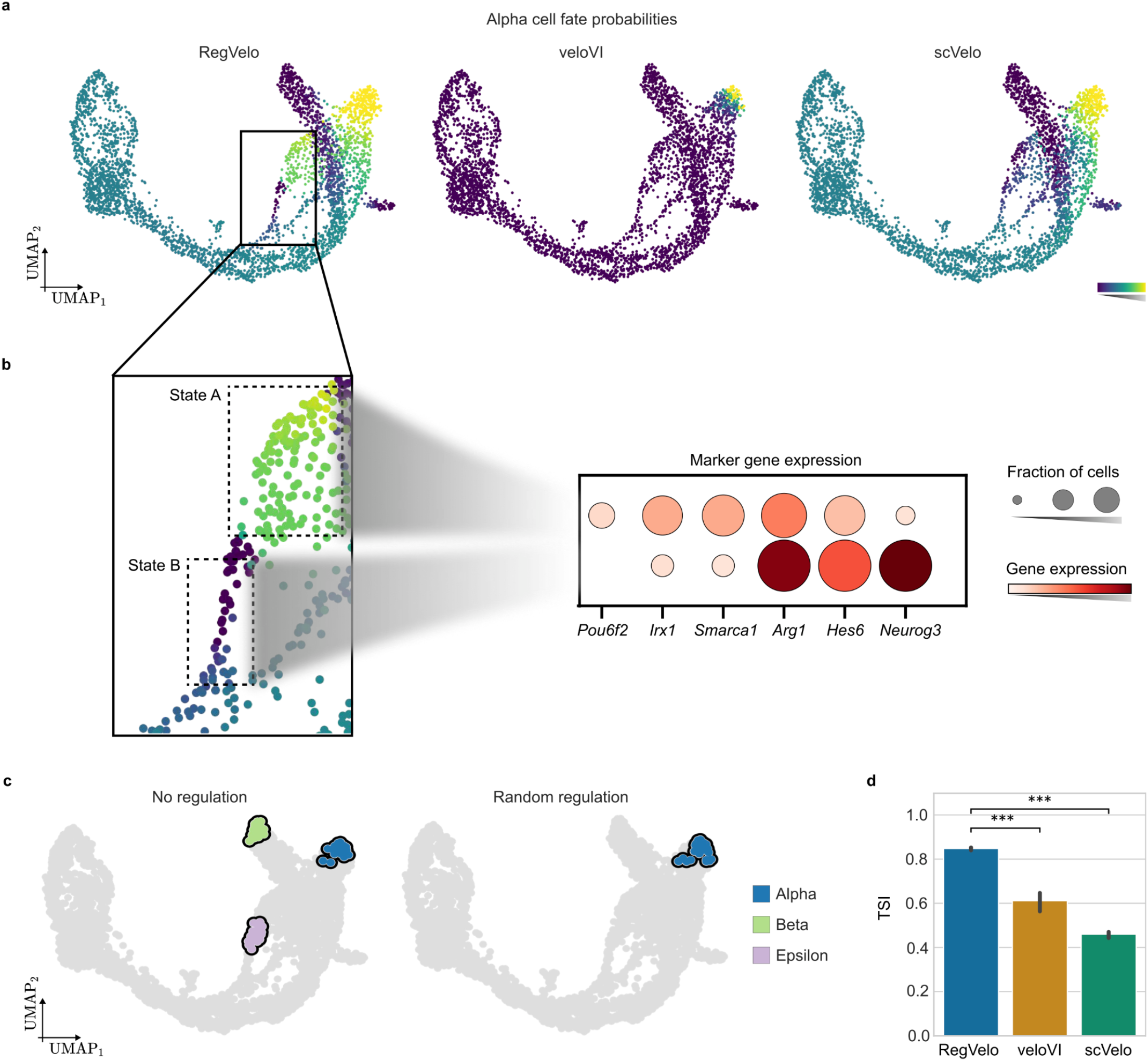
Benchmarking RegVelo on cell fate prediction with the pancreatic endocrine dataset. **a**. UMAP representation of pancreatic endocrinogenesis data, colored by fate probabilities using velocity estimates from RegVelo (left), veloVI (middle), and scVelo (right). **b**. UMAP embedding from **a**. zoomed in on Alpha cells (left) and marker gene expression in two substates (right; Methods). **c**. UMAP representation of pancreatic endocrinogenesis data, colored by identified terminal states using CellRank with RegVelo when omitting any regulation (left) or relying on a random GRN (right; Methods)). **d**. Comparison of the terminal state identification (TSI) score using RegVelo (blue), veloVI (orange), and scVelo (green) (*N* = 20 runs). We used a one-sided Welch’s *t*-test to assess significance (RegVelo vs. veloVI: P = 9.23 × 10^−8^; RegVelo vs. scVelo: *P* < 1 × 10^−10^; error bars correspond to 95% confidence intervals.

**Suppl. Fig 3.**
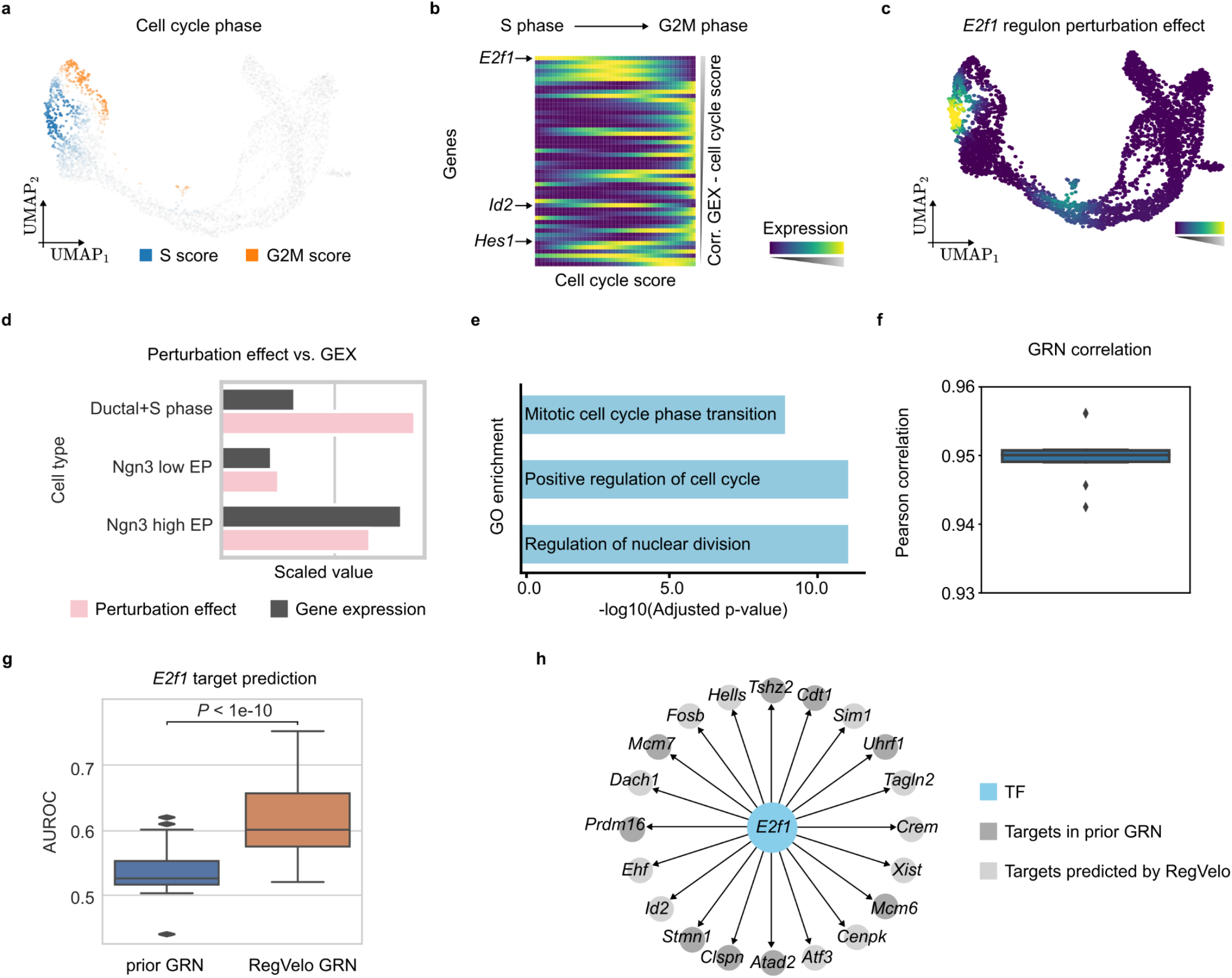
*E2f1* in silico perturbation analysis. **a**. UMAP embedding of pancreatic endocrinogenesis data colored by inferred S and G2M phases based on cell cycle score derived from gene expression. **b**. Smoothed gene expression along the experimentally derived cell cycling staging for the top 50 cell cycle score-correlated TFs, sorted by correlation from high to low. **c**. UMAP embedding of the pancreas dataset colored by local perturbation effects of *E2f1* regulon. **d**. Comparison of perturbation score and expression of *E2f1* in different cell types. **e**. Top 3 enriched GO pathways for the RegVelo predicted *E*2f1 targets; we used Fisher’s exact test to assess the significance and the Benjamini and Hochberg (BH) method to adjust the *p*-value. **f**. Correlation of RegVelo inferred GRN derived from pairs of runs of the RegVelo model. Box plots indicate the median (center line), interquartile range (hinges), and 1.5x interquartile range (whiskers) (*N* = 5 runs). **g**. Comparison of the AUROC of the GRN prediction using the prior GRN (blue) and RegVelo-inferred GRN (orange) (Methods). We used a one-sided Wilcoxon signed-rank test to assess significance (*P* < 1 × 10^−10^). Box plots indicate the median (center line), interquartile range (hinges), and 1.5x interquartile range (whiskers) (*N* = 100 comparisons). **h**. Top 20 targets of *E2f1* predicted by RegVelo.

**Suppl. Fig 4.**
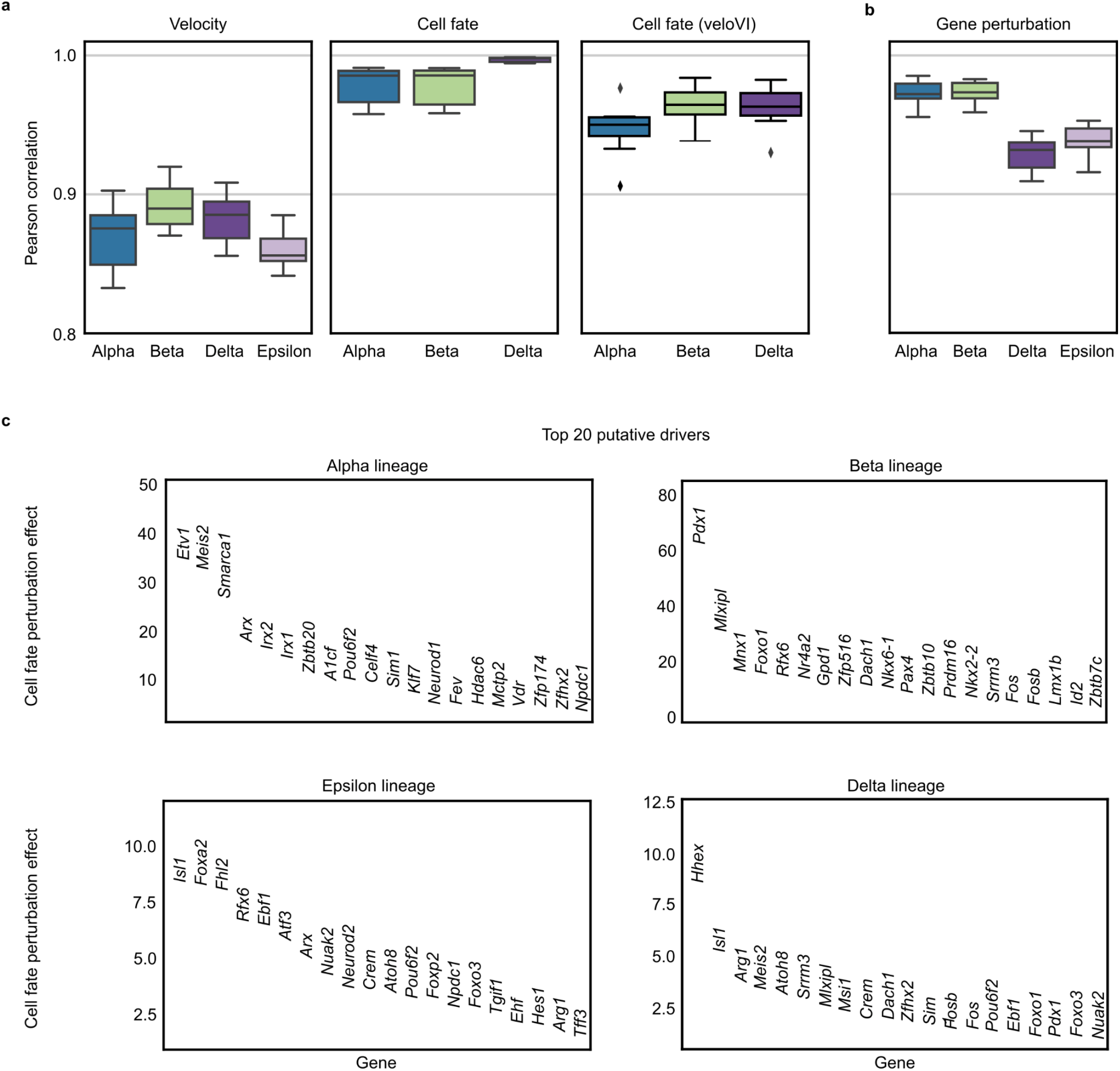
Identifying regulators related to cell fate decision during pancreatic endocrine. **a**,**b**. Pearson correlation of RegVelo inferred velocities (**a**., left), CellRank-computed cell fate probabilities based on RegVelo’s (**a**., middle) and veloVI’s (**a**., right) velocity estimates, and cell fate perturbation score across repeated model fits (**b**.). Box plots indicate the median (center line), interquartile range (hinges), and 1.5x interquartile range (whiskers) (*N* = 5 runs each). **c**. Top 20 putative lineage drivers identified with RegVelo’s estimate of cell fate perturbation effects.

**Suppl. Fig 5.**
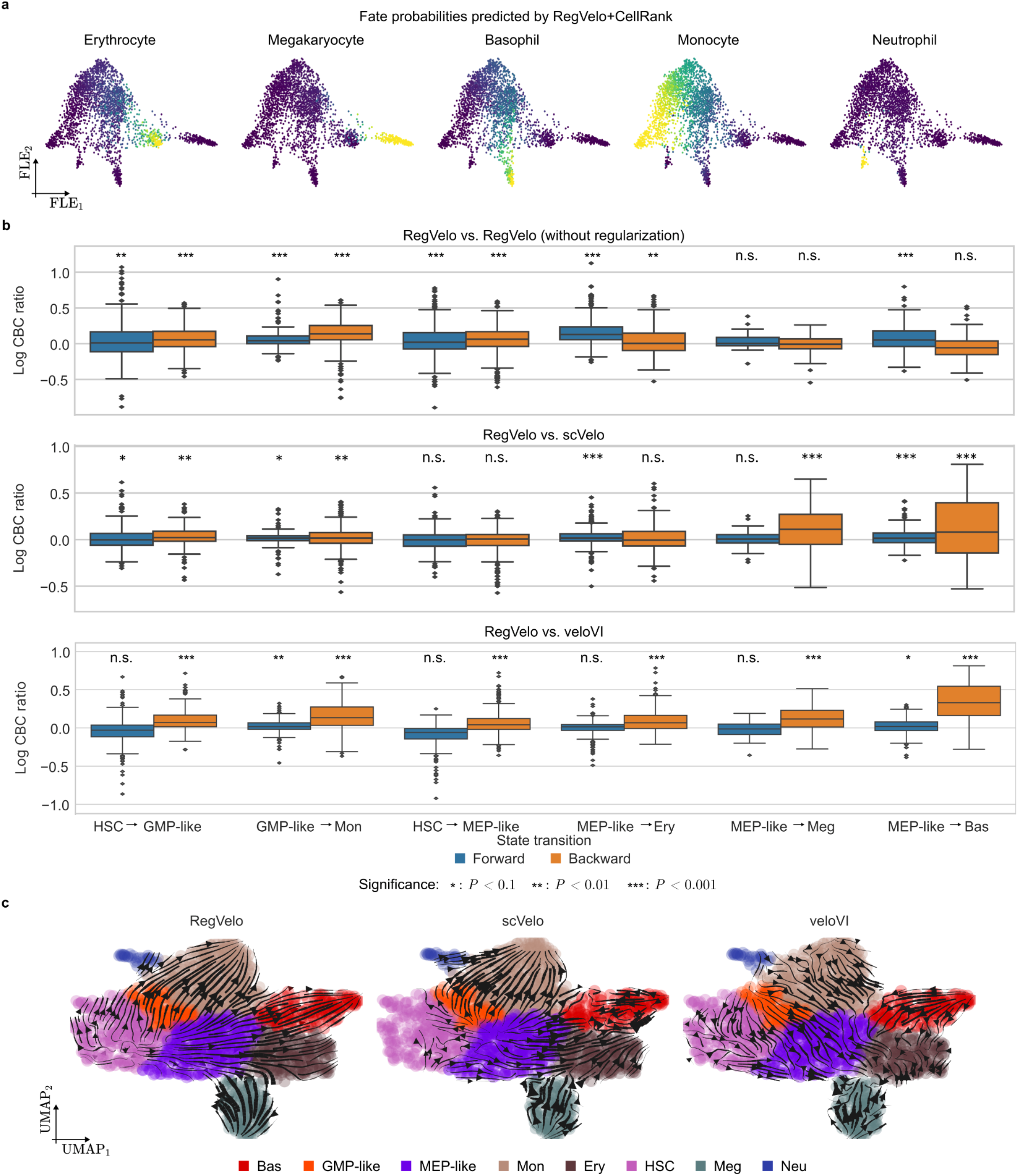
Benchmarking velocity inference for human hematopoiesis. **a**. CellRank-computed fate probabilities towards each identified terminal state using RegVelo-inferred velocities. **b**. Log-transformed ratio of cross-boundary correctness score considering both forward and backward flows of cell type transitions of RegVelo with Jacobian regularization (RegVelo), RegVelo without Jacobian regularization (RegVelo (without regularization)) (top), scVelo (middle) and veloVI (bottom) (HSC: hematopoietic stem cell, Meg: megakaryocyte, Ery: erythrocyte, Bas: basophil, MEP: megakaryocyte and erythrocyte progenitor-like, GMP: granulocyte and monocyte progenitor; Methods). We used one-sided Welch’s t-tests to test significance (Methods). Box plots indicate the median (center line), interquartile range (hinges), and 1.5x interquartile range (whiskers) (upper: HSC to GMP-like forward: *N* = 304 cells, *P* = 0.001; HSC to GMP-like backward *N* = 161 cells, *P* = 6.39 × 10^−6^; GMP-like to Mon forward: *N* = 152 cells, *P* = 1.23 × 10^−7^; GMP-like to Mon backward: *N* = 376 cells, *P* = 8.24 × 10^−38^. HSC to MEP-like forward: *N* = 309 cells, *P* = 0.0001; HSC to MEP-like backward: *N* = 446 cells, P = 3.17 × 10^−10^. MEP-like to Ery forward: *N* = 312 cells, *P* = 6.46 × 10^−41^; MEP-like to Ery backward: *N* = 220 cells, *P* = 0.012. MEP-like to Meg forward: *N* = 16 cells, *P* = 0.157; MEP-like to Meg backward: *N* = 66 cells, *P* = 0.835;. MEP-like to Bas forward: *N* = 161 cells, *P* = 1.33 × 10^−6^; MEP-like to Bas backward: *N* = 144 cells, *P* = 0.99. Middle: HSC to GMP-like forward: *P* = 0.032; HSC to GMP-like backward: P = 0.0008. GMP-like to Mon forward: P = 0.031; GMP-like to Mon backward: P = 0.0008. HSC to MEP-like forward: *P* = 0.58; HSC to MEP-like backward: P = 0.99. MEP-like to Ery forward: *P* = 1.71 × ^10−5^; MEP-like to Ery backward: *P* = 0.18. MEP-like to Meg forward: *P* = 0.31; MEP-like to Meg backward: *P* = 6.53 × 10^−6^. MEP-like to Bas forward: *P* = 0.0016; MEP-like to Bas backward: *P* = 1.16 × 10^−7^. Bottom: HSC to GMP-like forward: *P* = 0.099; HSC to GMP-like backward: *P* = 4.93 × 10^−16^. GMP-like to Mon forward: *P* = 0.012; GMP-like to Mon backward: *P* = 1.96 × 10^−44^; HSC to MEP-like forward: *P* = 1; HSC to MEP-like backward: *P* = 2.13 × 10^−18^. MEP-like to Ery forward: *P* = 0.453; MEP-like to Ery backward: *P* = 4.61 × 10^−18^. MEP-like to Meg forward: *P* = 0.655; MEP-like to Meg backward: *P* = 1.02 × 10^−7^. MEP-like to Bas forward: *P* = 0.038; MEP-like to Bas backward: *P* = 1.19 × 10^−39^. **c**. Velocity stream embedding of the velocity estimates using RegVelo (left), scVelo (middle), or veloVI (right).

**Suppl. Fig 6.**
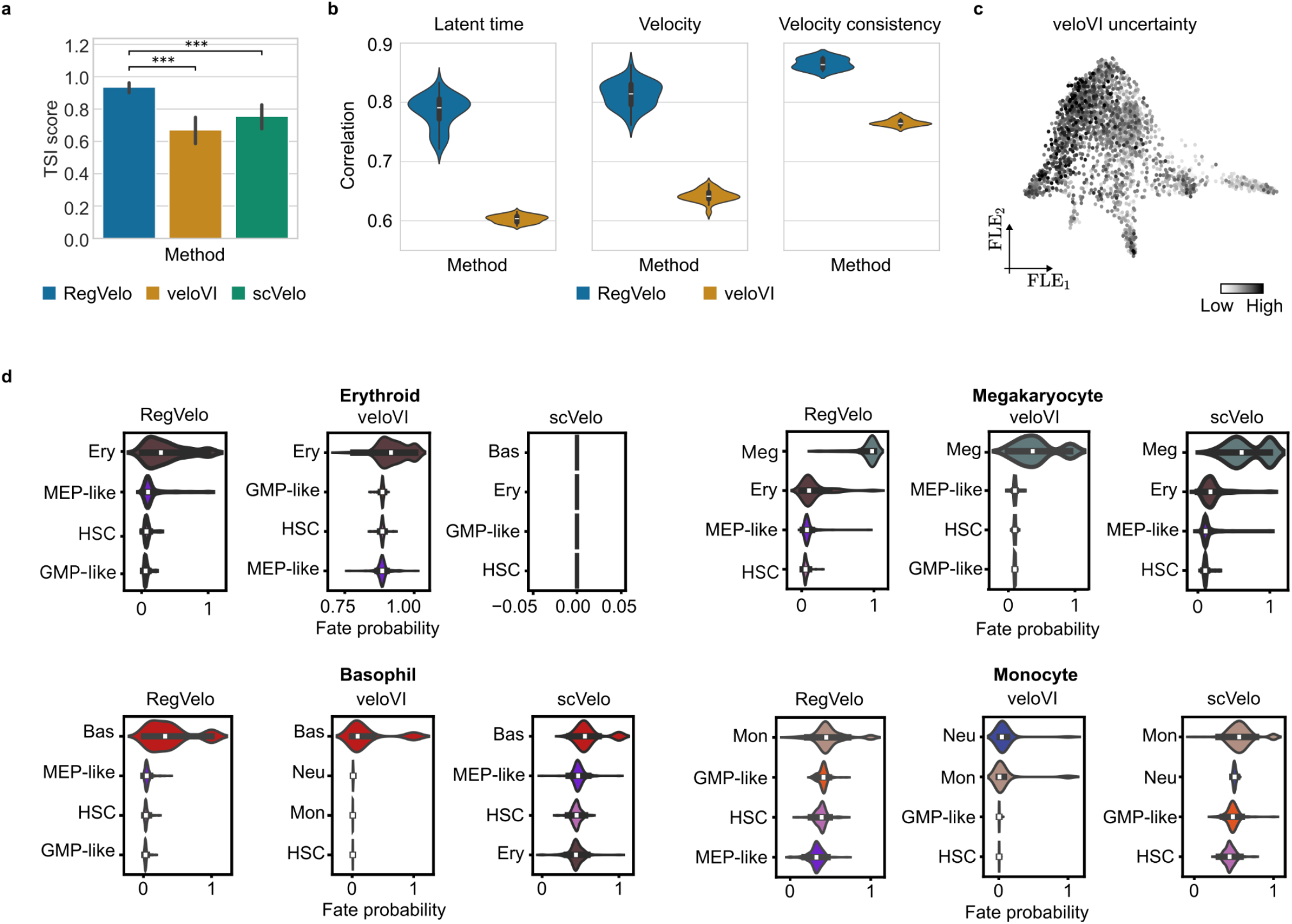
Benchmarking cell fate prediction for human hematopoiesis. **a**. TSI score using RegVelo (blue), veloVI (orange), and scVelo (green) (one-sided Welch’s *t*-tests, RegVelo vs. veloVI: *P* = 1.59 × 10^−5^; RegVelo vs. scVelo: *P* = 2.23 × 10^−5^). Error bars correspond to 95% confidence intervals (*N* = 20). **b**. Correlation of latent time and velocity estimates and velocity consistency across multiple runs using RegVelo and veloVI. Box plots indicate the median (center line), interquartile range (hinges), and 1.5x interquartile range (whiskers) (*N* = 5 runs). **c**. FLE embedding of hematopoiesis data colored by veloVI’s cell uncertainty. **d**. Distribution of CellRank’s fate probabilities towards four terminal states using RegVelo’s, veloVI’s, and scVelo’s velocities. For each lineage, we show the four cell types with the highest median cell fate probability for each method in the lineage.

**Suppl. Fig 7.**
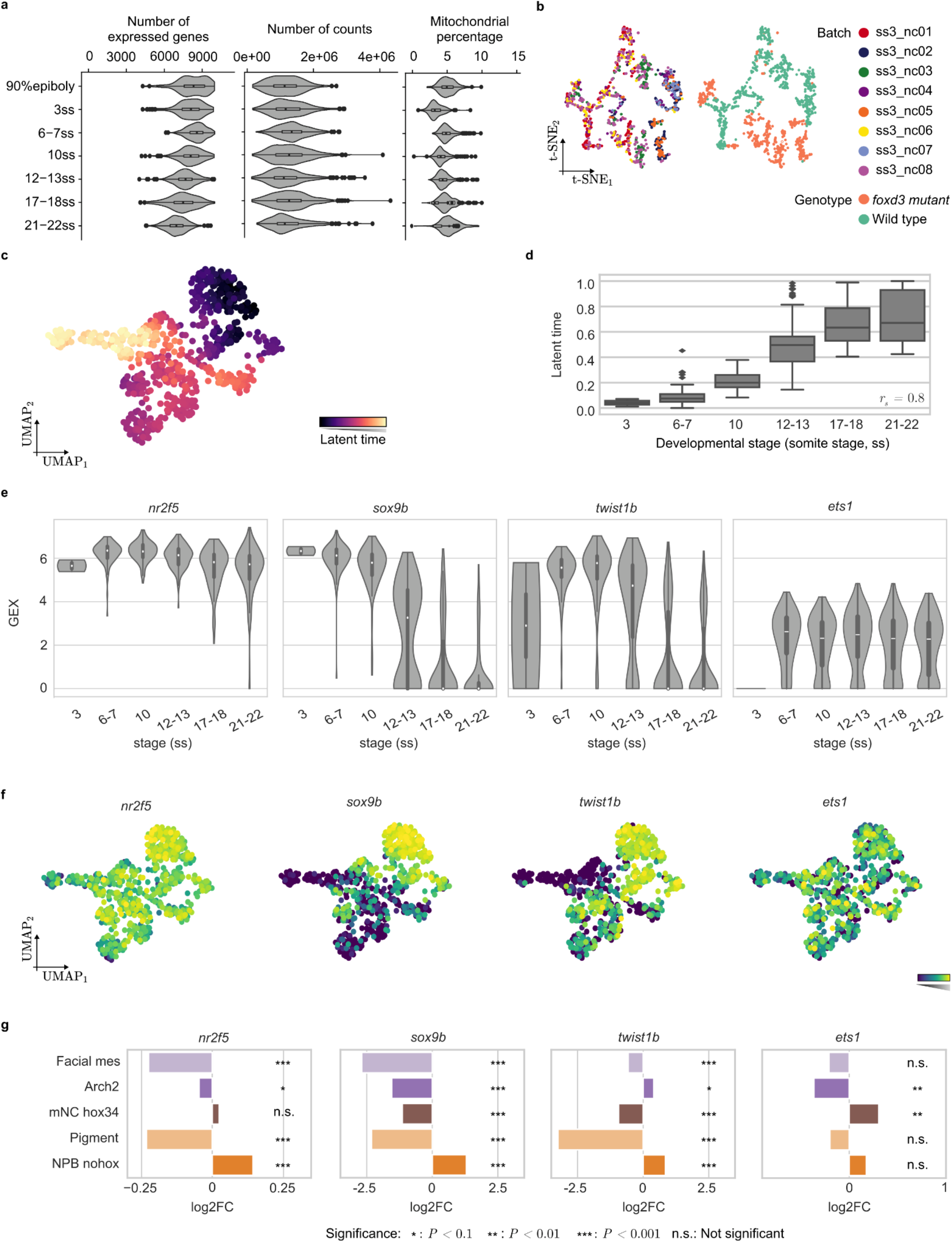
Gene expression patterns of lineage drivers during neural crest development. **a**. Violin plots show consistent quality across different developmental stages in the Smart-seq3 data. **b**. The eight distinct batches in the Smart-seq3 data overlap in the corresponding *t*-SNE embedding, signifying the absence of batch effects (left). At the same time, distinct cellular patterns between wild-type and *foxd3*-mutant neural crest cells exist (right). **c**. UMAP embedding colored by the RegVelo-inferred latent time. **d**. The distribution of the latent time stratified by embryonic developmental stages. Box plots indicate the median (center line), interquartile range (hinges), and 1.5x interquartile range (whiskers) (E3: *N* = 10 cells, E6-E7: *N* = 105 cells, E10: *N* = 105 cells, E12-E13: *N* = 156 cells, E17-E18: *N* = 146 cells, E21-E22: *N* = 183 cells). **e**. Distribution of gene expression of *nr2f5, sox9b, twist1b*, and *tfap2a*, stratified by embryo stage information. Box plots indicate the median (center line), interquartile range (hinges), and 1.5x interquartile range (whiskers) (E3: *N* = 10 cells, E6-E7: *N* = 105 cells, E10: *N* = 105 cells, E12-E13: *N* = 156 cells, E17-E18: *N* = 146 cells, E21-E22: *N* = 183 cells). **f**. UMAP embeddings colored by gene expression levels of *nr2f5, sox9b, twist1b*, and *ets1*. **g**. Log-transformed fold change of *nr2f5, sox9b, twist1b*, and *ets1* gene expression in five cell types (terminal or initial states). We used Wilcoxon rank-sum tests to test the significance per gene and per cell type.

**Suppl. Fig 8.**
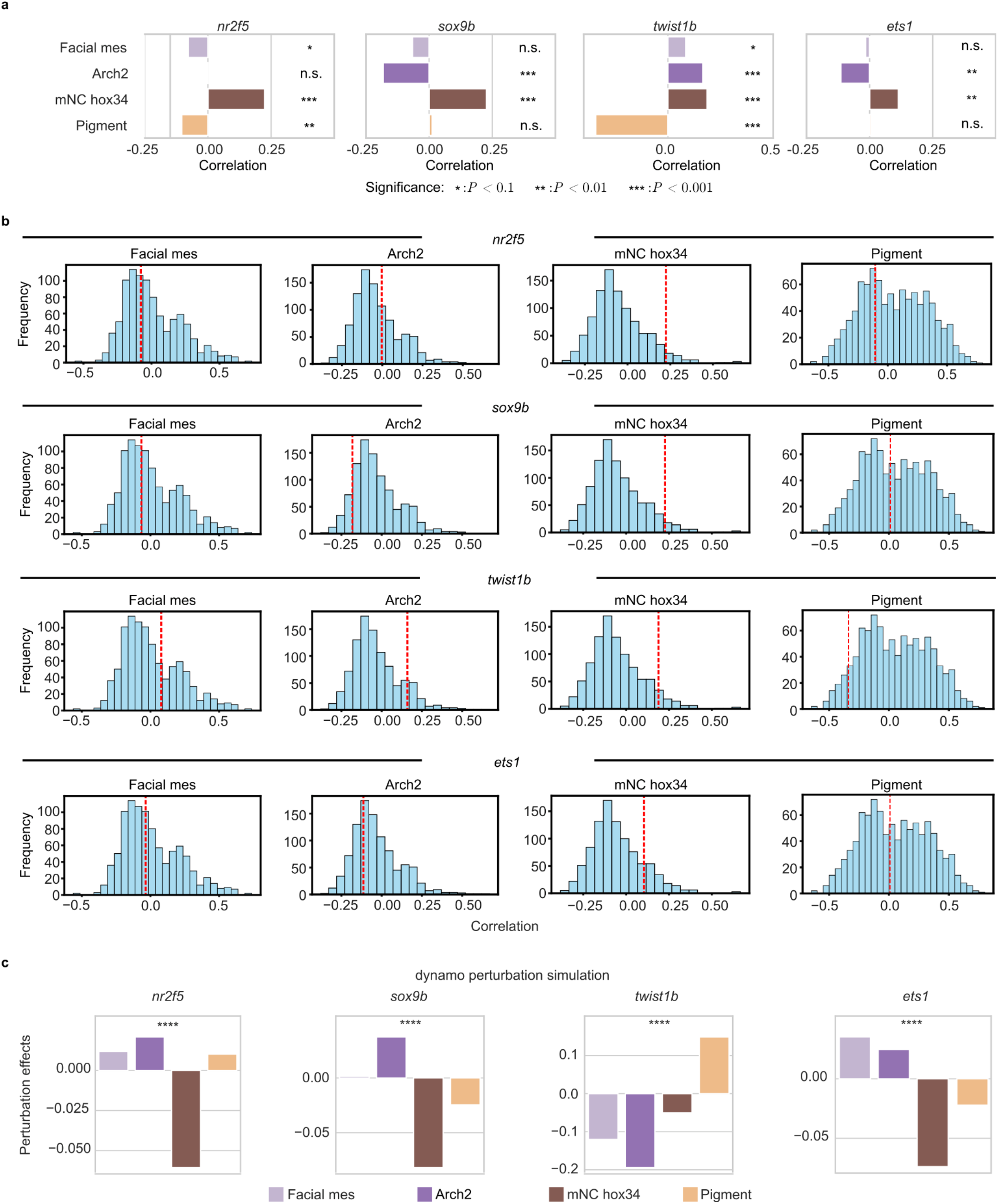
CellRank and Dynamo wrongly predict lineage drivers. **a**. CellRank-predicted Pearson correlation between gene expression and cell fate probabilities. We used the Welch’s t-tests to assess the significance. Each bar is colored according to the color code in Figure 4b. **b**. Distribution of the Pearson correlation between gene expression and fate probabilities for each lineage. The red dashed lines indicate the correlation between the corresponding TF (*nr2f5, sox9b, twist1b*, and *tfap2a*) expression and cell fate probability (*N* = 952 genes). **c**. Dynamo-predicted lineage perturbation effects of four cranial mesenchymal drivers indicate that in-silico knockouts of three out of four TFs show no depletion in mesenchymal fates.

**Suppl. Fig 9.**
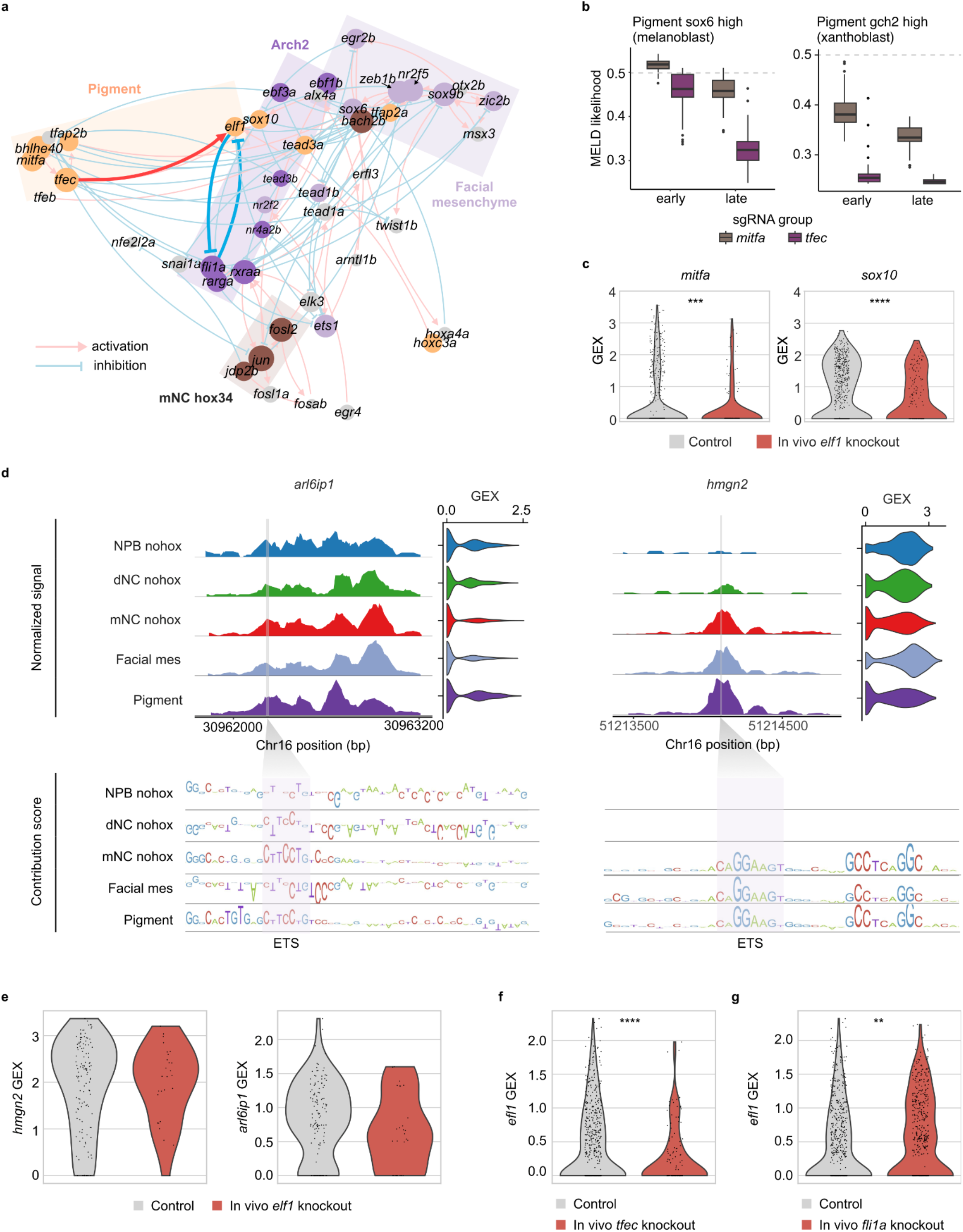
*elf1* is identified as an early pigment lineage driver, supported by its regulatory context. **a**.UMAP embedding of the inferred TF-TF network, where the distance represents the Jaccard index between TF pairs. The visualization highlights regulatory interactions between TFs inferred by RegVelo. The dot colors represent the TF-associated lineage as determined by RegVelo perturbation simulations, while the dot size indicates the predicted effect size of perturbations on the corresponding lineage; groups of closely related TFs were manually highlighted. **b**. In vivo Perturb-seq analysis shows profound depletion effects for *tfec* knockouts compared to *mitfa* in both early and late pigment lineages **c**. *mitfa* and *sox10* are significantly downregulated in *elf1*-knockout neural crest cells based on Perturb-seq data. **d**. Putative target genes *arl6ip1* and *hmgn2* downstream of *elf1*, with the ETS binding instances identified in their candidate *cis*-regulatory elements. **e**. Expression levels of *hmgn2* and *arl6ip1* are reduced but insignificantly in *elf1-*knockout pigment cells, likely due to synergistic transcriptional regulation. **f**. *elf1* expression levels are significantly decreased in *tfec* knockout, suggesting that *elf1* is downstream of *tfec*. **g**. *elf1* expression levels are significantly increased in *fli1a* knockout, supporting the toggle switch model between *elf1* and *fli1a*.

## Methods

### RegVelo model specification

Approaches for inferring RNA velocity models describe splicing dynamics through biophysical models like systems of ordinary differential equations (ODEs)^118^. Such models describe the transcription of precursor (unspliced) RNA *u* at rate α, followed by splicing into mature (spliced) mRNA *s* with rate β; spliced mRNA degrades at a rate *γ*, manifesting in the first-order approximation of a chemical master equation for gene *g* as

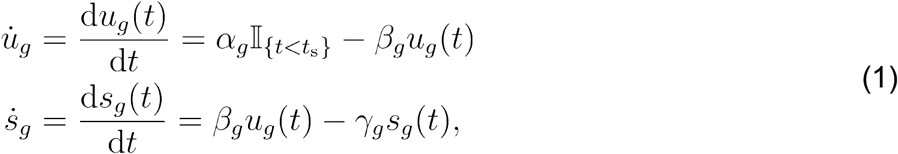

where 𝕀{*t*<*t*_*s*_} denotes the indicator function and *t*_*s*_ is a switch time. Throughout this work, we assume *N*_*C*_ cells and *N*_*G*_ genes.

Previously, we have devised two different parameter inference schemes based on either Expectation-Maximization (EM)^13^ or variational inference (VI)^14^. In these frameworks, we assumed gene transcription during an induction phase (*t* < *t*_*s*_), followed by a repression phase (*t* > *t*_*s*_). Although this dynamical modeling paradigm has enabled the recovery of novel biological insight^13^, both models neglect gene-gene dependencies and assume a piecewise constant transcription rate for each gene, thereby overlooking the dynamic changes in the transcription rate itself. Biological systems fulfill these assumptions only approximately^24,119^, however, and exhibit more complex dynamics^14,16^, instead.

Upstream regulators govern gene transcription, a process neglected so far. To include these non-linear dynamics in our model of transcriptional dynamics, we define gene- and cell-specific transcription rates α_*gn*_ that we parametrize with a neural network α. In this setup, we assume that mature mRNA of upstream regulators and a gene-specific base transcription rate describe the transcription rate via

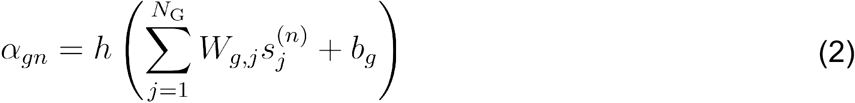

where is a nonlinear activation function, 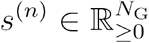 denotes the *n*-th sample, and *b*_*g*_ ∈ ℝ and 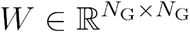 the base transcription rate and gene regulatory network (GRN) weight matrix, respectively.

The GRN weight matrix represents gene regulation, informed by a prior gene regulation graph *G*, curated from public datasets like cistrome^34^ or learned from single-cell, bulk epigenetic, or multi-ome sequencing data through GRN inference methods^23,24^. We represent the prior regulation graph *G* as a binary, skeleton matrix that encodes the interactions between transcripton factors (TFs) and target genes and provide two strategies for learning GRN structures from these priors: hard and soft constrained inference. In the case of hard constraints, *G* restricts the non-zero entries of the weight matrix to edges present in the prior graph, *i*.*e*.,

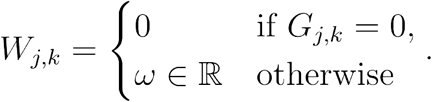

Conversely, for soft constraints, we allow learning new connections regularized by the prior graph *G* with

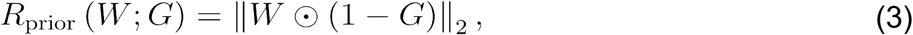

to favor TF-target connections present in the prior; ⊙ denotes the Hadamard product, *i*.*e*., element-wise product operation. The soft constraint, thus, allows for correcting missing links in the prior gene regulation graph.

In a realistic biological system, cells often undergo several internal orthogonal processes simultaneously like cell cycle versus differentiation, for example. In such cases, using a single latent time and a deterministic ODE system fails to model these biological processes jointly. Additionally, determining the number of biological processes and modeling them requires some prior information. Thus, similar to scVelo^13^ and veloVI^14^, we estimate gene-specific latent times *t*_g_ and aim to identify a unique latent time axis for each gene, combined with a global regulatory function to best describe the observed data.

The velocity function, defined by (1) and (2), governs cell state evolution, inducing a continuous vector field 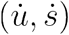. Given an initial condition (*u*_0_, *s*_0_), we simulate dynamics along each gene-specific latent time axis to estimate the integrated value for each gene and each cell as

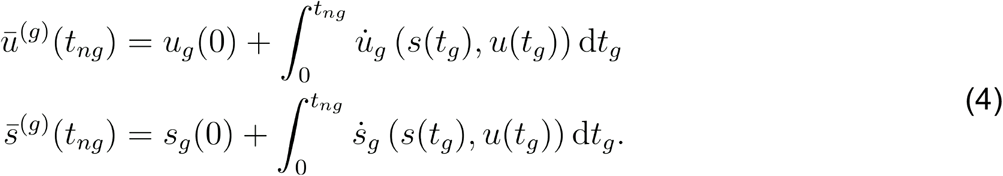

We simulate these state solutions for each gene efficiently through a parallelized numerical integration solver dopri5 implemented in the *torchode*^*37*^ Python package.

#### Model assumptions

We assume multiple time axes, representing different biological processes, but time-independent regulatory principles, *i*.*e*., a constant GRN weight matrix - a modeling choice that allows us to couple multiple gene dynamics. Similar to scVelo^13^ and veloVI^14^, we assume an initial gene expression of *u*(*t*_0_) = *s*(*t*_0_) = 0 at *t*_0_ = 0 and that each gene is on the same time scale. Here, we follow scVelo and veloVI and set *t*_max_ = 20. Lastly, we define the nonlinear activation function *h* in (2) as the softplus function

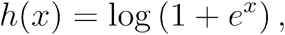

thus assuming that TFs predict the transcription rate of target genes based on additive effects.

### RegVelo generative process

As described above, we assume each gene has a specific latent time. We have already established in recent work that the inferred, gene-specific latent times have a low-rank structure, implying that we can generate the latent time matrix from multiple independent components that control cell dynamics^14^. Therefore, our model does not only couple genes through a shared regulatory principle encoded by *W* but also, like veloVI^14^, ties the gene-specific latent time via a local low-dimensional latent variable.

For each cell, we draw a low-dimensional latent variable from an isotropic Gaussian, *i*.*e*.,

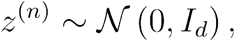

where *I*_*d*_ denotes the *d*-dimensional identity matrix. Here, we use *d* = 10 dimensions throughout this study. We define the gene-specific latent time as a function of this latent variable defined by

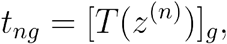

with 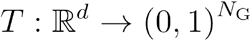 parametrized by a fully connected neural network. Utilizing the time information, we derive the state solution for each cell and gene through (4). Finally, we assume that the conditional normal distributions parametrized by θ

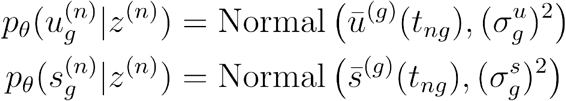

generates the observed data. Here, includes transcriptional regulation (*W*, *b*), kinetic parameters (β, *γ*), and neural network parameters.

Like veloVI, we consider the observed data {(*s*^(*n*)^, *u*^(*n*)^) |*n* ∈ {1,…, *N*_*C*_}} as nearest-neighbor smoothed expression data, which is also used as input for scVelo. RegVelo, like veloVI^14^, assumes that the smoothed spliced and unspliced abundances have been min-max scaled to the unit interval [0,1], independently for each gene. We also assume that the smoothed expression, representing an average of random variables, follows a sampling distribution centered on a mean value and is approximately normal.

### RegVelo inference

RegVelo aims to estimate the following parameters: (1) point estimates of kinetic parameters, including gene regulation weights summarized in the GRN matrix, base transcription, degradation, and splicing rate constants; (2) point estimates of the parameters of all neural networks; (3) a posterior distribution over the latent variable *z*, to model the underlying dynamic process. We use variational inference to approximate the posterior distribution and estimate all other parameters by maximizing the evidence lower bound (ELBO)^38^. Consequently, we model RNA velocity probabilistically by calculating it as a function of the variational posterior distribution of latent variables.

#### Variational posterior

We introduce the variational distribution *q*_ϕ_ parametrized by a neural network with parameter set ϕ to approximate the evidence function *log p*_ϕ_(*u*, *s*)

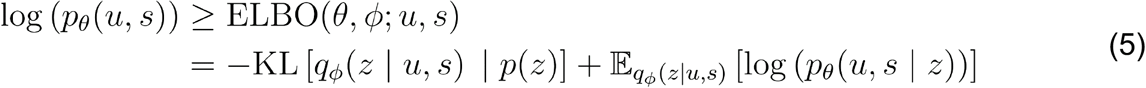

by maximizing the ELBO. We assume independent samples, allowing us to factorize the distribution *q*_ϕ_(*z*|*u*, *s*) and *p*_ϕ_(*u*, *s* |*z*) as

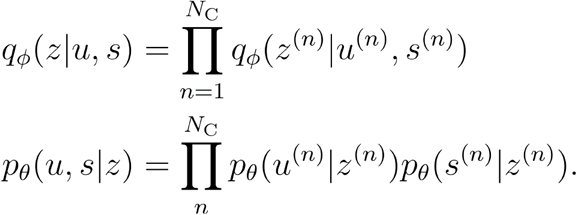

#### Dynamics regularization

RegVelo solves the dynamics through a numerical integrator, as the kinetic process is too complex to solve analytically. Avoiding poorly conditioned dynamics improves the accuracy of numerical integration^120^. Therefore, we add penalties on the regularity of the velocity function 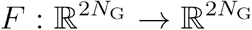 to ensure smoothness.

Here, we define the regularity as the total derivative of the velocity function with respect to time, *i*.*e*.,

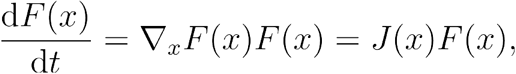

using the chain rule and with *x* = (*u*, *s*) and the Jacobian matrix *J*(*x*). To simplify the dynamics, we regularize the Jacobian matrix and velocity field to encourage small values: we use L2 regularization to penalize large entries of velocity entries, promoting a more regular vector field, and L1 regularization to penalize large entries in the Jacobian matrix to foster sparse gene regulation. Therefore, we define the regularization terms

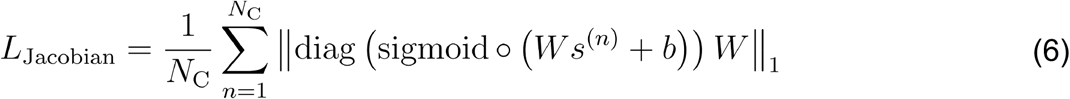

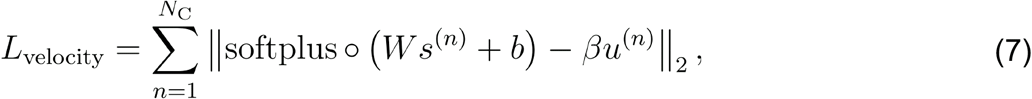

where, diag and sigmoid denote the operator to convert a vector into a diagonal matrix and the sigmoid activation function, respectively, and the operator to apply a scalar function component-wise. As the complexity of gene dynamics in our model stems primarily from gene regulation, we only regularize the Jacobian matrix and the unspliced RNA velocity to focus on aspects directly influenced by regulation.

Further, assuming that upstream regulators govern each gene transcription fully, we penalize large values of base transcription *b* with

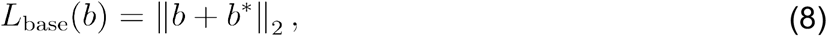

to keep the transcription rate near zero when upstream regulation effects diminish, *i*.*e*., *W s* = 0. We fixed *b*\* = 10 in this model. Finally, we define the dynamic regularization loss function as 0the sum of *L*_Jacobian_, *L*_velocity_ and *L*_base_.

#### RegVelo’s objective function

Combining (3), (5), (6), (7) and (8), our objective includes the three parts

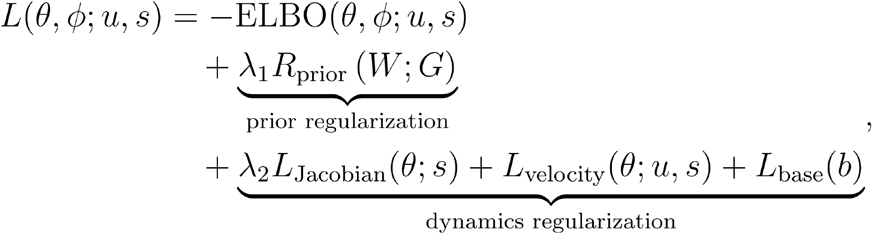

with tunable hyperparameters λ_1_ and λ_2_: λ_1_ controls the strengths of adding prior knowledge and λ_2_ governs the strengths of the L1 regularization of the Jacobian; increasing λ_2_ promotes greater sparsity in the learned GRN. By default, we set λ_1_ = 1 and λ_2_ = 0.

#### Network architecture and optimization

All neural networks are fully connected feedforward networks, with ReLU activation functions for hidden layers and softplus-transformations of the last layer to enforce non-negative kinetic parameters. We initialize the weight matrix *W* as a zero matrix, kinetic parameters, including splicing and degradation rates, as a constant values, and base transcription *b* and all neural network parameters through the default implementation in PyTorch (`torch.nn.init.xavier_uniform`_); following veloVI^14^, we initialize a unit splicing rate and estimate degradation based on a steady-state assumption. We use stochastic gradients along with the AdamW optimizer^121^ to optimize the loss function.

### Downstream tasks

#### Fitted abundance value

Let *u*\* (*n*) and *s*\*(*n*) be the random variable representing posterior predicted abundance values for unspliced and spliced RNA in cell, respectively. We denote their posterior distribution by *p*(*u*\*(*n*)|*u*^(*n*)^, *s*^(*n*)^) and *p*(*s*\*(*n*)|*u*^(*n*)^, *s*^(*n*)^) and define as

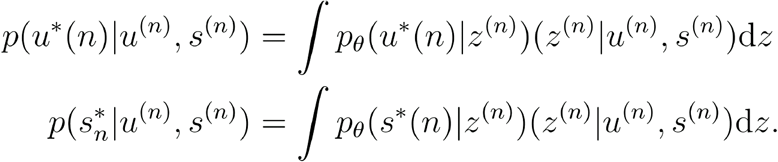

The fitted values for unspliced and spliced abundances are the means of these posterior distributions, *i*.*e*., 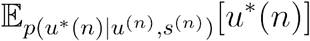 and 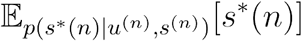, respectively. We estimate the mean value by sampling from the posterior distribution and taking the average.

#### Gene-wise latent time

We compute latent time for each gene and cell as 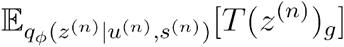 and estimate the expectation value by Monte Carlo sampling^122^, where we sample from the posterior distribution of *z*^(*n*)^ before calculating the average value of *T*(*z*^(*n*)^)_*g*_.

#### RNA velocity

We define the velocity of a specific gene in a particular cell as the function depending on the latent variable *z*^(*n*)^. To sample the velocity from the latent posterior distribution, we

1. Sample *z*^(*n*)^ from *q*_ϕ_(*z*^*(n)*^|*u*^*(n)*^, *s*^(*n*)^),
2. Calculate the latent time *t*_*ng*_ = *T*(*z*^*(n)*^)_*g*_,
3. Compute velocity in gene *g* and cell *n* using

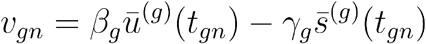

These steps provide velocity samples from the posterior predictive velocity distribution.

#### Velocity uncertainty estimation

As a variational inference-based model, veloVI^14^ defines an intrinsic uncertainty method to quantify uncertainty in each cell through sampling velocity vectors from the posterior distribution and calculate the variance of the cosine similarity of the samples; as an extension of veloVI, RegVelo follows the same approach.

#### Representing the underlying GRN

The gene regulation function (2) describes the regulation of unspliced RNA abundance of upstream regulators. We approximate the gene regulation function in the neighborhood of a specific *s*_*n*_ by first-order Taylor expansion

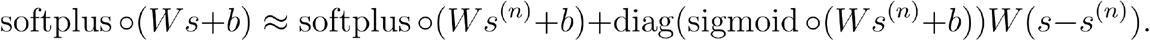

The coefficient of the first-order term denotes the strength of linear regulation. We define the GRN for cell *n* as

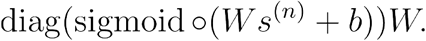

To define cell type-specific GRNs, let 𝒮(*c*) denote the set of cells of type *c*

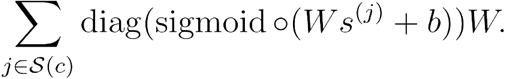

#### Simulation of perturbation effects

RegVelo generates cell dynamics based on gene regulation. By modifying the learned GRN in a trained RegVelo model, we simulate the in silico perturbed dynamics. Assuming we want to perturb the function of the TF *l* in a system, we mask the corresponding column of the weight matrix to generate a perturbed weight matrix

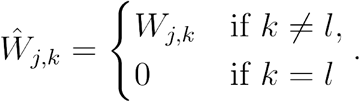

The perturbed weight matrix defines a new velocity function for numerical integration and induces a perturbed velocity *v*\*. We systematically compare the perturbed velocity with the original fitted velocity using the following metrics:

#### Local perturbation effects

We define the perturbation effects on each cell by computing the cosine dissimilarity

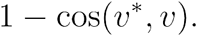

#### Cell fate perturbation effects

We quantify the perturbation effects on cell fate decisions by passing the different velocity estimates to the typical CellRank^15,16^ workflow: using the VelocityKernel in CellRank^15^ and assuming terminal states, we define the original cell fate probability matrix 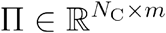 and perturbed probability matrix 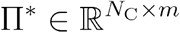, where each row represents a cell, and each column represents a terminal state. We can formulate the cell fate perturbation effects for the *k*-th terminal state as the *t*-test statistics

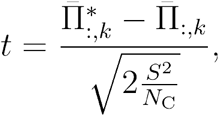

with sample means of fate probabilities toward the *k*-th terminal state 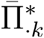 and 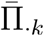, and the pooled variance *S*^2^ of the cell fate probabilities. To further quantify the depletion likelihood of each TF to the terminal state *k*, we used the normalized Mann-Whitney *U* statistic

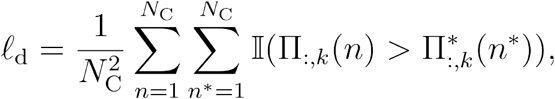

where 𝕀Π(*n*) > Π* (*n*\*))denotes the indicator function

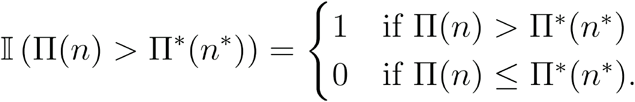

This statistic quantifies the probability that, in the predicted perturbation case, the cells will have lower cell fate probabilities toward the terminal state *k* than in the original unperturbed case. To intuitively compare the perturbation effect across lineages, we define the depletion score Δ_d_ by rescaling the depletion likelihood

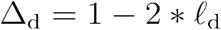

to indicate the perturbation effect when masking the TF regulon in the cell dynamics. Δ_d_ > 0 indicates enrichment effects on the *k*-th terminal state. Δ_d_ < 0 indicates depletion effects on the *k*-th terminal state.

### Benchmarking metrics

#### Velocity inference benchmark

We assess inferred velocities based on the average Pearson correlation between the ground truth and predicted gene velocity.

#### Latent time prediction benchmark

We define latent time of a cell as the mean value of its gene-specific latent times. Following, we calculate the Spearman correlation between predicted and ground truth cell latent times.

#### GRN inference benchmark

We benchmarked the GRN inference based on the area under the receiver operator characteristic curve (AUROC) using a set of ground truth TF-target connections *E*\* with scikit learn’s `roc_auc_score` function^123^. Specifically, we defined the ROC curve using the following procedure:

1. Rank all possible TF-target pairs (*j*, *k*) based on the absolute value of their predicted edge weight *W* _*j*, *k*_ in the inferred GRN.
2. For the TF-target pair (*j*, *k*), we define the ground truth label *y* (*j*, *k*) as

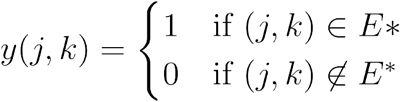
3. For thresholds τ ∈[τ_*min*_, τ_*max*_], we define the true positive and false positive rates based on the estimator

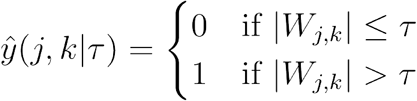

Here, we chose

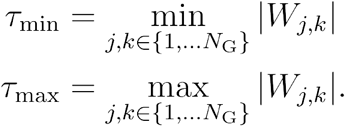

#### Parameter identifiability

Variational inference is sensitive to the parameter initialization because the ELBO is a nonconvex objective function^124^. We, thus, ensured the stability of estimates like the velocity, latent time, or gene regulatory network under different initialization by measuring the correlation of the estimates for each cell across multiple model runs. Here, we ran each probabilistic model overall five times and relied on Pearson correlation for all estimates except latent time where we computed the Spearman correlation to account for the ranked nature of the feature.

#### Velocity consistency

We expect the inferred vector field to be coherent in a uni-directional trajectory such as the cell cycle. We quantify a consistency score^13^ ζ for each cell *n* as the mean correlation of its velocity *v*(*n*) with its neighboring cell velocities, *i*.*e*.,

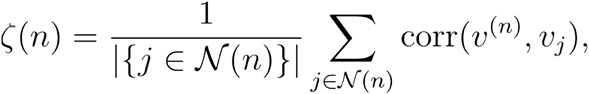

where 𝒩 (*n*) indicates the neighborhood of cell *n* and corr the Pearson correlation.

#### Cross-boundary correctness

Predicted cell state transitions need to align with ground truth cell state changes. Assuming we know the correct transition from a source cluster 𝒞_*A*_ to target cluster 𝒞_*B*_, we use the cross-boundary correctness (CBC) score^16,28^ to evaluate if our velocities align with the correct direction.

To compute the CBC score, we first identify the boundary between 𝒞_A_ as 𝒞_*B*_ as the set of all cells in 𝒞_*A*_ with at least one neighbor in 𝒞_*B*_

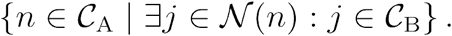

We then define the heuristic velocity 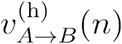 of observation as

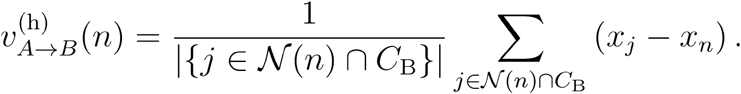

Following, we calculated the CBC score as the Pearson correlation between the inferred velocity *v*(*n*) and heuristic velocity 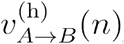. This score assesses the forward transition accuracy; we denote it as CBC ^(f)^ (*n*).

This definition of the CBC score only quantifies if the velocity of boundary 𝒞_A_ cells correctly points to their target state but fails to capture 𝒞_B_ observations pointing toward their ancestor state. To quantify this incorrect flow, we consider the heuristic velocity from 𝒞_B_ to 𝒞_A_ and defined the backward CBC score CBC ^(b)^ (*n*)as

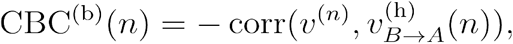

with Pearson correlation corr.

#### Terminal state identification

CellRank^15,16^ infers terminal states based on state change estimates combined with Markov chain theory. To benchmark different trajectory inference methods we rely on CellRank 2’s terminal state identification^16^ (TSI) score that quantifies how faithfully a method allows the recovery of terminal states with increasing number of macrostates that represent regions of the phenotypic manifold that cells are unlikely to leave. Given the ground-truth dynamics, CellRank recovers all terminal states of a given system and one terminal state for each macrostate considered.

We quantified TSI performance through the TSI score introduced by CellRank 2^16^. First, we used CellRank’s VelocityKernel *k* to estimate cell-cell transition probabilities based on the inferred RNA velocities and subsequently defined macrostates and identified terminal states. We represented the predicted terminal states as the macrostate with stability^15,16^ exceeding the threshold τ. Then, for each kernel matrix built upon the velocity estimates, we considered the step function *f*_*k*_ that maps the number of macrostates to the number of correctly predicted terminal states. Considering terminal states, an optimal method identifies *m* terminal states according to

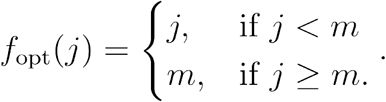

Based on these definitions, we calculated the TSI score as the area under the curve *f*_*k*_ relative to the area under the curve *f*_opt_. We calculated the TSI score for thresholds taken from 20 evenly spaced values of the unit interval, *i*.*e*., 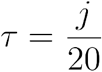, j ∈ {0,…,20}. Finally, we relied on a one-sided Welch’s t-test to identify the significantly best TSI performance.

#### Gene ranking score

Each approach for inferring cellular dynamics allows identifying and ranking potential lineage-associated genes of every lineage with CellRank^15,16^ or, to compare to competing approaches, by relying on least action path (LAP) analysis^22^. To assess this ranking, we curated a list of known lineage markers and regulators^125,126^ that an optimal method would rank the highest. Therefore, following the CellRank 2 method comparison workflow^16^, we evaluated the performance of each method as follows: first, consider a lineage, a set of known lineage-associated genes *D*, and a method *m*. Each method *m* assigns a ranking to a gene *g ∈ G*, denoted by r^(*m*)^(*g*), and, for each threshold T ∈ {1,…, | *G* |}, we counted how many lineage-associated genes rank among the top Tgenes with

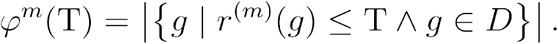

Next, to quantitatively evaluate the rankings, for each method *m*, we computed the area under the curve

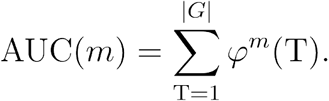

Since an optimal method ranks the lineage-associated genes highest, its AUC* is

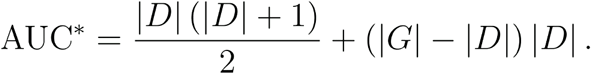

Finally, we evaluate the performance of methods relative to an optimal ranking through

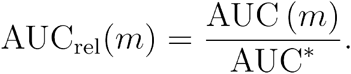

### Evaluating the impact of the GRN on dynamics inference

We developed two baseline models based on a pre-trained RegVelo model to assess the significance of GRNs in velocity inference. We created the first baseline, the *random GRN* model, by shuffling the estimated GRN from the pre-trained RegVelo model and used it as input for a new model. We retrained the original and new model but kept the GRN fixed. This approach allows assessing how the regulatory network affects the model’s cellular velocity and latent time estimates.

In the second baseline model, the *remove GRN* model, we removed all regulatory influences. As with the first baseline, we fixed the modified GRN and trained the model until convergence. This model served as a control to evaluate the impact of entirely omitting regulatory information on velocity inference.

We quantified the performance of these baseline models compared to the original RegVelo model by calculating the CBC score and the Spearman correlation of latent time estimates with ground truth. To gain statistical significance, we repeated this process 30 times for each model and used the Wilcoxon signed-rank test to compare the original model with each baseline, *i*.*e*., original vs. random GRN and original vs. removed GRN. This evaluation allowed us to determine the critical role that GRNs play in accurately modeling cellular dynamics.

#### Robustness to prior GRN estimates

RegVelo relies on a prior gene regulatory graph, and since this prior knowledge is usually inaccurate, the method must be robust to such imperfections. To assess the robustness of our method to inaccuracies in prior gene regulatory graphs, we utilized dyngen-simulated datasets^127^ as described in the Datasets section. Following, we generated corrupted prior gene regulatory graphs by randomly replacing different fractions of existing gene regulations with nonexistent ones based on the original gene regulation graph used in the dyngen simulations.

We simulated five levels of corrupted fractions {0.2, 0.4, 0.6, 0.8, 1.0}, each generating five different corrupted prior graphs to account for corruption randomness. We ran RegVelo with each of these corrupted prior graphs and evaluated performance based on the AUROC score of the inferred GRN, and velocity and latent time correlation. We used these metrics to determine how well RegVelo maintained accuracy and reliability despite the corruption in the prior knowledge.

### Quantifying perturbation effects

#### Dynamo-based perturbation prediction

To perform perturbation prediction with dynamo^22^, we followed the corresponding tutorial provided in the Python package’s documentation. In the first step, dynamo estimates cell velocities; we used `dynamo.tl.dynamics` to estimate kinetic rates and used `dynamo.tl.reduceDimension` to perform the dimensional reduction. Next, we used `dynamo.tl.cell_velocities` with argument `basis=‘pca’` to project cell velocities to the PCA space. We further learned a vector field function based on this projection with `dynamo.vf.VectorField` with argument `basis=‘pca’`.

Dynamo provides two methods to simulate perturbation effects: *KO* or *expression perturbation*. For *KO*-based perturbation simulation, the method directly removes the regulation of the TF through masking the Jacobian matrix and generates a perturbed vector field. Following our CellRank-based quantification of cell fate probabilities, we first used the dynamo-inferred velocities in the typical CellRank workflow. We further used `dynamo.pd.KO` with argument `store_vf_ko=True` to predict the velocities after perturbing a certain TF and quantified the corresponding perturbed cell fate probabilities. Finally, we used Pearson correlation to assess the change in cell fate probabilities towards each lineage to quantify the perturbation score.

For *expression perturbation-*based simulations, dynamo estimates perturbation effects by simulating perturbed gene expression and calculating the perturbation shift matrix, corresponding to the difference between the perturbed and the original gene expression. We computed the perturbation shift matrix on each cell through `dynamo.pd.perturbation` and quantified the perturbation score on each cell as the inner product between the perturbation shift and velocity matrices. Finally, we calculated the average perturbation score within each terminal cell type.

#### CellOracle-based perturbation prediction

To perform perturbation prediction with CellOracle^25^, we followed the tool’s tutorial: after defining the oracle object with `oracle = celloracle.Oracle()`, we injected the base gene regulatory graph via `oracle.import_TF_data`, calculated the PCA representation through `oracle.perform_PCA` and performed gene expression imputation with `oracle.knn_imputation`. Next, we estimated a GRN for each cell type with `oracle.get_links` and set the regularization parameter of Bayesian ridge regression `alpha=10` to prevent overfitting during GRN inference. Finally, we filtered edges of the GRNs with `links.filter_links` and fitted the GRNs for perturbation simulation by `oracle.fit_GRN_for_simulation`.

To perform TF perturbation simulations with CellOracle, the method requires a pseudotime-based reference vector field defined in 2D space. We used `Gradient_calculator` to calculate the reference gradient based on a pseudotime or latent time inferred by velocity models. We then defined grid points with `gradient.calculate_p_mass` and `gradient.calculate_mass_filter` and projected the reference gradient onto the grid points using `gradient.calculate_gradient`. For each TF, `oracle.simulate_shift` simulated the shift vector, and `oracle.estimate_transition_prob` estimated the cell transition probability matrix; we relied on `n_neighbors=30` neighbors and calculated the shift matrix in PCA space with `oracle.calculate_embedding_shift`. Next, we calculated the inner product between the perturbed shift vectors and reference velocity vectors in 2D space with `calculate_inner_product` and `calculate_digitized_ip`. Finally, we calculated the perturbation score within each lineage using `get_negative_PS_p_value`.

### Datasets

#### General processing

We conducted all analyses using Scanpy^128^ and scVelo^13^, using default parameters unless specified otherwise. We applied scVelo to filter out genes expressed in fewer than ten cells in the spliced and unspliced counts before normalizing their gene expression. If not stated otherwise, we selected the top 1000 highly variable genes (HVGs) based on gene dispersion parameters. We performed these filtering and normalization steps using `scvelo.pp.filter_and_normalize`. Next, we computed the top 30 PCA components and constructed a nearest-neighbor graph with 30 neighbors using `scanpy.tl.pca` and `scanpy.pp.neighbors`, respectively. We then calculated first-order moment matrices as smoothed spliced and unspliced gene expression on the nearest neighbor graph using `scvelo.pp.moments`. All velocity models take these moment matrices as input.

We applied gene-wise min-max scaling to preprocess both spliced and unspliced counts. Following the preprocessing employed by scVelo^13^ and veloVI^14^, we applied scVelo’s deterministic model and kept genes with a positive coefficient of determination *R*^2^ > 0 for further RNA velocity inference because RegVelo’s spliced equation with positive kinetic parameters *β* and *γ* implies a correlation between spliced and unspliced read counts; negative *R*^2^ values indicate poor model fit, suggesting the gene may not be informative for dynamical analysis. However, even if some transcription factors (TFs) do not meet the velocity gene selection criteria, their regulatory functions remain crucial for understanding broader gene regulation during differentiation. Therefore, we retained all TFs contained in the highly variable gene set unless specified otherwise.

We downloaded lists of TFs for humans and mice from the cisTarget database of the SCENIC software^17,23^, and for zebrafish, we retrieved the corresponding information from the AnimalTFDB v4.0 database^129^. Finally, if not specified otherwise, we used the *soft constraint* mode to allow RegVelo to learn new edges in the GRN.

#### Trajectory inference benchmark

We inputted all velocity estimations into the typical CellRank^15,16^ workflow for downstream analyses. To compare velocity estimations, we used only CellRank’s VelocityKernel to compute the transition matrix for macrostates estimation. Otherwise, we combined the VelocityKernel and ConnectivityKernel with weights 0.8 and 0.2, respectively, as shown in the CellRank tutorials. To benchmark trajectory inference performance, we used the CBC score provided by the kernel method `cbc` and the TSI score supplied by the GPCCA estimator `tsi`^16^; the Benchmarking metrics section provides the relevant details.

#### Driver gene analysis

For each velocity-based method, we rely on CellRank to infer lineage-associated genes, following the tutorials for the VelocityKernel. Further, we computed terminal states and their corresponding fate probabilities using the GPCCA estimator. We used the `compute_lineage_drivers` function to identify genes that correlate with specific lineages.

To identify putative cell cycle activation-related drivers, we utilized cell cycle markers to compute cell cycle scores CCS for the G1, G2/M, and S phases by applying `scvelo.tl.score_genes_cell_cycle`. We defined the cell cycle score through

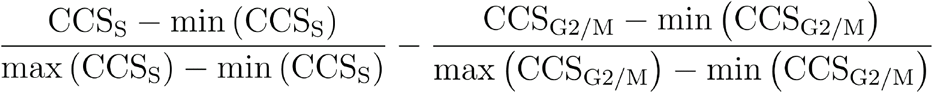

Finally, we calculated Pearson correlations between the expression levels of all TFs and this score to identify potential cell cycle regulators.

#### Perturbation screening

We simulated regulon knock-outs for each TF to identify lineage drivers through perturbation screening and estimated the resulting perturbed vector fields. After obtaining these perturbed velocities, we incorporated the perturbed and original velocities into the canonical CellRank workflow.

We used the GPCCA estimator^130^ to calculate fate probabilities and compare these probabilities with Pearson correlation, allowing us to quantify the effect of each perturbation on the lineages. To ensure the robustness of these perturbation predictions, we ran RegVelo five times, each with a different initialization for the perturbation simulations. Finally, after assessing the effects of perturbation on each gene and terminal state, we averaged the results from all five runs to obtain the final prediction.

#### Benchmarking related methods

We followed the provided tutorials for each velocity method^13,14,28,30,31^ to infer cell velocities. For scVelo, we first used `scvelo.tl.recover_dynamics` to estimate kinetic parameters and applied `scvelo.tl.velocity` with argument `mode=“dynamical”` to calculate velocities. For UniTVelo, we estimated velocities with `unitvelo.run_model` with the argument `FIT_OPTION=“1”`. In the case of VeloVAE, we initialized the model using `velovae.VAE` with argument `dim_z=10`, then used `velovae.train` to estimate velocities. We also ran a full Bayesian VeloVAE model with rate prior set as `‘alpha’: (0, 1.0), ‘beta’: (0, 0.5); ‘gamma’: (0, 0.5)`, and initialized the model with additional argument `full_vb=True`.

For TFVelo, we estimated the velocities with `TFvelo.tl.recover_dynamics`. We then extracted the estimated GRN from `adata.varm[‘fit_weights_final’]`, and used this GRN for the subsequent GRN benchmark. For cell2fate, we first applied `cell2fate.utils.get_max_modules` to estimate the number of modules, and estimated velocities through a `cell2fate.Cell2fate_DynamicalModel` model, trained with the `train()` method.

We followed the provided tutorials for each GRN inference method^25,31,46,131^. For CellOracle, we used the `fit_All_genes` function to compute the GRN, and for GRNBoost2, we used the `arboreto.algo.grnboost2` function to calculate the GRN. Importantly, GRNBoost2 does not estimate gene self-regulation but Pearson correlation always assigns self-regulation to every gene. To make the approach comparable to our method, we followed a previous GRN inference benchmark^132^ and removed all edges corresponding to self-regulation from the predicted and ground truth GRNs when calculating the AUROC score. Since spliceJAC requires pre-defined cell cluster information as input, we first ran `scanpy.tl.leiden` to find cell clusters, before applying `splicejac.tl.estimate_jacobian` for GRN estimation.

#### Cell cycling dataset

After training the RegVelo model and estimating velocities, we calculated the CBC score with the ground truth differentiation directions, including G1 phase to S phase and S phase to G2/M phase; the original studies provided the phase label information.

We calculated the velocity consistency score using `scvelo.tl.velocity_confidence`. To further validate our results, we used the fluorescent ubiquitination-based cell-cycle indicator (FUCCI) derived score^39^ as the proximal ground truth of cell cycling time and correlated it with inferred cell latent time using Spearman correlation.

#### Evaluating GRN inference

To evaluate GRN predictions through our AUROC score, we first defined the proximal ground truth GRN for the regulators by retrieving ground truth edges from ChIP-Atlas^133^, this resource provides putative binding scores of TF-target pairs derived from ChIP-seq datasets across different cell lines. We set the links with a MACS2^134,135^ score greater than 500 as the proximal ground truth links; this stringent significance threshold, which corresponds to a q-value less than 10^−5045,133^, ensures that our defined proximal ground truth links are reliable.

To calculate the AUROC score for each TF, we followed our pipeline outlined in the Bechnmark metrics section. To provide a comprehensive evaluation, we used the mean AUROC score across TFs with more than 50 targets in the cycling dataset to benchmark the GRN prediction.

#### Pancreatic endocrinogenesis

To infer the gene regulatory graph as prior knowledge for RegVelo, we began by utilizing the 10x multiome pancreatic endocrine datasets^48^ of the same mouse line (Ngn3-Venus fusion) and the same embryonic stages (E14.5 and E15.5) to the scRNA-seq dataset used in the scVelo study^47^. We then applied Pando^24^ to these 10x multiome datasets to infer the GRN with its default preprocessing and parameter settings.

#### Prior GRN definition

Pando’s GRN inference consists of two steps: first, it scans for motifs in candidate cis-regulatory element (CRE) regions on the genome with the `find_motifs` function. Second, it infers the GRN by integrating gene expression and peak accessible information using the `infer_grn` function. The first step requires TF-motif binding information and position weight matrices (PSM). Here, we used the mouse_pwms_v2 dataset, which collects mouse motifs from the CIS-BP database and corresponding PSMs, from the R package chromVARmotifs^136^ to provide this information. We inferred the prior GRN estimate through XGBoost^137^ and used Signac^138^ to link peaks with genes during the second step. We binarized the constructed GRN, *i*.*e*., set weights of nonzero edges to 1 and 0 otherwise; we used this binary graph as the prior gene regulatory graph for RegVelo.

#### Downstream analysis

We inputted the estimated velocity into the downstream CellRank workflow, used eight macrostates, and defined “Alpha”, “Beta”, “Delta”, and “Epsilon” as the terminal states.

#### Epsilon cell substate identification

We defined sub-clusters in Epsilon cells by first subsetting to the annotated Epsilon cells from the pancreatic endocrine dataset. We then computed PCA embedding of this subset and constructed a neighborhood graph using 30 PCs and 30 neighbors with `scanpy.tl.pca` and `scanpy.pp.neighbors` functions. Next, we identified the sub-clusters using the Leiden algorithm (`scanpy.tl.leiden`). We defined the sub-cluster with the highest average alpha cell fate probabilities as State A (Supplementary Fig. 2b). To further characterize both sub-clusters, we used the Wilcoxon rank-sum test to identify differentially expressed genes.

#### Human hematopoiesis

We conducted all analyses on the dataset preprocessed by the original study^22^. All other preprocessing followed our standard preprocessing pipeline.

#### Prior GRN inference

We curated prior gene regulatory graphs from two sources: putative gene regulation based on multi-omics datasets and experimental ChIP-seq datasets. We used a publicly available putative GRN inferred by the multiome GRN inference model *Dictys* on multi-omics bone marrow mononuclear cells^26^. Since this putative GRN may be incomplete, we also curated a GRN from human hematopoiesis ChIP-seq datasets of the UniBind database^35,71^. By merging both GRNs, we created a prior gene regulatory graph for RegVelo input.

We ran RegVelo with *soft constraints*, with unregularized (λ_2_ = 0) and regularized (λ_2_ = 1) settings for comparison. We inputted the estimated velocity into the downstream CellRank workflow to compute seven macrostates and defined “Mon”, “Bas”, “Ery”, “Meg” and “Neu” as the terminal states.

#### Method comparison

We compared all velocity estimations with generic trajectory inference benchmark procedures, using the PCA representation as the input to calculate the CBC score. We calculated the forward CBC score and backward CBC score for each method by the kernel method `cbc`, with ground-truth transitions: “HSC” to “GMP-like”, “HSC” to “MEP-like”, “GMP-like” to “Mon”, “MEP-like” to “Ery”, “MEP-like” to “Meg” and “MEP-like” to “Bas”. We further used Welch’s *t*-test to assess if the log ratio of CBC score of the regularized RegVelo model and the other model including unregularized RegVelo, scVelo, and veloVI models was significantly higher than zero.

To compare how RegVelo and veloVI predicted uncertainty in terminal states of the human hematopoiesis dataset, we first calculated the median uncertainty in progenitor cells (HSC, MEP-like, and GMP-like) and normalized the terminal state uncertainties based on this median. We then calculated the log_2_ ratio of the normalized uncertainties between veloVI and RegVelo. We applied Welch’s *t*-test to assess whether veloVI’s uncertainty was significantly higher than RegVelo’s in terminal states.

To evaluate the estimated gene rankings, for each system that we analyzed, we manually compiled a list of known driver TFs from the literature for each lineage (Supplementary Tables 1 and 2). For lineage-associated genes, we curated genes from CellMarker 2.0^126^ and other relevant literature^125^.

#### Dynamo-based analysis

To predict the drivers using dynamo, we strictly followed the analysis pipeline provided by the tool’s tutorial^22^. We calculated moment matrices using dynamo-inferred neighbor graph and group `label=“time”`. Following, we used the dynamo function `dynamo.tl.dynamics` to estimate cell velocities with arguments `group=“time”`, `one_shot_method=“sci_fate”`, `model=“deterministic”`.

With the stable fixed points given by the dynamo tutorials, we identified putative lineage drivers using dynamo’s least action path analysis. Firstly, we projected the dynamo velocity field onto the PCA space with the function `dynamo.tl.cell_velocities` with `basis=“pca”` and learned a vector field function based on this projection via the function `dynamo.vf.VectorField` with `basis=“pca”`. Next, we defined the terminal states as the 30 nearest neighbors computed in UMAP space of each stable fixed point using the function `dynamo.tools.utils.nearest_neighbors`. We computed the least actions path starting from each cell in the HSC cluster to each cell in the clusters labeled “Mon” or “Ery” with `dynamo.pd.least_action`. We further identified the genes correlated with the path using `dynamo.pd.GeneTrajectory` and ranked the lineage-associated genes with `dynamo.vf.rank_genes`.

#### Zebrafish neural crest development

##### Fish husbandry and Smart-seq3 single-cell RNA-Seq

Animal experiments complied with UK Home Office protocols under the Animals (Scientific Procedures) Act 1986, and followed the Guide for the Care and Use of Laboratory Animals^139^. All vertebrate animal research was conducted at Oxford University Biomedical Services. Adult fish care followed established protocols, and embryos were staged using standard methods under an Olympus dissecting stereomicroscope^140^.

We generated Smart-seq3 data to complement the 10x multiome data used to recontrust the cranial NC GRN^82^. Embryos were harvested by crossing zebrafish lines Gt(foxd3-mCherry)^ct110R^ and Gt(foxd3-Citrine)ct110^141^, incubating to the desired stages, and dissociating with Liberase TM (Roche) for 25 minutes at 32°C. Cell suspension was filtered through 100-µm strainers (PluriSelect), and stained with 1-3 µg/ml DAPI. Live Citrine-positive (Citrine+Cherry-) and double-positive (Citrine+Cherry+) single cells were isolated using fluorescence-activated cell sorting (FACS) on a Sony MA900 instrument.

To generate full-transcript-length, deep scRNA-Seq data, we prepared the Smart-seq3 libraries following the protocol^81^. During FACS, cells were sorted into individual wells containing 3-µl lysis buffer dispensed by the MANTIS liquid handler (FORMULATRIX). The lysis buffer consisted of 5% Polyethylene Glycol 8000 (Sigma), 0.1% Triton X-100 (Sigma), 0.5 U/µl RNAse inhibitor (Takara), 0.5 µM OligoT_30_VN (IDT), 0.5 mM/each dNTPs (Thermo Fisher) and nuclease-free water (Thermo Fisher). Note that these concentrations reflect their concentrations in the 4-µl reverse transcription reaction. The plates were briefly centrifuged, quickly frozen on dry ice and stored at −80°C. In brief, the plates of sorted cells were transferred on dry ice from −80°C, incubated at 72°C for 10 min, and cooled down to 4°C. Quickly, 1 µl/well reverse transcription master mix containing 25 mM Tris-HCl pH8 (Invitrogen), 30 mM NaCl (Invitrogen), 2.5 mM MgCl_2_ (Invitrogen), 1 mM GTP (Thermo Fisher), 8 mM DTT (Thermo Fisher), 0.5 U/µl RNAse inhibitor (Takara), 2 µM TSO (IDT), 2 U/µl Maxima H-minus (Life Technologies) was added. The reverse transcription program was 42°C 90 min, 10x (50°C 2 min, 42°C 2 min), 85°C 5 min and 4°C hold. MANTIS added 6 µl/well PCR mix containing 1x KAPA HiFi HotStart buffer, 0.3 mM/each dNTPs, 0.5 mM MgCl_2_, 0.5 µM forward primer, 0.1 µM reverse primer and 1 U/µl polymerase (Roche). The PCR program was 98°C 3 min, 23x (98°C 20 s, 65°C 30 s, 72°C 4 min), 85°C 5 min and 4°C hold. These cDNA samples were cleaned up by 0.6:1 AMPure XP beads and relocated into 384-well plates using FxP (Beckman Coulter), quantified by Quant-iT PicoGreen (Invitrogen) on CLARIOstar (BMG Labtech), quality checked by TapeStation 4200 and High Sensitivity D5000 assays (Agilent) and normalized to 0.3 ng/µl using MANTIS. Next, 150 pg/µl cDNA was tagmented by adding 0.5 µl Smart-seq3 in-house Tagmentation buffer (40 mM Tris-HCl pH7.5, 20 mM MgCl_2_, 20% Dimethylformamide and UltraPure water in 4x buffer), 0.08 µl Amplicon Tagmentation Mix (Illumina) and UltraPure water for a total reaction volume of 2 µl and incubating at 55°C for 10 min. The tagmentation was stopped by adding 0.5 μl of 0.2 % SDS and incubated at room temperature for 5 min. Then 1.5 µl Nextera index primer pairs and 3 µl PCR master mix (1x Phusion HF buffer, 0.2 mM/each dNTPs, 2 U/µl Phusion HF) were added to each well of the tagmentation product and the PCR program was 72°C 3 min, 98°C 3 min, 12x (98°C 10 s, 55°C 30 s, 72°C 30 s), 72°C 5 min and 4°C hold. The libraries were pooled, cleaned up by 0.6:1 AMPure XP beads and sequenced on NovaSeq for 150-bp paired end reads (Illumina).

##### Zebrafish Smart-seq3 data analysis

To preprocess our Smart-seq3 data, fastq files for each Smart-seq3 pool library were merged into read 1, read 2, index 1, and index 2 four fastq files. Next, we applied zUMIs^142^ to the fastq files with the expected barcodes file containing the Nextera v2 index combinations, the GRCz11 Ensembl 105 reference genome built by STAR v2.7.3a ^143^ and the parameters including barcode binning of 1, no automatic barcode detection, and the minimum number of reads per cell of 0.

##### Dynamic inference via RegVelo

We used the ultra-deep full-length Smart-seq3 scRNA-seq dataset during neural crest cell development for RegVelo implementation. We first filtered out genes expressing fewer than ten cells in spliced and unspliced matrices. To fully model the transcription regulation dynamics, we selected the top 8,000 genes for further downstream analysis.

We utilized the neural crest GRN^82^ built by SCENIC+ using the 10x multiome datasets as the input for RegVelo. After training models, we inputted inferred velocity estimations into the CellRank workflow, proceeded with eight macrostates and defined “facial mesenchyme”, “pharyngeal arch 2”, “mNC hox34” and “pigment” as the terminal states in this system.

##### Driver gene prediction benchmarking

Besides our canonical lineage driver identification through TF expression via CellRank^16^, we also correlated TF activity with cell fate probabilities to rank TFs.

To quantify TF activity in zebrafish datasets with SCENIC+’s inferred TF regulon, we used AUCell^17^ to score each TF-regulon activity within each cell through `AUCell_calcAUC.` Following, we calculated the correlation of each regulon activity with cell fate probabilities and ranked the genes according to their correlation coefficients within each lineage. Finally, we evaluated the identified putative drivers in mNC head mesenchymal and Pigment lineage using classical drivers as ground truth (Supplementary Table 1).

##### Perturbation effects prediction and benchmarking

We applied dynamo and CellOracle on the zebrafish dataset following the generic steps described in the Quantifying perturbation effect section. When incorporating dynamo inferred velocities into the CellRank workflow, we computed two transition matrices using the VelocityKernel and ConnectivityKernel and combined them with 0.8 and 0.2 weights, respectively; we computed eight macrostates and defined “facial mesenchyme”, “pharyngeal arch 2”, “mNC hox34” and “pigment” as the terminal states.

The CellOracle perturbation analysis required information on a base GRN and pseudotime; we used the SCENIC+-inferred GRN^82^ as the base GRN structure. For pseudotime estimates, we estimated pseudotime with scVelo in an unsupervised manner, using `scvelo.tl.recover_dynamics` and `scvelo.tl.velocity` to estimate kinetics, followed by `scvelo.tl.latent_time` to estimate the latent time for each cell.

##### In vivo Perturb-seq of zebrafish neural crest development

To systematically validate RegVelo’s predicted perturbation effects, we implemented CRISPR/Cas9 knockouts and Perturb-seq as previously described^82^. Our Perturb-seq dataset on zebrafish neural crest development included knockouts of seven regulons: *tfec, mitfa, bhlhe40, tfeb, elf2, nr2f2*, and *nr2f5*. For benchmarking, we combined this dataset with an additional one^82^, resulting in 22 conditions: 14 single-gene, 5 two-gene, and 3 multi-gene knockouts. We generated F_0_ knockout crispants by injecting the mixture of 250 ng/µl single guide RNAs (sgRNAs), 3.36 µM EnGen Cas9, and 0.05% Phenol red into 1-cell-stage embryos from crosses of *Gt(foxd3-Citrine)*^*ct110*^ and wild-type lines. At the 21-somite stage, foxd3-Citrine-positive cells were isolated by flow cytometry. Gene expression and CRISPR libraries were prepared according to the CG000510 Chromium NextGEM Single Cell 5’ v2 CRISPR User Guide (Rev B, 10x Genomics) and sequenced on NextSeq and NovaSeq platforms (Illumina).

##### Quantification of in vivo perturbation effects

The zebrafish Perturb-seq data were processed following an established workflow^82^. Briefly, raw data were preprocessed with CellRanger using SC5P-R2 chemistry, aligning reads to the reference genome that incorporated mCherry and Citrine genes along with GRCz11 Ensembl105. Cellular genotypes were assigned based on sgRNA counts, with ambient RNA contamination and doublets removed.

Cell clustering was performed using Seurat, and cell state annotations were enhanced by integrating multiome-gene expression (GEX) data with reciprocal PCA (RPCA)^144^. Neural crest clusters were isolated for further clustering and visualized using the PHATE method^145^. Perturbation effects were quantified using MELD^146^ by comparing each knockout condition to the non-targeting control. Latent time was inferred from the multiome dataset^82^ via RPCA and averaging latent time of neighboring cells. Within specific lineages, cells were categorized into early and late stages based on median latent time. MELD then calculated depletion-enrichment likelihoods on the 0-1 scale, with late-stage cells’ average likelihood providing a measure of short-term lineage-specific effects for each knockout.

##### TF binding site prediction

To predict the TF binding sites, we used the interpretation of previously trained deep learning models^82^ by ChromBPNet^100^, de novo motif discovery results^82^ predicted by TFmodisco-lite^100^ and hit calling results^82^ by Finemo^100^. For visualization, we used Signac^138^ to plot the pseudobulk chromatin accessibility tracks of the multiome ATAC data, and used the WashU Epigenome Browser^147^ to visualize the ChromBPNet profile contribution scores, which indicated the base-resolution importance of DNA sequences for chromatin accessibility.

##### Whole-mount hybridization chain reaction (HCR) and phenotypic check

To visualize the mRNA in situ, we applied HCR^101^ targeting *elf1* mRNA in zebrafish embryos by following the ‘In situ HCR v3.0 protocol’ for whole-mount zebrafish larvae from Molecular Instruments. Hybridized embryos were embedded in 1% low-melting point agarose and imaged with confocal microscopes (Zeiss). To check the phenotypes of wild-type and knockout embryos, we performed live imaging with MVX10 stereomicroscope (Olympus).

## References

1. Pijuan-Sala, B., Griffiths, J.A., Guibentif, C., Hiscock, T.W., Jawaid, W., Calero-Nieto, F.J., Mulas, C., Ibarra-Soria, X., Tyser, R.C.V., Ho, D.L.L., et al. (2019). A single-cell molecular map of mouse gastrulation and early organogenesis. Nature 566, 490–495.

2. Vento-Tormo, R., Efremova, M., Botting, R.A., Turco, M.Y., Vento-Tormo, M., Meyer, K.B., Park, J.-E., Stephenson, E., Polanski, K., Goncalves, A., et al. (2018). Single-cell reconstruction of the early maternal-fetal interface in humans. Nature 563, 347–353.

3. Litvinuková, M., Talavera-López, C., Maatz, H., Reichart, D., Worth, C.L., Lindberg, E.L., Kanda, M., Polanski, K., Heinig, M., Lee, M., et al. (2020). Cells of the adult human heart. Nature 588, 466–472.

4. Waddington, C.H. (2014). The Strategy of the Genes 1st Edition. (Routledge).

5. Saelens, W., Cannoodt, R., Todorov, H., and Saeys, Y. (2019). A comparison of single-cell trajectory inference methods. Nat. Biotechnol. 37, 547–554.

6. Wolf, F.A., Hamey, F.K., Plass, M., Solana, J., Dahlin, J.S., Göttgens, B., Rajewsky, N., Simon, L., and Theis, F.J. (2019). PAGA: graph abstraction reconciles clustering with trajectory inference through a topology preserving map of single cells. Genome Biol. 20, 59.

7. Haghverdi, L., Büttner, M., Wolf, F.A., Buettner, F., and Theis, F.J. (2016). Diffusion pseudotime robustly reconstructs lineage branching. Nat. Methods 13, 845–848.

8. Bendall, S.C., Davis, K.L., Amir, E.-A.D., Tadmor, M.D., Simonds, E.F., Chen, T.J., Shenfeld, D.K., Nolan, G.P., and Pe’er, D. (2014). Single-cell trajectory detection uncovers progression and regulatory coordination in human B cell development. Cell 157, 714–725.

9. Qiu, X., Mao, Q., Tang, Y., Wang, L., Chawla, R., Pliner, H.A., and Trapnell, C. (2017). Reversed graph embedding resolves complex single-cell trajectories. Nat. Methods 14, 979–982.

10. Setty, M., Kiseliovas, V., Levine, J., Gayoso, A., Mazutis, L., and Pe’er, D. (2019). Characterization of cell fate probabilities in single-cell data with Palantir. Nat. Biotechnol. 37, 451–460.

11. Street, K., Risso, D., Fletcher, R.B., Das, D., Ngai, J., Yosef, N., Purdom, E., and Dudoit, S. (2018). Slingshot: cell lineage and pseudotime inference for single-cell transcriptomics. BMC Genomics 19, 477.

12. La Manno, G., Soldatov, R., Zeisel, A., Braun, E., Hochgerner, H., Petukhov, V., Lidschreiber, K., Kastriti, M.E., Lönnerberg, P., Furlan, A., et al. (2018). RNA velocity of single cells. Nature 560, 494–498.

13. Bergen, V., Lange, M., Peidli, S., Wolf, F.A., and Theis, F.J. (2020). Generalizing RNA velocity to transient cell states through dynamical modeling. Nat. Biotechnol. 38, 1408– 1414.

14. Gayoso, A., Weiler, P., Lotfollahi, M., Klein, D., Hong, J., Streets, A., Theis, F.J., and Yosef, N. (2024). Deep generative modeling of transcriptional dynamics for RNA velocity analysis in single cells. Nat. Methods 21, 50–59.

15. Lange, M., Bergen, V., Klein, M., Setty, M., Reuter, B., Bakhti, M., Lickert, H., Ansari, M., Schniering, J., Schiller, H.B., et al. (2022). CellRank for directed single-cell fate mapping. Nat. Methods 19, 159–170.

16. Weiler, P., Lange, M., Klein, M., Pe’er, D., and Theis, F. (2024). CellRank 2: unified fate mapping in multiview single-cell data. Nat. Methods. 10.1038/s41592-024-02303-9.

17. Aibar, S., González-Blas, C.B., Moerman, T., Huynh-Thu, V.A., Imrichova, H., Hulselmans, G., Rambow, F., Marine, J.-C., Geurts, P., Aerts, J., et al. (2017). SCENIC: single-cell regulatory network inference and clustering. Nat. Methods 14, 1083–1086.

18. Matsumoto, H., Kiryu, H., Furusawa, C., Ko, M.S.H., Ko, S.B.H., Gouda, N., Hayashi, T., and Nikaido, I. (2017). SCODE: an efficient regulatory network inference algorithm from single-cell RNA-Seq during differentiation. Bioinformatics 33, 2314–2321.

19. Specht, A.T., and Li, J. (2017). LEAP: constructing gene co-expression networks for single-cell RNA-sequencing data using pseudotime ordering. Bioinformatics 33, 764–766.

20. Chan, T.E., Stumpf, M.P.H., and Babtie, A.C. (2017). Gene Regulatory Network Inference from Single-Cell Data Using Multivariate Information Measures. Cell Syst 5, 251–267.e3.

21. Qiu, X., Rahimzamani, A., Wang, L., Ren, B., Mao, Q., Durham, T., McFaline-Figueroa, J.L., Saunders, L., Trapnell, C., and Kannan, S. (2020). Inferring Causal Gene Regulatory Networks from Coupled Single-Cell Expression Dynamics Using Scribe. Cell Syst 10, 265–274.e11.

22. Qiu, X., Zhang, Y., Martin-Rufino, J.D., Weng, C., Hosseinzadeh, S., Yang, D., Pogson, A.N., Hein, M.Y., Hoi Joseph Min, K., Wang, L., et al. (2022). Mapping transcriptomic vector fields of single cells. Cell 185, 690–711.e45.

23. Bravo González-Blas, C., De Winter, S., Hulselmans, G., Hecker, N., Matetovici, I., Christiaens, V., Poovathingal, S., Wouters, J., Aibar, S., and Aerts, S. (2023). SCENIC+: single-cell multiomic inference of enhancers and gene regulatory networks. Nat. Methods 20, 1355–1367.

24. Fleck, J.S., Jansen, S.M.J., Wollny, D., Zenk, F., Seimiya, M., Jain, A., Okamoto, R., Santel, M., He, Z., Camp, J.G., et al. (2023). Inferring and perturbing cell fate regulomes in human brain organoids. Nature 621, 365–372.

25. Kamimoto, K., Stringa, B., Hoffmann, C.M., Jindal, K., Solnica-Krezel, L., and Morris, S.A. (2023). Dissecting cell identity via network inference and in silico gene perturbation. Nature 614, 742–751.

26. Wang, L., Trasanidis, N., Wu, T., Dong, G., Hu, M., Bauer, D.E., and Pinello, L. (2023). Dictys: dynamic gene regulatory network dissects developmental continuum with single-cell multiomics. Nat. Methods 20, 1368–1378.

27. Yuan, Q., and Duren, Z. (2024). Inferring gene regulatory networks from single-cell multiome data using atlas-scale external data. Nat. Biotechnol. 10.1038/s41587-024-02182-7.

28. Gao, M., Qiao, C., and Huang, Y. (2022). UniTVelo: temporally unified RNA velocity reinforces single-cell trajectory inference. Nat. Commun. 13, 6586.

29. Gu, Y., Blaauw, D., and Welch, J.D. (2022). Bayesian inference of RNA velocity from multi-lineage single-cell data. bioRxiv. 10.1101/2022.07.08.499381.

30. Aivazidis, A., Memi, F., Kleshchevnikov, V., Clarke, B., Stegle, O., and Bayraktar, O. (2023). Model-based inference of RNA velocity modules improves cell fate prediction. bioRxiv. 10.1101/2023.08.03.551650.

31. Li, J., Pan, X., Yuan, Y., and Shen, H.-B. (2024). TFvelo: gene regulation inspired RNA velocity estimation. Nat. Commun. 15, 1387.

32. Lambert, S.A., Jolma, A., Campitelli, L.F., Das, P.K., Yin, Y., Albu, M., Chen, X., Taipale, J., Hughes, T.R., and Weirauch, M.T. (2018). The Human Transcription Factors. Cell 172, 650–665.

33. Ma, S., Zhang, B., LaFave, L.M., Earl, A.S., Chiang, Z., Hu, Y., Ding, J., Brack, A., Kartha, V.K., Tay, T., et al. (2020). Chromatin Potential Identified by Shared Single-Cell Profiling of RNA and Chromatin. Cell 183, 1103–1116.e20.

34. Liu, T., Ortiz, J.A., Taing, L., Meyer, C.A., Lee, B., Zhang, Y., Shin, H., Wong, S.S., Ma, J., Lei, Y., et al. (2011). Cistrome: an integrative platform for transcriptional regulation studies. Genome Biol. 12, R83.

35. Zhang, S., Pyne, S., Pietrzak, S., Halberg, S., McCalla, S.G., Siahpirani, A.F., Sridharan, R., and Roy, S. (2023). Inference of cell type-specific gene regulatory networks on cell lineages from single cell omic datasets. Nat. Commun. 14, 3064.

36. Chen, C.-H., Zheng, R., Tokheim, C., Dong, X., Fan, J., Wan, C., Tang, Q., Brown, M., Liu, J.S., Meyer, C.A., et al. (2020). Determinants of transcription factor regulatory range. Nat. Commun. 11, 2472.

37. Lienen, M., and Günnemann, S. (2022). torchode: A Parallel ODE Solver for PyTorch. 10.48550/ARXIV.2210.12375.

38. Kingma, D.P., and Welling, M. (2013). Auto-Encoding Variational Bayes. 10.48550/ARXIV.1312.6114.

39. Mahdessian, D., Cesnik, A.J., Gnann, C., Danielsson, F., Stenström, L., Arif, M., Zhang, C., Le, T., Johansson, F., Schutten, R., et al. (2021). Spatiotemporal dissection of the cell cycle with single-cell proteogenomics. Nature 590, 649–654.

40. Schiebinger, G., Shu, J., Tabaka, M., Cleary, B., Subramanian, V., Solomon, A., Gould, J., Liu, S., Lin, S., Berube, P., et al. (2019). Optimal-Transport Analysis of Single-Cell Gene Expression Identifies Developmental Trajectories in Reprogramming. Cell 176, 928–943.e22.

41. Taylor, S.S., and McKeon, F. (1997). Kinetochore localization of murine Bub1 is required for normal mitotic timing and checkpoint response to spindle damage. Cell 89, 727–735.

42. Bandara, L.R., Lam, E.W., Sørensen, T.S., Zamanian, M., Girling, R., and La Thangue, N.B. (1994). DP-1: a cell cycle-regulated and phosphorylated component of transcription factor DRTF1/E2F which is functionally important for recognition by pRb and the adenovirus E4 orf 6/7 protein. EMBO J. 13, 3104–3114.

43. Nielsen, C.F., Zhang, T., Barisic, M., Kalitsis, P., and Hudson, D.F. (2020). Topoisomerase IIα is essential for maintenance of mitotic chromosome structure. Proc. Natl. Acad. Sci. U. S. A. 117, 12131–12142.

44. Ogata, H., Goto, S., Sato, K., Fujibuchi, W., Bono, H., and Kanehisa, M. (1999). KEGG: Kyoto Encyclopedia of Genes and Genomes. Nucleic Acids Res. 27, 29–34.

45. Zou, Z., Ohta, T., Miura, F., and Oki, S. (2022). ChIP-Atlas 2021 update: a data-mining suite for exploring epigenomic landscapes by fully integrating ChIP-seq, ATAC-seq and Bisulfite-seq data. Nucleic Acids Res. 50, W175–W182.

46. Moerman, T., Aibar Santos, S., Bravo González-Blas, C., Simm, J., Moreau, Y., Aerts, J., and Aerts, S. (2019). GRNBoost2 and Arboreto: efficient and scalable inference of gene regulatory networks. Bioinformatics 35, 2159–2161.

47. Bastidas-Ponce, A., Tritschler, S., Dony, L., Scheibner, K., Tarquis-Medina, M., Salinno, C., Schirge, S., Burtscher, I., Böttcher, A., Theis, F.J., et al. (2019). Comprehensive single cell mRNA profiling reveals a detailed roadmap for pancreatic endocrinogenesis. Development 146. 10.1242/dev.173849.

48. Klein, D., Palla, G., Lange, M., Klein, M., Piran, Z., Gander, M., Meng-Papaxanthos, L., Sterr, M., Bastidas-Ponce, A., Tarquis-Medina, M., et al. (2023). Mapping cells through time and space with moscot. bioRxiv. 10.1101/2023.05.11.540374.

49. Yu, X.-X., Qiu, W.-L., Yang, L., Wang, Y.-C., He, M.-Y., Wang, D., Zhang, Y., Li, L.-C., Zhang, J., Wang, Y., et al. (2021). Sequential progenitor states mark the generation of pancreatic endocrine lineages in mice and humans. Cell Res. 31, 886–903.

50. Bohuslavova, R., Fabriciova, V., Lebrón-Mora, L., Malfatti, J., Smolik, O., Valihrach, L., Benesova, S., Zucha, D., Berkova, Z., Saudek, F., et al. (2023). ISL1 controls pancreatic alpha cell fate and beta cell maturation. Cell Biosci. 13, 53.

51. Baron, M., Veres, A., Wolock, S.L., Faust, A.L., Gaujoux, R., Vetere, A., Ryu, J.H., Wagner, B.K., Shen-Orr, S.S., Klein, A.M., et al. (2016). A Single-Cell Transcriptomic Map of the Human and Mouse Pancreas Reveals Inter- and Intra-cell Population Structure. Cell Syst 3, 346–360.e4.

52. Duvall, E., Benitez, C.M., Tellez, K., Enge, M., Pauerstein, P.T., Li, L., Baek, S., Quake, S.R., Smith, J.P., Sheffield, N.C., et al. (2022). Single-cell transcriptome and accessible chromatin dynamics during endocrine pancreas development. Proc. Natl. Acad. Sci. U. S. A. 119, e2201267119.

53. Heller, R.S., Jenny, M., Collombat, P., Mansouri, A., Tomasetto, C., Madsen, O.D., Mellitzer, G., Gradwohl, G., and Serup, P. (2005). Genetic determinants of pancreatic epsilon-cell development. Dev. Biol. 286, 217–224.

54. Shan, B., and Lee, W.H. (1994). Deregulated expression of E2F-1 induces S-phase entry and leads to apoptosis. Mol Cell Biol 14, 8166–8173.

55. Zhu, W., Giangrande, P.H., and Nevins, J.R. (2004). E2Fs link the control of G1/S and G2/M transcription. EMBO J. 23, 4615–4626.

56. Zhang, Y., Xing, Y., Zhang, L., Mei, Y., Yamamoto, K., Mak, T.W., and You, H. (2012). Regulation of cell cycle progression by forkhead transcription factor FOXO3 through its binding partner DNA replication factor Cdt1. Proc. Natl. Acad. Sci. U. S. A. 109, 5717– 5722.

57. Tien, A.L., Senbanerjee, S., Kulkarni, A., Mudbhary, R., Goudreau, B., Ganesan, S., Sadler, K.C., and Ukomadu, C. (2011). UHRF1 depletion causes a G2/M arrest, activation of DNA damage response and apoptosis. Biochem. J 435, 175–185.

58. Rubin, C.I., and Atweh, G.F. (2004). The role of stathmin in the regulation of the cell cycle. J. Cell. Biochem. 93, 242–250.

59. Lasorella, A., Iavarone, A., and Israel, M.A. (1996). Id2 specifically alters regulation of the cell cycle by tumor suppressor proteins. Mol. Cell. Biol. 16, 2570–2578.

60. Liang, C., and Stillman, B. (1997). Persistent initiation of DNA replication and chromatin-bound MCM proteins during the cell cycle in cdc6 mutants. Genes Dev. 11, 3375–3386.

61. Wu, S.C., and Benavente, C.A. (2018). Chromatin remodeling protein HELLS is upregulated by inactivation of the RB-E2F pathway and is nonessential for osteosarcoma tumorigenesis. Oncotarget 9, 32580–32592.

62. Li, G., Shen, J., Cheng, W., Wang, X., Wang, D., Song, Y., Chen, Y., Li, X., Zhang, M., Ding, Y., et al. (2024). CENPK orchestrates ovarian cancer progression via GOLPH3-Mediated activation of mTOR signaling. Mol. Cell. Endocrinol. 589, 112253.

63. Collombat, P., Hecksher-Sørensen, J., Krull, J., Berger, J., Riedel, D., Herrera, P.L., Serup, P., and Mansouri, A. (2007). Embryonic endocrine pancreas and mature beta cells acquire alpha and PP cell phenotypes upon Arx misexpression. J. Clin. Invest. 117, 961–970.

64. Gao, T., McKenna, B., Li, C., Reichert, M., Nguyen, J., Singh, T., Yang, C., Pannikar, A., Doliba, N., Zhang, T., et al. (2014). Pdx1 maintains *β* cell identity and function by repressing an α cell program. Cell Metab. 19, 259–271.

65. Pan, F.C., Brissova, M., Powers, A.C., Pfaff, S., and Wright, C.V.E. (2015). Inactivating the permanent neonatal diabetes gene Mnx1 switches insulin-producing β-cells to a δ-like fate and reveals a facultative proliferative capacity in aged β-cells. Development 142, 3637– 3648.

66. Zhang, J., McKenna, L.B., Bogue, C.W., and Kaestner, K.H. (2014). The diabetes gene Hhex maintains δ-cell differentiation and islet function. Genes Dev. 28, 829–834.

67. Cota, P., Saber, L., Taskin, D., Jing, C., Bastidas-Ponce, A., Vanheusden, M., Shahryari, A., Sterr, M., Burtscher, I., Bakhti, M., et al. (2023). NEUROD2 function is dispensable for human pancreatic *β* cell specification. Front. Endocrinol. 14, 1286590.

68. Soyer, J., Flasse, L., Raffelsberger, W., Beucher, A., Orvain, C., Peers, B., Ravassard, P., Vermot, J., Voz, M.L., Mellitzer, G., et al. (2010). Rfx6 is an Ngn3-dependent winged helix transcription factor required for pancreatic islet cell development. Development 137, 203– 212.

69. Barile, M., Imaz-Rosshandler, I., Inzani, I., Ghazanfar, S., Nichols, J., Marioni, J.C., Guibentif, C., and Göttgens, B. (2021). Coordinated changes in gene expression kinetics underlie both mouse and human erythroid maturation. Genome Biol 22, 197.

70. Weiler, P., Van den Berge, K., Street, K., and Tiberi, S. (2023). A Guide to Trajectory Inference and RNA Velocity. Methods Mol Biol 2584, 269–292.

71. Puig, R.R., Boddie, P., Khan, A., Castro-Mondragon, J.A., and Mathelier, A. (2021). UniBind: maps of high-confidence direct TF-DNA interactions across nine species. BMC Genomics 22, 482.

72. Iwasaki, H., Mizuno, S.-I., Wells, R.A., Cantor, A.B., Watanabe, S., and Akashi, K. (2003). GATA-1 converts lymphoid and myelomonocytic progenitors into the megakaryocyte/erythrocyte lineages. Immunity 19, 451–462.

73. Chen, H.M., Zhang, P., Voso, M.T., Hohaus, S., Gonzalez, D.A., Glass, C.K., Zhang, D.E., and Tenen, D.G. (1995). Neutrophils and monocytes express high levels of PU.1 (Spi-1) but not Spi-B. Blood 85, 2918–2928.

74. Zhang, P., Zhang, X., Iwama, A., Yu, C., Smith, K.A., Mueller, B.U., Narravula, S., Torbett, B.E., Orkin, S.H., and Tenen, D.G. (2000). PU.1 inhibits GATA-1 function and erythroid differentiation by blocking GATA-1 DNA binding. Blood 96, 2641–2648.

75. Nerlov, C., Querfurth, E., Kulessa, H., and Graf, T. (2000). GATA-1 interacts with the myeloid PU.1 transcription factor and represses PU.1-dependent transcription. Blood 95, 2543–2551.

76. Horak, C.E., Mahajan, M.C., Luscombe, N.M., Gerstein, M., Weissman, S.M., and Snyder, M. (2002). GATA-1 binding sites mapped in the beta-globin locus by using mammalian chIp-chip analysis. Proc. Natl. Acad. Sci. U. S. A. 99, 2924–2929.

77. Aliee, H., Richter, T., Solonin, M., Ibarra, I., Theis, F., and Kilbertus, N. (2022). Sparsity in continuous-depth neural networks. arXiv [cs.LG]. 10.48550/ARXIV.2210.14672.

78. Edens, B.M., Stundl, J., Urrutia, H.A., and Bronner, M.E. (2024). Neural crest origin of sympathetic neurons at the dawn of vertebrates. Nature 629, 121–126.

79. Kim, S., Morgunova, E., Naqvi, S., Goovaerts, S., Bader, M., Koska, M., Popov, A., Luong, C., Pogson, A., Swigut, T., et al. (2024). DNA-guided transcription factor cooperativity shapes face and limb mesenchyme. Cell 187, 692–711 e26.

80. Hu, Z., and Sauka-Spengler, T. (2022). Cellular plasticity in the neural crest and cancer. Curr. Opin. Genet. Dev. 75, 101928.

81. Hagemann-Jensen, M., Ziegenhain, C., Chen, P., Ramsköld, D., Hendriks, G.-J., Larsson, A.J.M., Faridani, O.R., and Sandberg, R. (2020). Single-cell RNA counting at allele and isoform resolution using Smart-seq3. Nat. Biotechnol. 38, 708–714.

82. Hu, Z., Mayes, S., Wang, W., Santos-Pereira, J.M., Theis, F., and Sauka-Spengler, T. (2024). Single-cell multi-omics, spatial transcriptomics and systematic perturbation decode circuitry of neural crest fate decisions. bioRxiv. 10.1101/2024.09.17.613303.

83. Rocha, M., Singh, N., Ahsan, K., Beiriger, A., and Prince, V.E. (2020). Neural crest development: insights from the zebrafish. Dev. Dyn. 249, 88–111.

84. Opdecamp, K., Nakayama, A., Nguyen, M.T., Hodgkinson, C.A., Pavan, W.J., and Arnheiter, H. (1997). Melanocyte development in vivo and in neural crest cell cultures: crucial dependence on the Mitf basic-helix-loop-helix-zipper transcription factor. Development 124, 2377–2386.

85. Aoki, Y., Saint-Germain, N., Gyda, M., Magner-Fink, E., Lee, Y.-H., Credidio, C., and Saint-Jeannet, J.-P. (2003). Sox10 regulates the development of neural crest-derived melanocytes in Xenopus. Dev. Biol. 259, 19–33.

86. Spokony, R.F., Aoki, Y., Saint-Germain, N., Magner-Fink, E., and Saint-Jeannet, J.-P. (2002). The transcription factor Sox9 is required for cranial neural crest development in Xenopus. Development 129, 421–432.

87. Okeke, C., Paulding, D., Riedel, A., Paudel, S., Phelan, C., Teng, C.S., and Barske, L. (2022). Control of cranial ectomesenchyme fate by Nr2f nuclear receptors. Development 149. 10.1242/dev.201133.

88. Soo, K., O’Rourke, M.P., Khoo, P.-L., Steiner, K.A., Wong, N., Behringer, R.R., and Tam, P.P.L. (2002). Twist function is required for the morphogenesis of the cephalic neural tube and the differentiation of the cranial neural crest cells in the mouse embryo. Dev. Biol. 247, 251–270.

89. Wang, C., Kam, R.K.T., Shi, W., Xia, Y., Chen, X., Cao, Y., Sun, J., Du, Y., Lu, G., Chen, Z., et al. (2015). The proto-oncogene transcription factor Ets1 regulates neural crest development through histone deacetylase 1 to mediate output of bone morphogenetic protein signaling. J. Biol. Chem. 290, 21925–21938.

90. Yan, Y.-L., Willoughby, J., Liu, D., Crump, J.G., Wilson, C., Miller, C.T., Singer, A., Kimmel, C., Westerfield, M., and Postlethwait, J.H. (2005). A pair of Sox: distinct and overlapping functions of zebrafish sox9 co-orthologs in craniofacial and pectoral fin development. Development 132, 1069–1083.

91. Das, A., and Crump, J.G. (2012). Bmps and id2a act upstream of Twist1 to restrict ectomesenchyme potential of the cranial neural crest. PLoS Genet. 8, e1002710.

92. Replogle, J.M., Norman, T.M., Xu, A., Hussmann, J.A., Chen, J., Cogan, J.Z., Meer, E.J., Terry, J.M., Riordan, D.P., Srinivas, N., et al. (2020). Combinatorial single-cell CRISPR screens by direct guide RNA capture and targeted sequencing. Nat. Biotechnol. 38, 954– 961.

93. Pelea, O., Mayes, S., Ferry, Q.R.V., Fulga, T.A., and Sauka-Spengler, T. (2024). Specific Modulation of CRISPR Transcriptional Activators through RNA-Sensing Guide RNAs in Mammalian Cells and Zebrafish Embryos. eLife 12. 10.7554/eLife.87722.2.

94. Barske, L., Rataud, P., Behizad, K., Del Rio, L., Cox, S.G., and Crump, J.G. (2018). Essential role of Nr2f nuclear receptors in patterning the vertebrate upper jaw. Dev. Cell 44, 337–347.e5.

95. Saunders, L.M., Mishra, A.K., Aman, A.J., Lewis, V.M., Toomey, M.B., Packer, J.S., Qiu, X., McFaline-Figueroa, J.L., Corbo, J.C., Trapnell, C., et al. (2019). Thyroid hormone regulates distinct paths to maturation in pigment cell lineages. Elife 8. 10.7554/eLife.45181.

96. Howard, A.G. th, Baker, P.A., Ibarra-Garcia-Padilla, R., Moore, J.A., Rivas, L.J., Tallman, J.J., Singleton, E.W., Westheimer, J.L., Corteguera, J.A., and Uribe, R.A. (2021). An atlas of neural crest lineages along the posterior developing zebrafish at single-cell resolution. Elife 10. 10.7554/eLife.60005.

97. Johnson, S.L., Nguyen, A.N., and Lister, J.A. (2011). mitfa is required at multiple stages of melanocyte differentiation but not to establish the melanocyte stem cell. Dev. Biol. 350, 405–413.

98. Tatarakis, D., Cang, Z., Wu, X., Sharma, P.P., Karikomi, M., MacLean, A.L., Nie, Q., and Schilling, T.F. (2021). Single-cell transcriptomic analysis of zebrafish cranial neural crest reveals spatiotemporal regulation of lineage decisions during development. Cell Rep 37, 110140.

99. Petratou, K. (2016). Investigating gene regulatory networks underlying zebrafish pigment cell development.

100. Nair, S., Ameen, M., Sundaram, L., Pampari, A., Schreiber, J., Balsubramani, A., Wang, Y.X., Burns, D., Blau, H.M., Karakikes, I., et al. (2023). Transcription factor stoichiometry, motif affinity and syntax regulate single-cell chromatin dynamics during fibroblast reprogramming to pluripotency. bioRxivorg, 2023.10.04.560808.

101. Choi, H.M.T., Schwarzkopf, M., Fornace, M.E., Acharya, A., Artavanis, G., Stegmaier, J., Cunha, A., and Pierce, N.A. (2018). Third-generation in situ hybridization chain reaction: multiplexed, quantitative, sensitive, versatile, robust. Development 145. 10.1242/dev.165753.

102. Hagedorn, L., Suter, U., and Sommer, L. (1999). P0 and PMP22 mark a multipotent neural crest-derived cell type that displays community effects in response to TGF-beta family factors. Development 126, 3781–3794.

103. Huang, H.-Y., Dai, E.-S., Liu, J.-T., Tu, C.-T., Yang, T.-C., and Tsai, H.-J. (2009). The embryonic expression patterns and the knockdown phenotypes of zebrafish ADP-ribosylation factor-like 6 interacting protein gene. Dev. Dyn. 238, 232–240.

104. Eliason, S., Su, D., Pinho, F., Sun, Z., Zhang, Z., Li, X., Sweat, M., Venugopalan, S.R., He, B., Bustin, M., et al. (2022). HMGN2 represses gene transcription via interaction with transcription factors Lef-1 and Pitx2 during amelogenesis. J. Biol. Chem. 298, 102295.

105. Tian, T., and Burrage, K. (2006). Stochastic models for regulatory networks of the genetic toggle switch. Proc Natl Acad Sci U S A 103, 8372–8377.

106. Bunne, C., Roohani, Y., Rosen, Y., Gupta, A., Zhang, X., Roed, M., Alexandrov, T., AlQuraishi, M., Brennan, P., Burkhardt, D.B., et al. (2024). How to build the virtual cell with artificial intelligence: Priorities and opportunities. arXiv [q-bio.QM]. 10.48550/ARXIV.2409.11654.

107. Madison, B.J., Clark, K.A., Bhachech, N., Hollenhorst, P.C., Graves, B.J., and Currie, S.L. (2018). Electrostatic repulsion causes anticooperative DNA binding between tumor suppressor ETS transcription factors and JUN-FOS at composite DNA sites. J Biol Chem 293, 18624–18635.

108. Soldatov, R., Kaucka, M., Kastriti, M.E., Petersen, J., Chontorotzea, T., Englmaier, L., Akkuratova, N., Yang, Y., Haring, M., Dyachuk, V., et al. (2019). Spatiotemporal structure of cell fate decisions in murine neural crest. Science 364. 10.1126/science.aas9536.

109. Aiello-Couzo, N.M., and Kang, Y. (2020). A bridge between melanoma cell states. Nat Cell Biol 22, 913–914.

110. Kulesa, P.M., Kasemeier-Kulesa, J.C., Teddy, J.M., Margaryan, N.V., Seftor, E.A., Seftor, R.E.B., and Hendrix, M.J.C. (2006). Reprogramming metastatic melanoma cells to assume a neural crest cell-like phenotype in an embryonic microenvironment. Proc. Natl. Acad. Sci. U. S. A. 103, 3752–3757.

111. Chen, Z., King, W.C., Hwang, A., Gerstein, M., and Zhang, J. (2022). : Single-cell transcriptomic deep velocity field learning with neural ordinary differential equations. Sci Adv 8, eabq3745.

112. Singh, R., Wu, A.P., Mudide, A., and Berger, B. (2024). Causal gene regulatory analysis with RNA velocity reveals an interplay between slow and fast transcription factors. Cell Syst 15, 462–474.e5.

113. Lotfollahi, M., Wolf, F.A., and Theis, F.J. (2019). scGen predicts single-cell perturbation responses. Nat Methods 16, 715–721.

114. Kirschenbaum, D., Xie, K., Ingelfinger, F., Katzenelenbogen, Y., Abadie, K., Look, T., Sheban, F., Phan, T.S., Li, B., Zwicky, P., et al. (2024). Time-resolved single-cell transcriptomics defines immune trajectories in glioblastoma. Cell 187, 149–165.e23.

115. Ramanathan, M., Porter, D.F., and Khavari, P.A. (2019). Methods to study RNA-protein interactions. Nat. Methods 16, 225–234.

116. Brannan, K.W., Chaim, I.A., Marina, R.J., Yee, B.A., Kofman, E.R., Lorenz, D.A., Jagannatha, P., Dong, K.D., Madrigal, A.A., Underwood, J.G., et al. (2021). Robust single-cell discovery of RNA targets of RNA-binding proteins and ribosomes. Nat. Methods 18, 507–519.

117. Ozadam, H., Tonn, T., Han, C.M., Segura, A., Hoskins, I., Rao, S., Ghatpande, V., Tran, D., Catoe, D., Salit, M., et al. (2023). Single-cell quantification of ribosome occupancy in early mouse development. Nature 618, 1057–1064.

118. Zeisel, A., Köstler, W.J., Molotski, N., Tsai, J.M., Krauthgamer, R., Jacob-Hirsch, J., Rechavi, G., Soen, Y., Jung, S., Yarden, Y., et al. (2011). Coupled pre-mRNA and mRNA dynamics unveil operational strategies underlying transcriptional responses to stimuli. Mol Syst Biol 7, 529.

119. Yu, L., Wei, Y., Duan, J., Schmitz, D.A., Sakurai, M., Wang, L., Wang, K., Zhao, S., Hon, G.C., and Wu, J. (2021). Blastocyst-like structures generated from human pluripotent stem cells. Nature 591, 620–626.

120. Finlay, C., Jacobsen, J.-H., Nurbekyan, L., and Oberman, A.M. (2020). How to train your neural ODE: the world of Jacobian and kinetic regularization. 10.48550/ARXIV.2002.02798.

121. Loshchilov, I., and Hutter, F. (2017). Decoupled weight decay regularization. 10.48550/ARXIV.1711.05101.

122. Shapiro, A. (2003). Monte Carlo Sampling Methods. In Handbooks in Operations Research and Management Science Handbooks in operations research and management science. (Elsevier), pp. 353–425.

123. Pedregosa, F., Varoquaux, G., Gramfort, A., Michel, V., Thirion, B., Grisel, O., Blondel, M., Müller, A., Nothman, J., Louppe, G., et al. (2012). Scikit-learn: Machine Learning in Python.

124. Blei, D.M., Kucukelbir, A., and McAuliffe, J.D. (2017). Variational inference: A review for statisticians. J. Am. Stat. Assoc. 112, 859–877.

125. Merryweather-Clarke, A.T., Atzberger, A., Soneji, S., Gray, N., Clark, K., Waugh, C., McGowan, S.J., Taylor, S., Nandi, A.K., Wood, W.G., et al. (2011). Global gene expression analysis of human erythroid progenitors. Blood 117, e96–e108.

126. Hu, C., Li, T., Xu, Y., Zhang, X., Li, F., Bai, J., Chen, J., Jiang, W., Yang, K., Ou, Q., et al. (2023). CellMarker 2.0: an updated database of manually curated cell markers in human/mouse and web tools based on scRNA-seq data. Nucleic Acids Res. 51, D870– D876.

127. Cannoodt, R., Saelens, W., Deconinck, L., and Saeys, Y. (2021). Spearheading future omics analyses using dyngen, a multi-modal simulator of single cells. Nat. Commun. 12, 3942.

128. Wolf, F.A., Angerer, P., and Theis, F.J. (2018). SCANPY: large-scale single-cell gene expression data analysis. Genome Biol. 19, 15.

129. Shen, W.-K., Chen, S.-Y., Gan, Z.-Q., Zhang, Y.-Z., Yue, T., Chen, M.-M., Xue, Y., Hu, H., and Guo, A.-Y. (2023). AnimalTFDB 4.0: a comprehensive animal transcription factor database updated with variation and expression annotations. Nucleic Acids Res. 51, D39– D45.

130. Reuter, B., Fackeldey, K., and Weber, M. (2019). Generalized Markov modeling of nonreversible molecular kinetics. J. Chem. Phys. 150, 174103.

131. Bocci, F., Zhou, P., and Nie, Q. (2022). spliceJAC: transition genes and state-specific gene regulation from single-cell transcriptome data. Mol. Syst. Biol. 18, e11176.

132. Pratapa, A., Jalihal, A.P., Law, J.N., Bharadwaj, A., and Murali, T.M. (2020). Benchmarking algorithms for gene regulatory network inference from single-cell transcriptomic data. Nat. Methods 17, 147–154.

133. Oki, S., Ohta, T., Shioi, G., Hatanaka, H., Ogasawara, O., Okuda, Y., Kawaji, H., Nakaki, R., Sese, J., and Meno, C. (2018). ChIP-Atlas: a data-mining suite powered by full integration of public ChIP-seq data. EMBO Rep. 19. 10.15252/embr.201846255.

134. Gaspar, J.M. (2018). Improved peak-calling with MACS2. bioRxiv. 10.1101/496521.

135. Zhang, Y., Liu, T., Meyer, C.A., Eeckhoute, J., Johnson, D.S., Bernstein, B.E., Nusbaum, C., Myers, R.M., Brown, M., Li, W., et al. (2008). Model-based analysis of ChIP-Seq (MACS). Genome Biol. 9, R137.

136. Schep, A.N., Wu, B., Buenrostro, J.D., and Greenleaf, W.J. (2017). chromVAR: inferring transcription-factor-associated accessibility from single-cell epigenomic data. Nat. Methods 14, 975–978.

137. Chen, T., and Guestrin, C. (2016). XGBoost. In Proceedings of the 22nd ACM SIGKDD International Conference on Knowledge Discovery and Data Mining (ACM). 10.1145/2939672.2939785.

138. Stuart, T., Srivastava, A., Madad, S., Lareau, C.A., and Satija, R. (2021). Single-cell chromatin state analysis with Signac. Nat. Methods 18, 1333–1341.

139. Westerfield, M. (2000). The Zebrafish Book: A Guide for the Laboratory Use of Zebrafish (Danio Rerio) (University of Oregon Press).

140. Kimmel, C.B., Ballard, W.W., Kimmel, S.R., Ullmann, B., and Schilling, T.F. (2005). Stages of embryonic development of the zebrafish. Dev. Dyn. 203, 253–310.

141. Hochgreb-Hagele, T., and Bronner, M.E. (2013). A novel FoxD3 gene trap line reveals neural crest precursor movement and a role for FoxD3 in their specification. Dev. Biol. 374, 1–11.

142. Parekh, S., Ziegenhain, C., Vieth, B., Enard, W., and Hellmann, I. (2018). zUMIs - A fast and flexible pipeline to process RNA sequencing data with UMIs. Gigascience 7. 10.1093/gigascience/giy059.

143. Dobin, A., Davis, C.A., Schlesinger, F., Drenkow, J., Zaleski, C., Jha, S., Batut, P., Chaisson, M., and Gingeras, T.R. (2012). STAR: ultrafast universal RNA-seq aligner. Bioinformatics 29, 15–21.

144. Hao, Y., Hao, S., Andersen-Nissen, E., Mauck, W.M., Zheng, S., Butler, A., Lee, M.J., Wilk, A.J., Darby, C., Zager, M., et al. (2021). Integrated analysis of multimodal single-cell data. Cell 184, 3573–3587.e29.

145. Moon, K.R., van Dijk, D., Wang, Z., Gigante, S., Burkhardt, D.B., Chen, W.S., Yim, K., van den Elzen, A., Hirn, M.J., Coifman, R.R., et al. (2019). Visualizing structure and transitions in high-dimensional biological data. Nat. Biotechnol. 37, 1482–1492.

146. Burkhardt, D.B., Stanley, J.S., 3rd, Tong, A., Perdigoto, A.L., Gigante, S.A., Herold, K.C., Wolf, G., Giraldez, A.J., van Dijk, D., and Krishnaswamy, S. (2021). Quantifying the effect of experimental perturbations at single-cell resolution. Nat. Biotechnol. 39, 619–629.

147. Li, D., Hsu, S., Purushotham, D., Sears, R.L., and Wang, T. (2019). WashU Epigenome Browser update 2019. Nucleic Acids Res. 47, W158–W165.

