## Supplementary notes for "RegVelo: gene-regulatory-informed dynamics of single cells"

#### Supplementary Note 1

Single-cell transcriptomic data is noisy, sparse, and high-dimensional, making velocity inference challenging. Additionally, single-cell data is traditionally snapshot data, *i.e.*, experimental assays measure cells a single time instead of tracking them over a time span. As such, assessing if the estimated state change is correct purely based on the gene expression data is impossible. To benchmark the performance of velocity inference nonetheless, we simulated single-cell gene expression based on a gene regulatory network (GRN) informed stochastic differential equation (SDE) and the simulation framework dyngen[1].

We first compared model performance on a simulated GRN including six genes. This network includes three toggle switches in a symmetrical hierarchical structure representing a previously used GRN for modeling cell fate decision-making during differentiation[2,3] (Supplementary Fig. 1a). Using an SDE solver[4], we defined a recursive system of equations to simulate unspliced and spliced RNA abundances, where each regulator uses a nonlinear Hill-type function to activate or suppress target gene expression, with logic OR gates employed (Methods). We ran 100 simulations with reaction rates sampled from a log-normal multivariate distribution and time events following a Poisson distribution (Supplementary Note 2)

RegVelo regularizes the Jacobian matrix of the dynamical with the L1 norm, denoted by J-L1 regularization, to ensure smooth trajectories rather than directly regularizing the weight matrix W of the transcription predictor neural network (w-L1 regularization; Figure 1a). To test the contribution of this component to RegVelo's performance, we benchmarked latent time and GRN inference of the J-L1 and w-L1 regularization models. We evaluated latent time inference using Spearman's correlation between estimates and ground truth and assessed GRN inference by the area under the receiver operating characteristic (AUROC; Methods): Overall, constraining the Jacobian led to significantly improved GRN identification with stable latent time inference compared to w-L1 regularization (two-sided Welch's *t*-test, $P < 0.001$; Supplementary Fig. 1b; Methods).

We further benchmarked RegVelo against veloVI[5] and scVelo's[6] *EM* model; all other models for inferring dynamics inference failed to run successfully on the simulated data (Methods). The velocities and latent time inferred by RegVelo demonstrated a significantly higher correlation with ground truth compared to veloVI and scVelo (two-sided Welch's *t*-test, $P < 0.001$; Supplementary Fig. 1c). In addition, we benchmarked RegVelo against other methods for GRN inference and assessed model performance by the AUROC; RegVelo again significantly performed better than the remaining methods in predicting ground truth gene regulations (two-sided Welch's *t*-test, $P < 0.001$; Supplementary Fig. 1d).

This toy simulation model offered valuable insights as it is tractable and interpretable but also simplistic. To generate a more realistic system, we employed the dyngen[1] simulation framework that improves GRN simulation by incorporating protein modality simulation, offering a more complete representation of gene expression dynamics. We simulated 50 new scRNA-seq datasets with known ground truth cell-specific GRNs and RNA velocity (Methods).

Among all velocity inference models, RegVelo achieved the highest Pearson correlation between estimated and ground truth velocities (Supplementary Fig. 2a). Like scVelo and veloVI, RegVelo also estimates gene-specific latent time that we compared against ground truth and found again that RegVelo consistently outperformed scVelo and veloVI (Supplementary Fig. 2b). We further benchmarked RegVelo against five other methods for GRN inference, where RegVelo yielded consistent results and significantly outperformed all other methods (two-sided Welch's *t*-test, $P < 0.05$; Supplementary Fig. 2c).

The possibility to incorporate prior knowledge of gene regulation to infer cell dynamics is one advantage of RegVelo. However, prior GRNs are often incomplete or even incorrect. To assess RegVelo's robustness to a prior GRN and, thus, potential errors in gene regulation, we randomly removed varying fractions of edges of the true GRN of our simulated datasets. We then used the newly generated GRN as prior knowledge for RegVelo to estimate velocities, latent time, and the GRN; we repeated this process five times for each corruption rate to assess statistical significance. Notably, RegVelo consistently demonstrated strong identifiability (Spearman correlation > 0.9) of the GRN weights, velocities, and latent times, showcasing its robustness (Supplementary Fig. 2d). Furthermore, as corruption rates decreased, RegVelo consistently improved its GRN inference (Supplementary Fig. 2e); the velocity and latent time inference performance are also stable under different corruption rates (Supplementary Fig. 2f, g).

### Supplementary Note 2

We provide details on the implementation and data processing of our analysis of simulated data as a supplement to our main Methods section.

We simulated a toy GRN containing six genes and three toggle switch network motifs (Supplementary Fig. 1a): following a previous simulation method based on Hill functions[3], we used 12 stochastic differential equations (SDEs) to describe the regulatory OR-gate dynamics among the six genes A, B, C, D, E, and F, where toggle switches positively regulate each other.

$$
\begin{aligned}
\mathrm{d}u_{\mathrm{A}} &= \left[\frac{\kappa_{-}^{h_{-}}}{s_{\mathrm{B}}(t)^{h_{-}}+\kappa_{-}^{h_{-}}} - \beta_{\mathrm{A}} u_{\mathrm{A}}(t)\right]\mathrm{d}t + \sigma_s \mathrm{d}W_t \\
\mathrm{d}s_{\mathrm{A}} &= \left[\beta_{\mathrm{A}} u_{\mathrm{A}}(t) - \gamma_{\mathrm{A}} s_{\mathrm{A}}(t)\right]\mathrm{d}t + \sigma_s \mathrm{d}W_t \\
\mathrm{d}u_{\mathrm{B}} &= \left[\frac{\kappa_{-}^{h_{-}}}{s_{\mathrm{A}}(t)^{h_{-}}+\kappa_{-}^{h_{-}}} - \beta_{\mathrm{B}} u_{\mathrm{B}}(t)\right]\mathrm{d}t + \sigma_s \mathrm{d}W_t \\
\mathrm{d}s_{\mathrm{B}} &= \left[\beta_{\mathrm{B}} u_{\mathrm{B}}(t) - \gamma_{\mathrm{B}} s_{\mathrm{B}}(t)\right]\mathrm{d}t + \sigma_s \mathrm{d}W_t \\
\mathrm{d}u_{\mathrm{C}} &= \left[\alpha_{max}\left(\frac{s_{\mathrm{A}}(t)^{h_{+}}}{s_{\mathrm{A}}(t)^{h_{+}}+\kappa_{+}^{h_{-}}} + \frac{\kappa_{-}^{h_{-}}}{s_{\mathrm{D}}(t)^{h_{-}}+\kappa_{-}^{h_{-}}}\right) - \beta_{\mathrm{C}} u_{\mathrm{C}}(t)\right]\mathrm{d}t + \sigma_s \mathrm{d}W_t \\
\mathrm{d}s_{\mathrm{C}} &= \left[\beta_{\mathrm{C}} u_{\mathrm{C}}(t) - \gamma_{\mathrm{C}} s_{\mathrm{C}}(t)\right]\mathrm{d}t + \sigma_s \mathrm{d}W_t \\
\mathrm{d}u_{\mathrm{D}} &= \left[\alpha_{max}\left(\frac{s_{\mathrm{A}}(t)^{h_{+}}}{s_{\mathrm{A}}(t)^{h_{+}}+\kappa_{+}^{h_{-}}} + \frac{\kappa_{-}^{h_{-}}}{s_{\mathrm{C}}(t)^{h_{-}}+\kappa_{-}^{h_{-}}}\right) - \beta_{\mathrm{D}} u_{\mathrm{D}}(t)\right]\mathrm{d}t + \sigma_s \mathrm{d}W_t \mathrm{d}s_{\mathrm{D}} \\
&= \left[\beta_{\mathrm{D}} u_{\mathrm{D}}(t) - \gamma_{\mathrm{D}} s_{\mathrm{D}}(t)\right]\mathrm{d}t + \sigma_s \mathrm{d}W_t \\
\mathrm{d}u_{\mathrm{E}} &= \left[\alpha_{max}\left(\frac{s_{\mathrm{B}}(t)^{h_{+}}}{s_{\mathrm{B}}(t)^{h_{+}}+\kappa_{+}^{h_{-}}} + \frac{\kappa_{-}^{h_{-}}}{s_{\mathrm{F}}(t)^{h_{-}}+\kappa_{-}^{h_{-}}}\right) - \beta_{\mathrm{E}} u_{\mathrm{E}}(t)\right]\mathrm{d}t + \sigma_s \mathrm{d}W_t \mathrm{d}s_{\mathrm{E}} \\
&= \left[\beta_{\mathrm{E}} u_{\mathrm{E}}(t) - \gamma_{\mathrm{E}} s_{\mathrm{E}}(t)\right]\mathrm{d}t + \sigma_s \mathrm{d}W_t \\
\mathrm{d}u_{\mathrm{F}} &= \left[\alpha_{max}\left(\frac{s_{\mathrm{B}}(t)^{h_{+}}}{s_{\mathrm{B}}(t)^{h_{+}}+\kappa_{+}^{h_{-}}} + \frac{\kappa_{-}^{h_{-}}}{s_{\mathrm{E}}(t)^{h_{-}}+\kappa_{-}^{h_{-}}}\right) - \beta_{\mathrm{F}} u_{\mathrm{F}}(t)\right]\mathrm{d}t + \sigma_s \mathrm{d}W_t \\
\mathrm{d}s_{\mathrm{F}} &= \left[\beta_{\mathrm{F}} u_{\mathrm{F}}(t) - \gamma_{\mathrm{F}} s_{\mathrm{F}}(t)\right]\mathrm{d}t + \sigma_s \mathrm{d}W_t,
\end{aligned}
$$

with $\mathrm{d}W_t \sim \mathcal{N}(0,1)$ independently and identically distributed, and $\sigma_s$ denoting the system noise level; based on a previously suggested noise level setting[3], we set $\sigma_s = 1$ for simulation. $\kappa$ represents the dissociation constant set to $\kappa_+ = 4$ for activation and $\kappa_- = 2$ for inhibition. For hill coefficients $h$, we used $h_+ = 2$ and $h_- = 2$. These parameters result in a system that generates highly variable gene expression dynamics across all six genes, enabling us to study the effects of gene regulation.

We then simulated a time vector with 1500 Poisson-distributed time steps with parameter $\lambda = 1$ as suggested by scVelo[6]. To align with scVelo's parameter simulation strategy, we sampled kinetic parameters $\theta = (\alpha_{\max}, \beta, \gamma)$ from a log-normal distribution where $\log(\theta) \sim \mathcal{N}(\mu, \Sigma)$. We set $\mu = (5, 0.5, 0.125)$ and defined a symmetric covariance matrix

$$\Sigma = \begin{bmatrix} 0.4^2 & 0.4^2 \cdot 0.2 & 0.4^2 \cdot 0.2 \\ 0.4^2 \cdot 0.2 & 0.4^2 & 0.4^2 \cdot 0.8 \\ 0.4^2 \cdot 0.2 & 0.4^2 \cdot 0.8 & 0.4^2 \end{bmatrix}$$

We simulated the dynamics with the SDE solver *torchsde*[7] with the Euler-Maruyama method[8] and sampled parameters and time values 100 times to generate 100 datasets under different conditions.

#### Analysis of SDE simulation datasets

We implemented two distinct models to compare the Jacobian (J-L1) to the weight matrix (w-L1) regularization, each incorporating a unique L1 loss function: for J-L1, we defined the loss function $L$ as

$$L = -\text{ELBO} + \lambda L_{\text{Jacobian}},$$

with

$$L_{\text{Jacobian}} = \frac{1}{N} \sum_{n=1}^{N} \left\| \text{diag}\left(\text{sigmoid} \circ (W s_n + b)\right) W \right\|_1,$$

Where the operator $\circ$ applies a scalar function component-wise, and for w-L1

$$L = -\text{ELBO} + \lambda \left| W \right|,$$

where $|W|$ denotes the L1 norm of the weight matrix

We tested each model with three distinct regularization parameter settings $\lambda \in \{0, 0.5, 1\}$ to evaluate the models' performance across different levels of regularization strength. We ran both models on each dataset and calculated the Spearman correlation between estimated latent time and ground truth, and AUROC to evaluate GRN prediction accuracy accordingly.

#### Dyngen simulation

Dyngen[1] allowed us to simulate gene expression dynamics in different ways with high-dimensional GRN and consider the effect of translation processes. To use dyngen for benchmarking, we followed the tool's tutorial with default settings to perform a single-cell dataset simulation. We simulated single-cell datasets with a bifurcating trajectory by setting the backbone through `backbone <- backbone_bifurcating()`; we considered the GRN and RNA velocity simultaneously by setting `compute_cellwise_grn=TRUE` and `compute_rna_velocity=TRUE`.

Following the guidelines provided in the dyngen example tutorial[1], we executed the simulation with the two GRN parameter settings 100 genes, 90 targets, 65 TFs, and 300 cells and 200 genes, 100 targets, 100 TFs, and 1000 cells. This allowed us to benchmark methods on different numbers of genes and cells, providing a comprehensive evaluation. For each parameter set, we simulated 25 distinct datasets to account for simulation variability.

### Supplementary Figures

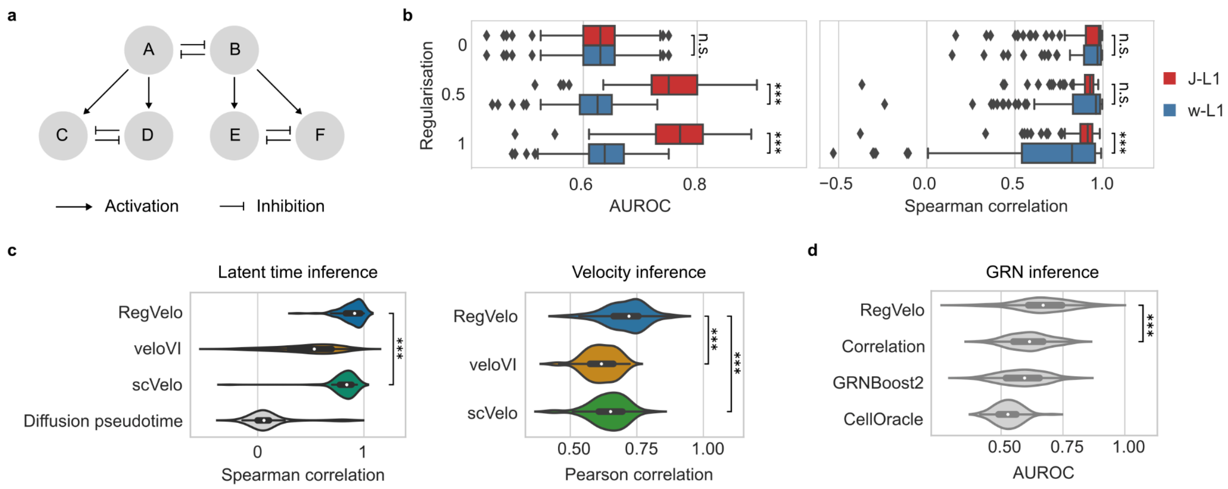


**Suppl. Fig. 1 | Benchmarking RegVelo on a simulated dataset based on a predefined GRN.**

**a.** Graphical representation of the GRN used for data simulation; each letter identifies one gene. **b.** Benchmark of GRN (left) and latent time inference (right) on simulated data using two regularization approaches and different regularization weights (Methods). Box plots indicate the median (center line), interquartile range (hinges), and 1.5x interquartile range (whiskers) ($N = 100$ simulated repeats). **c**. Comparison of inferred latent times (left), velocities (middle) and GRN (right) with ground truth using RegVelo, veloVI, scVelo (*EM* model), or diffusion pseudotime; ($N = 100$ each; left: one-sided Welch's *t*-test, $P = 5.41 \times 10^{-5}$; middle: one-sided Welch's *t*-test, RegVelo vs. veloVI $P = 7.13 \times 10^{-8}$, RegVelo vs. scVelo $P = 3.31 \times 10^{-9}$; right: $P = 1.91 \times 10^{-5}$).

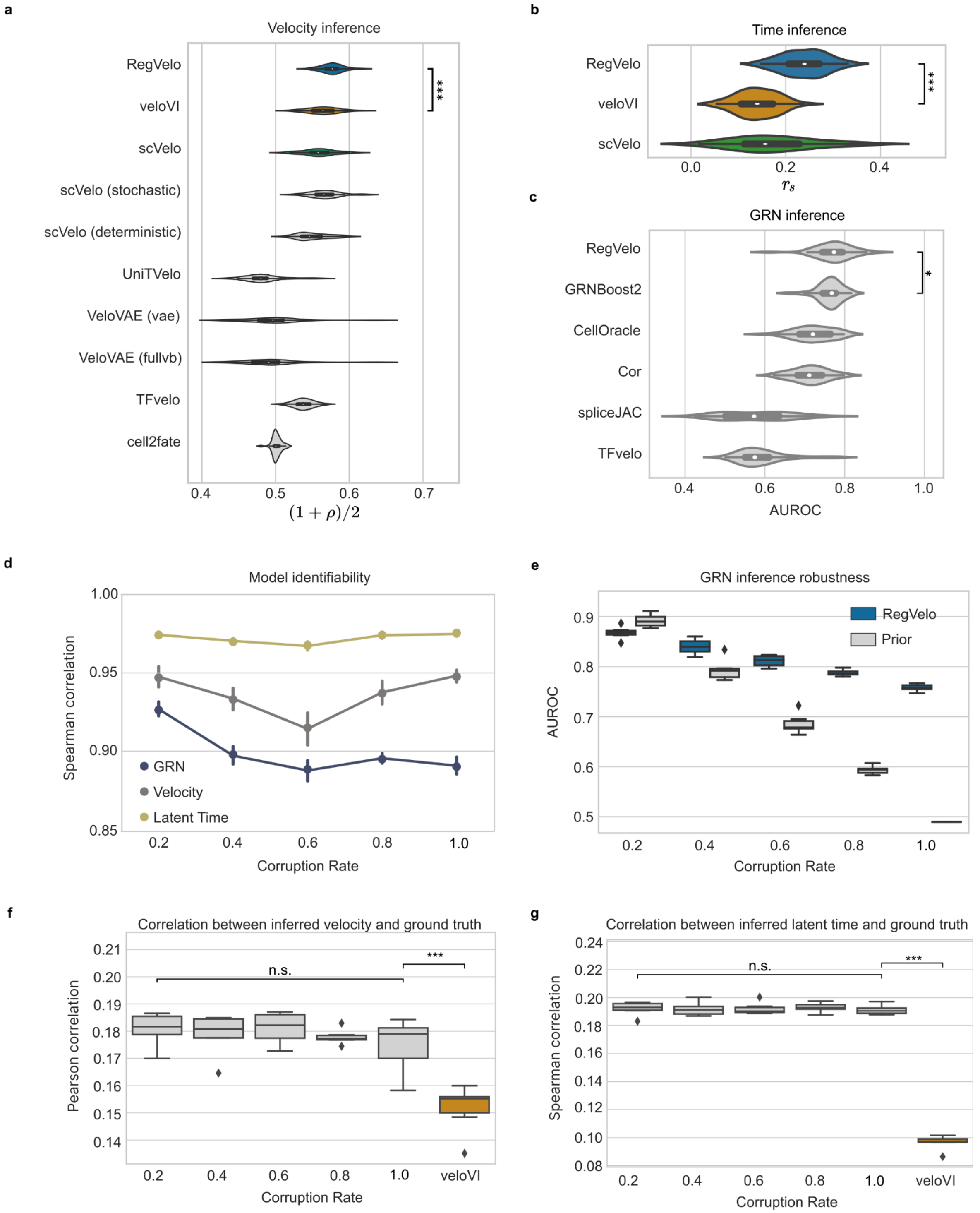


**Suppl. Fig. 2 | Benchmarking RegVelo on dyngen simulations.**
**a-c.** Comparison of RegVelo's and competing approaches' estimated velocities (**a.**), GRN (**b.**), and latent time (**c.**) against ground truth. Box plots indicate the median (center line) and interquartile range (hinges) ( $N = 50$ each; one-sided Welch's t-test, **a.**:

$P = 8.45 \times 10^{-12}$; **b.**: $P = 8.99 \times 10^{-5}$; **c.**: $P < 10 \times 10^{-20}$). **d**. Pairwise correlation of GRN (yellow), velocity (brown), and latent time (purple) estimates across different runs of RegVelo with altered prior GRNs (Methods). For each prior GRN corruption rate, we generated five corrupted prior GRNs and showed the mean performance with the corresponding 95% confidence interval. **e.** Comparison of GRN prediction using corrupted prior GRNs (prior, grey) and RegVelo's estimated GRN (RegVelo, blue). Box plots indicate the median (center line), interquartile range (hinges), and 1.5x interquartile range (whiskers) ($N = 5$ runs). **f-g.** Pearson correlation between ground truth and estimated velocities (**f.**) and latent time (**g.**) on simulated data using RegVelo (grey) under different prior GRN corruption rates and veloVI (orange). Box plots indicate the median (center line), interquartile range (hinges), and whiskers at 1.5x interquartile range ($N = 5$ runs each; one-sided Welch's t-test, **f.**: $P = 1.21 \times 10^{-6}$; **g.**: $P < 10 \times 10^{-11}$; n.s.: not significant).

### References


1. Cannoodt, R., Saelens, W., Deconinck, L., and Saeys, Y. (2021). Spearheading future omics analyses using dyngen, a multi-modal simulator of single cells. Nat Commun *12*, 3942.

2. Haghverdi, L., Büttner, M., Wolf, F.A., Buettner, F., and Theis, F.J. (2016). Diffusion pseudotime robustly reconstructs lineage branching. Nat Methods *13*, 845–848.

3. Ocone, A., Haghverdi, L., Mueller, N.S., and Theis, F.J. (2015). Reconstructing gene regulatory dynamics from high-dimensional single-cell snapshot data. Bioinformatics *31*, i89–i96.

4. Li, X., Wong, T.-K.L., Chen, R.T.Q., and Duvenaud, D. (2020). Scalable Gradients for Stochastic Differential Equations.

5. Gayoso, A., Weiler, P., Lotfollahi, M., Klein, D., Hong, J., Streets, A., Theis, F.J., and Yosef, N. (2024). Deep generative modeling of transcriptional dynamics for RNA velocity analysis in single cells. Nat Methods *21*, 50–59.

6. Bergen, V., Lange, M., Peidli, S., Wolf, F.A., and Theis, F.J. (2020). Generalizing RNA velocity to transient cell states through dynamical modeling. Nat. Biotechnol. *38*, 1408–1414.

7. Li, X., Wong, T.-K.L., Chen, R.T.Q., and Duvenaud, D. (2020). Scalable gradients for stochastic differential equations. https://doi.org/10.48550/ARXIV.2001.01328.

8. Kloeden, P.E., and Platen, E. (2011). Numerical Solution of Stochastic Differential Equations (Springer Science & Business Media).
