## Supplementary tables for "RegVelo: gene-regulatory-informed dynamics of single cells"

#### Supplementary Table 1

##### Erythroid lineage drivers

| Gene | Function | Reference |
| --- | --- | --- |
| GATA1 | GATA1 plays a pivotal role by activating erythroid-specific genes and repressing non-erythroid lineage genes. | Ferreira R, Ohneda K, Yamamoto M, Philipsen S: GATA1 function, a paradigm for transcription factors in hematopoiesis. Mol Cell Biol 2005, 25:1215-1227. |
| GFI1B | GFI1B supports erythroid differentiation by repressing genes that promote alternative lineages and maintaining the quiescence of progenitor cells. | Vassen L, Beauchemin H, Lemsaddek W, Krongold J, Trudel M, Moroy T: Growth factor independence 1b (gfi1b) is important for the maturation of erythroid cells and the regulation of embryonic globin expression. PLoS One 2014, 9:e96636. |
| NFIA | NFIA belongs to the NF1 family of transcription factors which is required for erythroid differentiation. | Velten, Lars, Simon F. Haas, Simon Raffel, Sandra Blaszkiewicz, Saiful Islam, Bianca P. Hennig, Christoph Hirche et al. "Human haematopoietic stem cell lineage commitment is a continuous process." Nature cell biology 19, no. 4 (2017): 271-281. |
| LMO2 | LMO2 contributes to the expression of GATA-1 target genes in a context-dependent manner, through modulating the assembly of the components of GATA-SCL/TAL1 complex at endogeneous loci. | Inoue, Ai, Tohru Fujiwara, Yoko Okitsu, Yuna Katsuoka, Noriko Fukuhara, Yasushi Onishi, Kenichi Ishizawa, and Hideo Harigae. "Elucidation of the role of LMO2 in human erythroid cells." Experimental Hematology 41, no. 12 (2013): 1062-1076. |

| | | |
|---|---|---|
| TAL1 | is essential for the establishment of the erythroid lineage by regulating the expression of genes involved in erythroid progenitor proliferation and survival. | Kassouf MT, Hughes JR, Taylor S, McGowan SJ, Soneji S, Green AL, Vyas P, Porcher C: Genome-wide identification of TAL1's functional targets: insights into its mechanisms of action in primary erythroid cells. Genome Res 2010, 20:1064-1083. |

### Supplementary Table 2

#### Monocyte lineage drivers

| Gene | Function | Reference |
|---|---|---|
| SPI1 | The master regulator of myeloid lineage differentiation. | Smith LT, Hohaus S, Gonzalez DA, Dziennis SE, Tenen DG: PU. 1 (Spi-1) and C/EBP alpha regulate the granulocyte colony-stimulating factor receptor promoter in myeloid cells. 1996. |
| TCF4 | TCF4 is a transcription factor in the E-protein family, which is known to influence hematopoiesis (blood cell formation). Highly expressed in Monocyte/dendritic cell progenitors. | Velten, Lars, Simon F. Haas, Simon Raffel, Sandra Blaszkiewicz, Saiful Islam, Bianca P. Hennig, Christoph Hirche et al. "Human haematopoietic stem cell lineage commitment is a continuous process." Nature cell biology 19, no. 4 (2017): 271-281. |
| STAT6 | Monocyte differentiation is driven by IL-4 – and acts through STAT6. | de Juan, Alba, Darawan Tabtim-On, Alice Coillard, Burkhard Becher, Christel Goudot, and Elodie Segura. "The aryl hydrocarbon receptor shapes monocyte transcriptional responses to interleukin-4 by prolonging STAT6 binding to promoters." Science Signaling 17, no. 858 |

| | | |
|---|---|---|
| | | (2024): eadn6324. |
| MEF2C | MEF2C promotes precursor cell development towards monocyte fate. | Pon, Julia R., and Marco A. Marra. "MEF2 transcription factors: developmental regulators and emerging cancer genes." Oncotarget 7, no. 3 (2015): 2297. |
